# Single-nucleoid imaging in whole cells defines the dynamics of the mtDNA life cycle

**DOI:** 10.64898/2026.06.01.725017

**Authors:** Dane M. Wolf, Endri Mjeku, Mayuko Segawa, Alva M. Casey, Nejma Belaadi, Robin J.M. Franklin, Michael P. Murphy, Julien Prudent, Nick S. Jones, Patrick F. Chinnery

## Abstract

The mitochondrial genome (mtDNA) is essential for oxidative phosphorylation, and mammalian cells typically contain hundreds or thousands of copies. Although specific aspects of mtDNA replication and degradation have been explored, a complete account of the mtDNA life cycle has remained elusive, particularly for nondividing cells. Using super-resolution and 4D lattice light-sheet imaging, we quantified the full mtDNA life cycle in quiescent primary human cells. We show that cells maintain steady state by replicating and degrading 1.5 ± 0.2% of total mtDNA content each hour, a remarkably rapid flux. Younger mtDNA molecules are closer to the nucleus, spared from degradation, and serve as foci for further replication events. Non-proliferative cells regulate nucleoid density within the mitochondrial network, and the mitochondrial membrane potential sustains mtDNA copy number. These findings provide a foundational understanding of the dynamics underlying mtDNA homeostasis and a mechanism explaining how mutations accumulate in aging and disease.

## Main

Mitochondria are essential for producing adenosine triphosphate (ATP) through oxidative phosphorylation (OXPHOS)^1,2^. Maintaining a functional respiratory chain requires the coordination of the cell’s dual genome architecture, compartmentalized in the nucleus and mitochondria^3^. Mitochondrial DNA (mtDNA) encodes 13 core subunits of the electron transport chain (ETC) and ATP synthase, a subset of 37 genes retained by the mitochondrial genome^4^, representing a minute, though pivotal, fraction of the more than 1,100 mitochondrial genes^5^ exported to the nucleus early in eukaryotic evolution^6,7^. Maintenance of the mitochondrial genome is therefore crucial for ensuring OXPHOS functionality across the lifespan of the organism^8^.

In contrast to nuclear DNA (nDNA), the mitochondrial genome is circular^9^ and compacted into nucleoprotein structures called nucleoids^10^, each harboring around 1.4 mtDNA molecules^11^. Individual cells typically contain hundreds or thousands of nucleoids, and their maintenance and distribution throughout the mitochondrial network depend on mitochondrial dynamics^12–15^. Though it is well known that mtDNA maintenance reflects “relaxed replication,” where new copies are made and existing ones are degraded continuously^16,17^, it is important to note that these dynamics are poorly understood in non-proliferative cells, despite the essential role of postmitotic cells in both normal physiology as well as aging and disease^18^. Indeed, the control mechanisms in non-proliferative cells must be distinct from those in proliferative cells, because the maintenance of mtDNA copy number (CN) in non-proliferative cells is not linked to the cell cycle, which upregulates mtDNA synthesis in preparation for cytokinesis^19^, ensuring sufficient CN in daughter cells. Notwithstanding this fundamentally different mode of mtDNA regulation, the specific kinetics of mtDNA replication in quiescent or postmitotic cells have yet to be elucidated.

Since most mtDNA mutations stem from replication errors by the mitochondrial Polymerase γ (Polγ), which is 10-to 100-fold more error-prone than nDNA polymerases^20–22^, the replication rate constitutes a key determinant of the speed at which mutant mitochondrial genomes accumulate, driving aging and disease^23,24^. Bulk estimates of mtDNA turnover have yielded heterogeneous results and are derived from sampling multiple cell types—mixing proliferative and non-proliferative lineages^25^—whereas more refined single-cell studies have considered only proliferative cells^26^, necessarily representing out-of-equilibrium conditions due to the ongoing influence of the cell cycle. Evidence also exists for distinct mtDNA subpopulations, based on differing levels of compaction by mitochondrial transcription factor A (TFAM)^26–29^, as well as subcellular^30,31^ and suborganella^32^ spatial distributions. Nevertheless, a unified understanding of mtDNA dynamics, unconfounded by the cell cycle and relevant to aging postmitotic cells, is lacking. Consequently, the mutational dynamics of the mitochondrial genome, which drive aging and disease, also remain ill-defined.

Here, we overcome these limitations by combining cutting-edge 3D super-resolution and 4D lattice light-sheet imaging, pulse-chase and mtDNA-depletion assays, along with in-depth mathematical modelling of whole individual quiescent primary human cells. Altogether, we provide a comprehensive model with quantitative parameter estimates, including replication and degradation rates, subpopulation interchange frequencies, and relative subpopulation sizes—definitively mapping the spatiotemporal dynamics of the mtDNA life cycle.

## Results

Long-lived, terminally differentiated cells populating most solid organs accumulate mtDNA mutations, which contribute to functional decline and age-associated disease in humans^23^. To investigate the kinetics and regulatory mechanisms of mtDNA in a non-proliferative context, we arrested the cell cycle in primary adult human fibroblasts (Extended Data Fig. 1a–c). We next leveraged the Zeiss Elyra 7 with 3D lattice structured illumination microscopy (SIM) and SIM^2^, which has a lateral resolution down to ~60 nm—comparable to that of stimulated emission depletion (STED), with a lateral resolution of ~50 nm—to visualize the total nucleoid population, including newly replicated mtDNA molecules labelled with 5-ethynyl-2’-deoxyuridine (EdU), within their respective cellular and mitochondrial network volumes (Fig. 1a–c).

**Figure 1:**
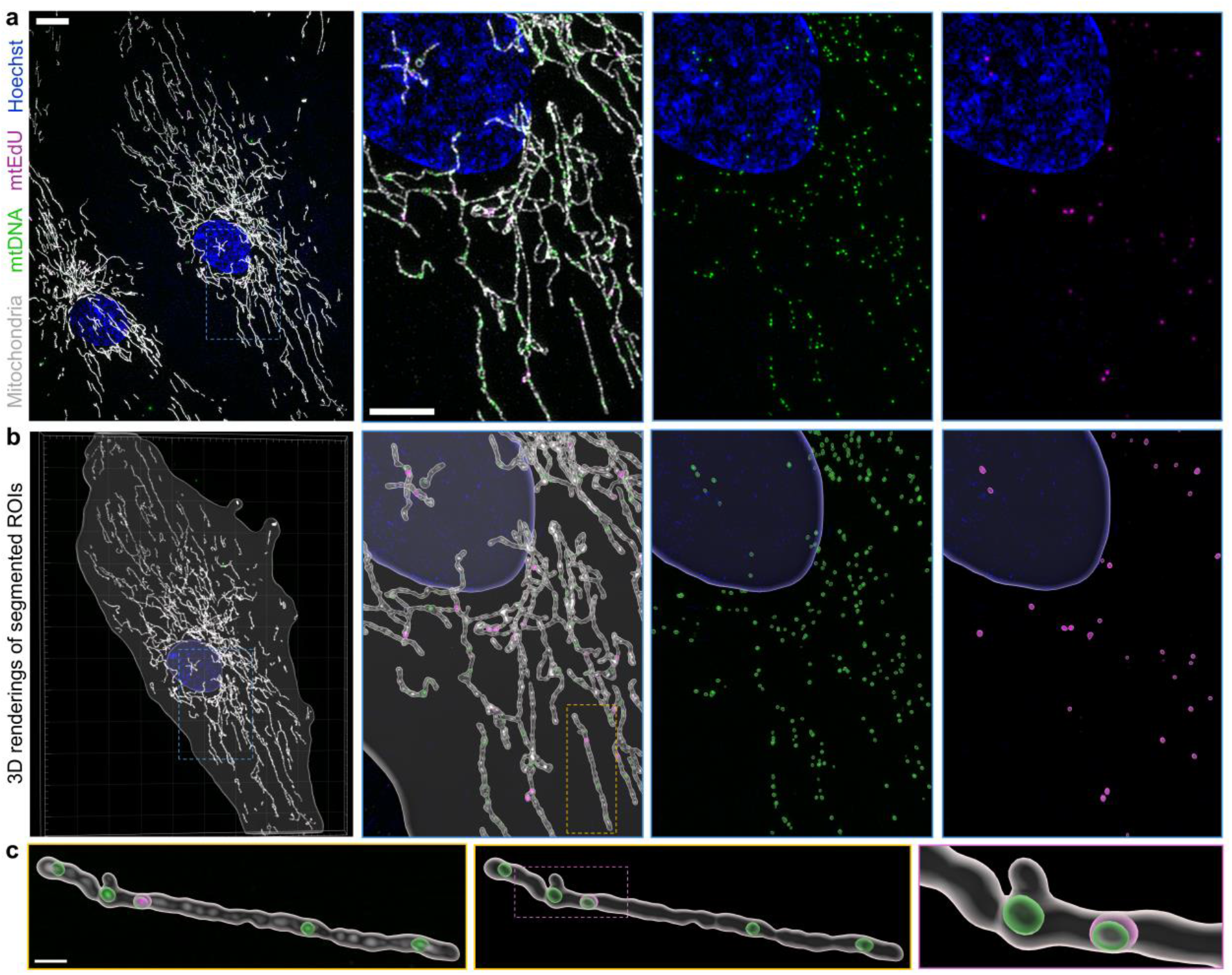
Approach for 3D super-resolution imaging and segmentation of total and replicating mtDNA in quiescent primary human fibroblasts. **a**, Representative 3D lattice SIM image of quiescent primary human fibroblasts, treated with 10 µM EdU for 7 h, prior to fixation and immunostaining, with Hoechst labelling nuclei (blue), anti-TOMM20 labelling mitochondria (gray), anti-DNA labelling the whole population of mtDNA (green), and EdU labelling the subset of replicating nucleoids (mtEdU) (magenta). Machine learning was used to estimate cellular volume (transparent) from background FI of all four channels. Scale bars = 20 µm (left), 5 µm (middle, inset). **b**, Overlaid renderings of cellular, mitochondrial, and nucleoid volumes. **c**, Individual mitochondrion showing 3D segmentation of organelle (gray) and nucleoids (green), with and without EdU incorporation (magenta). Scale bar = 500 nm.

Whether individual mammalian cells regulate absolute mitochondrial genome levels or adjust them according to specific parameters has remained a key question in mitochondrial biology. Analyzing the relationship between total nucleoid number, mitochondrial network volume, and cellular volume revealed strong pairwise Pearson correlations (*r* = 0.820 − 0.950, *p* < 10^−4^) in individual quiescent primary human fibroblasts as well as various proliferative mammalian cell types (Fig. 2a–l), confirming that patterns previously observed in proliferative contexts^33^ also extend to nondividing cells. Genetic perturbations of mitochondrial dynamics have been widely reported to affect mtDNA CN^34–43^. Assessing the morphology of the total mitochondrial network volumes, we did not find that the 3D shape descriptors sphericity and ellipticity correlated with nucleoid number in untreated, baseline conditions (Extended Data Fig. 1d–i). Together, these data indicate both nondividing and proliferative mammalian cells control nucleoid density rather than absolute nucleoid number.

**Figure 2:**
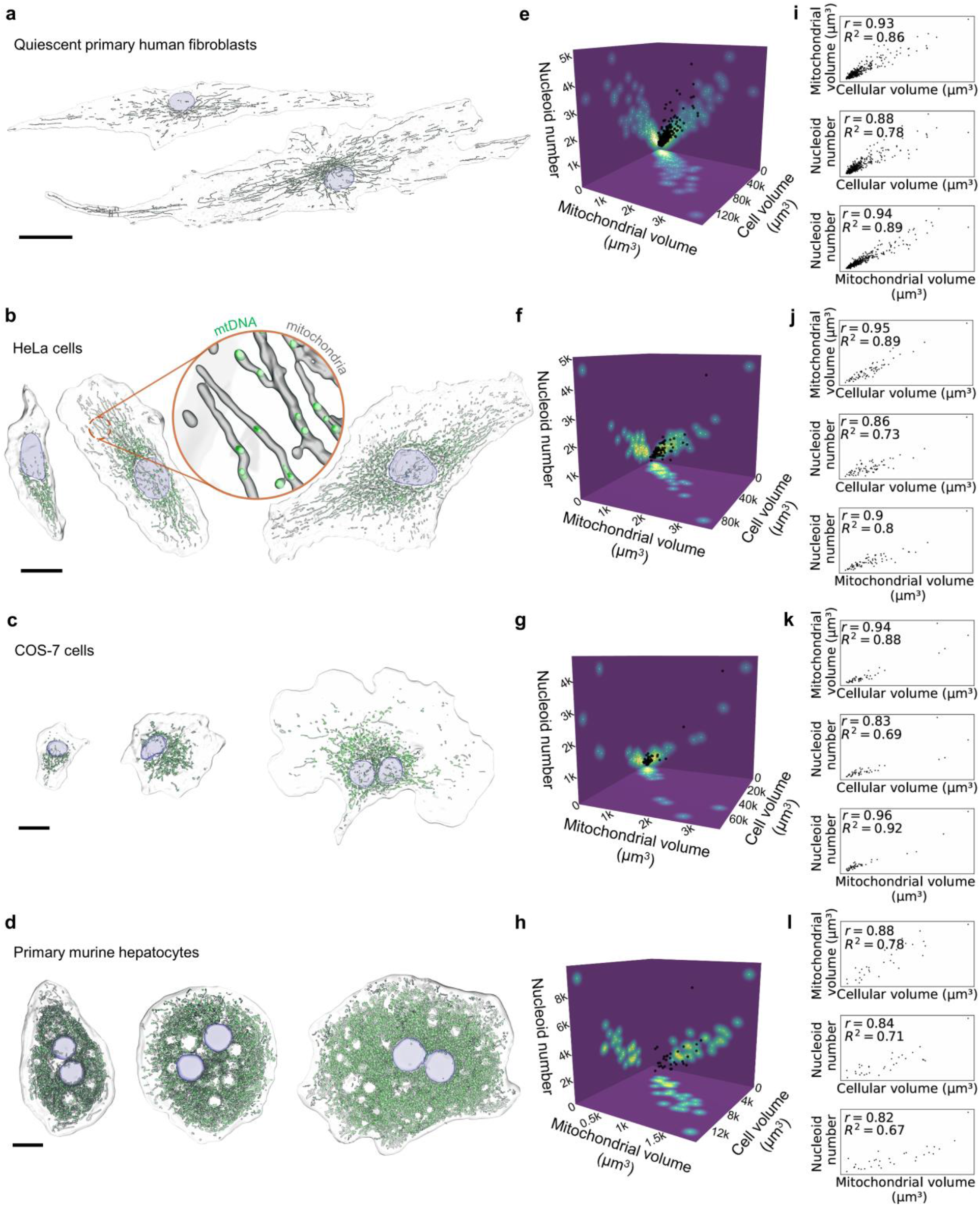
Nucleoid number correlates linearly with mitochondrial network and cellular volumes in a variety of mammalian cell types. **a**, Representative lattice SIM renderings of immunostained quiescent primary human fibroblasts, with Hoechst (blue), anti-TOMM20 (gray), and anti-DNA (green) labelling nuclei, mitochondria, and mtDNA, respectively. Cellular boundaries (transparent) were estimated using machine learning. Scale bar = 20 µm. **b**–**d**, Representative lattice SIM renderings of live HeLa (b) and COS-7 (c) cells, and Airyscan 2 renderings of live primary murine hepatocytes (d), stained with SYBR Green labelling nuclei (blue) and mtDNA (green) and TMRM labelling mitochondria (gray). CellMask Plasma Membrane Stain was used to label plasma membranes (b, c), and machine learning was used to estimate cellular boundaries (d). Scale bars = 20 µm for HeLa and COS-7 cells; scale bar = 10 µm for hepatocytes. **e**–**h**, 3D scatter plots of nucleoid number, mitochondrial network volume, and cellular volume, with 2D heatmap projections of fibroblast (e), HeLa (f), COS-7 (g) cells, and primary murine hepatocytes (h), illustrating multidimensional linearity. **i**–**l**, pairwise plots of nucleoid number, mitochondrial network volume, and cellular volume of fibroblast (i), HeLa (j), COS-7 (k) cells, and primary murine hepatocytes (l), showing pairwise linearity. Data reflect 3D analysis of 381 (a, e, i), 73 (b, f, j), 48 (c, g, k) cells from 3 independent experiments; and 31 cells (d, h, l) from ≥ 3 biological replicates. Statistical tests: Pearson correlation coefficient (*r*) and coefficient of determination (*R*^2^) are shown (I to L). *P*-values for all correlations were *p* < 10^−4^.

### Mitochondrial network volume controls nucleoid density in individual cells

Having established that nucleoids maintain a constant density within the cell (Fig. 2), we next assessed whether this specifically reflects the mitochondrial network volume or cellular volume. For all cell types examined, except primary murine hepatocytes, we observed that both the Pearson correlation and the r-squared value between nucleoid number and cellular volume were smaller than those between both nucleoid number and mitochondrial network volume, or mitochondrial network volume and cellular volume (Fig. 2i–l). This suggested that cellular volume is not the prime regulator of nucleoid number but influences mitochondrial network volume, which in turn determines nucleoid number. We hypothesized, therefore, that non-proliferative cells endeavor to maintain a constant nucleoid density within the mitochondrial network volume, rather than the cellular volume, as in proliferative yeast cells^33^. (The possibility that nucleoid number scales with mitochondrial network volume according to a scaling factor was also explored; see Supplementary Information Section 1.1.)

To examine this hypothesis in steady-state conditions, we ran three separate conditional independence statistical tests (KCIT^44^, GCM^45^, and CMIKnn^46^)—assessing whether two variables are still dependent when accounting for the influence of a third—on the 381 quiescent primary human fibroblasts (see Methods, Supplementary Information 7.2). All three tests placed markedly less weight on the connection between nucleoid number and cellular volume than on the connection between nucleoid number and mitochondrial network volume, or between mitochondrial network volume and cellular volume (Fig. 3a). These findings were recapitulated in COS-7 and HeLa cells as well as primary murine hepatocytes (73, 48, and 31 cells, respectively) (Extended Data Fig. 2a). To gain additional insight, we performed live-cell imaging with lattice light-sheet microscopy, obtaining 4D longitudinal data of 19 individual, quiescent primary human fibroblasts sampled every 2 hours for 1 day, consisting of 247 data points in total (Fig. 3b, c, and Supplementary Video 1). The distributions of nucleoid number, mitochondrial network volume, and cellular volume did not significantly change over time (*p* = 1.0, *p* = 0.81, *p* = 0.81, respectively) (Fig. 3d), with the same behavior also observable at shorter time scales with a finer temporal resolution (Extended Data Fig. 3a–c, and Supplementary Video 2), indicating steady-state conditions. The fact that the cells were in steady state allowed us to test for both linear^47^ and neural Granger causality^48^ (see Methods and Supplementary Information 7.1), each of which assesses whether future values of one time series can be predicted using past values of another. Using the longer-time-scale data, neither method placed a Granger-causal connection between nucleoid number and cellular volume, but both corroborated that there are Granger-causal links between nucleoid number and mitochondrial network volume, and between mitochondrial network volume and cellular volume (Fig. 3e–g). Intriguingly, variations in the size of the mitochondrial network Granger cause variations in cell size (as well as vice-versa), a pattern that has been identified in proliferative cells in both mammals^49^ and yeast^50^. While individually these approaches should be interpreted with caution, the overall pattern indicates, in mammalian cells, nucleoid density is primarily regulated by mitochondrial network volume^51^ and ranges from 1 to 3 nucleoids per cubic micron (Extended Data Fig. 3d).

**Figure 3:**
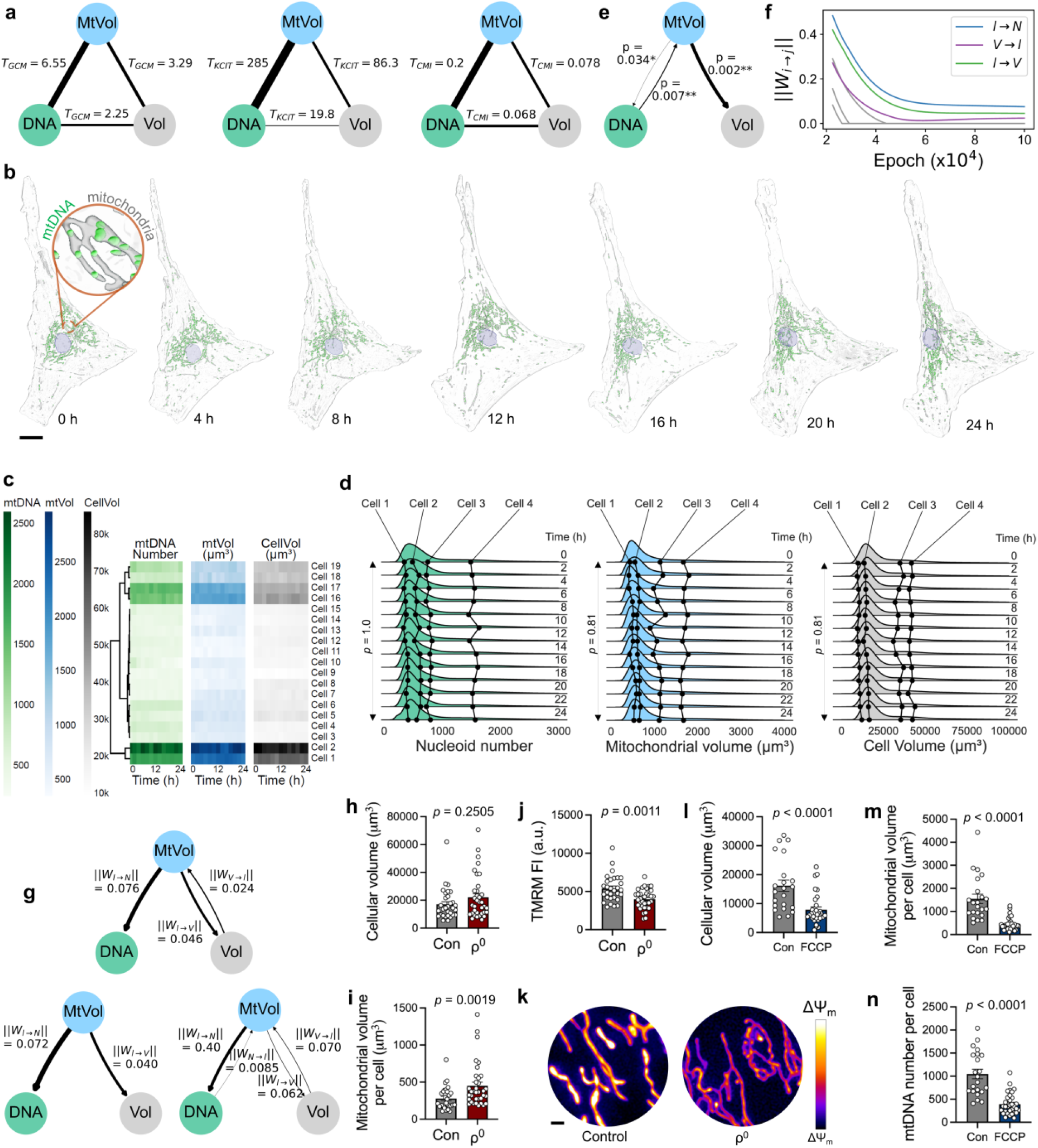
Nucleoid number and cellular volume interact indirectly through mitochondrial network volume sustained by mitochondrial membrane potential (ΔΨ_m_). **a**, Graphical representations of the results of conditional independence testing. Edges are weighted and labelled by test statistic, showing a weaker association between nucleoid number and cellular volume for all tests; *left:* GCM, *middle:* KCIT, *right:* CMIknn. **b**, Representative, 4D lattice light-sheet renderings of a single quiescent primary human fibroblast, showing cell (transparent), nucleus (blue), mitochondria (gray), and mtDNA (green). Scale bar = 20 µm. **c**, Heatmaps of nucleoid number, mitochondrial network volume, and cellular volume for each cell measured during 4D lattice light-sheet imaging, with dendrogram, showing cells of varying sizes in our sample and clear visual correlation between their mtDNA number and volumes. **d**, Distribution of nucleoid number, mitochondrial network volume, and cellular volume over time from 4D lattice light-sheet imaging data, with plots from representative single cells overlaid, demonstrating minimal changes in cellular volume, total mitochondrial volume, and nucleoid number, and mathematically indicating a statistically insignificant change in distribution between 0 and 24 h. **e**, Graphical representation of the results of linear Granger causality, with edges labelled by *p*-value, weighted inversely to *p*-value, showing no statistically significant connection between nucleoid number and cellular volume. **f**, Causal parameters ||*W*_*i→j*_||, signifying the strength of the signal from *i* to *j* (see Methods and Supplementary Informat ion 7.1.2), over time during neural Granger training for regularization parameter *λ* = 3 × 10^−3^. Non-causal connections are sent to 0, whereas causal connections plateau above 0. Nucleoid number, mitochondrial network volume, and cellular volume are labelled *N, l, V*, respectively. **g**, Graphical representations of the results of neural Granger causality for three different regularization parameters *λ*, weighted and labelled by the causal parameters (see Methods and Supplementary Information 7.1.2), showing no neural Granger causal connection between nucleoid number and cellular volume for all parameterizations. **h**–**j**, Results of ρ^0^ cells: mtDNA depletion does not affect cellular volume (h), increases mitochondrial network volume (i), and decreases but sustains ΔΨ_m_ (J). **k**, Live-cell lattice SIM maximum-intensity projections of mitochondria labelled with TMRM. Fire LUT indicates greater polarization of yellowish white mitochondria. Scale bar = 1 µm. **l**–**n**, ΔΨ_m_ controls cellular (l) and mitochondrial network (m) volume as well as nucleoid (n) number. Data reflect 3D analysis of 381 (a), 67 (h–k), and 54 (l–n) cells, and 4D analysis of 19 (b–g) cells from 3 independent experiments. Data (h–j, l–n) are mean ± error of the mean (s.e.m.). *P*-values were determined using Analysis of Variance (ANOVA) partial *F*-tests (e), two-tailed Student’s *t*-tests (h–j, l–n), and Kolmogorov-Smirnov (KS) tests (d).

We next addressed directly whether mtDNA CN influences cellular volume by examining mtDNA-null (ρ^0^) human fibroblasts immediately following cell-cycle arrest. We observed that, despite the lack of mtDNA, ρ^0^ cells maintained their size but exhibited greater mitochondrial network volume (Fig. 3h, i, and Extended Data Fig. 4a–c), aligning with our earlier conclusions. A peculiarity of mtDNA-depleted cells is that, while they are incapable of OXPHOS^52^, they nevertheless expend energy from glycolysis to maintain a mitochondrial membrane potential (ΔΨ_m_)^53^ (Fig. 3j, k). Mitochondria not only generate ATP but also harbor key enzymes associated with numerous metabolic pathways^54^; moreover, the ΔΨ_m_ is critical for mitochondrial protein import^55–59^. We conjectured, therefore, that mitochondria in ρ^0^ cells maintain ΔΨ_m_ to support their diverse roles in cellular homeostasis, including the maintenance of cell size.

To test this hypothesis, we treated control quiescent primary human fibroblasts with the uncoupler carbonyl cyanide-p-trifluoromethoxyphenylhydrazone (FCCP) to dissipate the ΔΨ_m_. Depolarization not only induced cells to shrink but also decreased mitochondrial network volume and nucleoid number (Fig. 3l–n, and Extended Data Fig. 4d, e). Probing further the bioenergetic underpinnings of mtDNA CN regulation, we challenged these fibroblasts by culturing them in phosphate buffered saline (PBS). The cells shrank, accompanied by a decline in mitochondrial network volume and nucleoid number (Extended Data Fig. 4f–j). Together, these data indicate that, although cellular and mitochondrial network volumes are strongly correlated with nucleoid number, mtDNA is not essential for their maintenance; however, these basic parameters are regulated by ΔΨ_m_ as well as nutrient availability.

### mtDNA molecules replicate and degrade at 1.5% per hour in non-proliferative cells

Having determined that mammalian cells specifically maintain nucleoid density in proportion to total mitochondrial volume, we proceeded to investigate mtDNA replication kinetics in quiescent primary human fibroblasts by monitoring the incorporation of EdU, the nucleoside analogue of thymidine, for different durations (Fig. 4a). We found that cells with larger mitochondrial network volumes contained more EdU-labelled nucleoids (mtEdU), in direct proportion to each other (Fig. 4b), from which we deduced that the cellular mtDNA replication n rate scales in proportion to mitochondrial network volume and hence in proportion to nucleoid number (Fig. 2). This indicates a per-capita replication rate that is on average constant and in agreement with our modelling (Fig. 4b). We observed that mtEdU number increased over the course of the time points measured (Fig. 4c, d), while nucleoid number, cellular volume, and mitochondrial network volume remained stable (Fig. 4c–e). The measured rate of mtEdU increase closely matched the mathematical model (Fig. 4f), from which we inferred a per-capita mtDNA replication rate of 1.5 ± 0.2% per hour (Fig. 4g). For a cell of average size with 900 nucleoids, this corresponds to 13.5 ± 2 replication events per hour (Fig. 4c).

**Figure 4:**
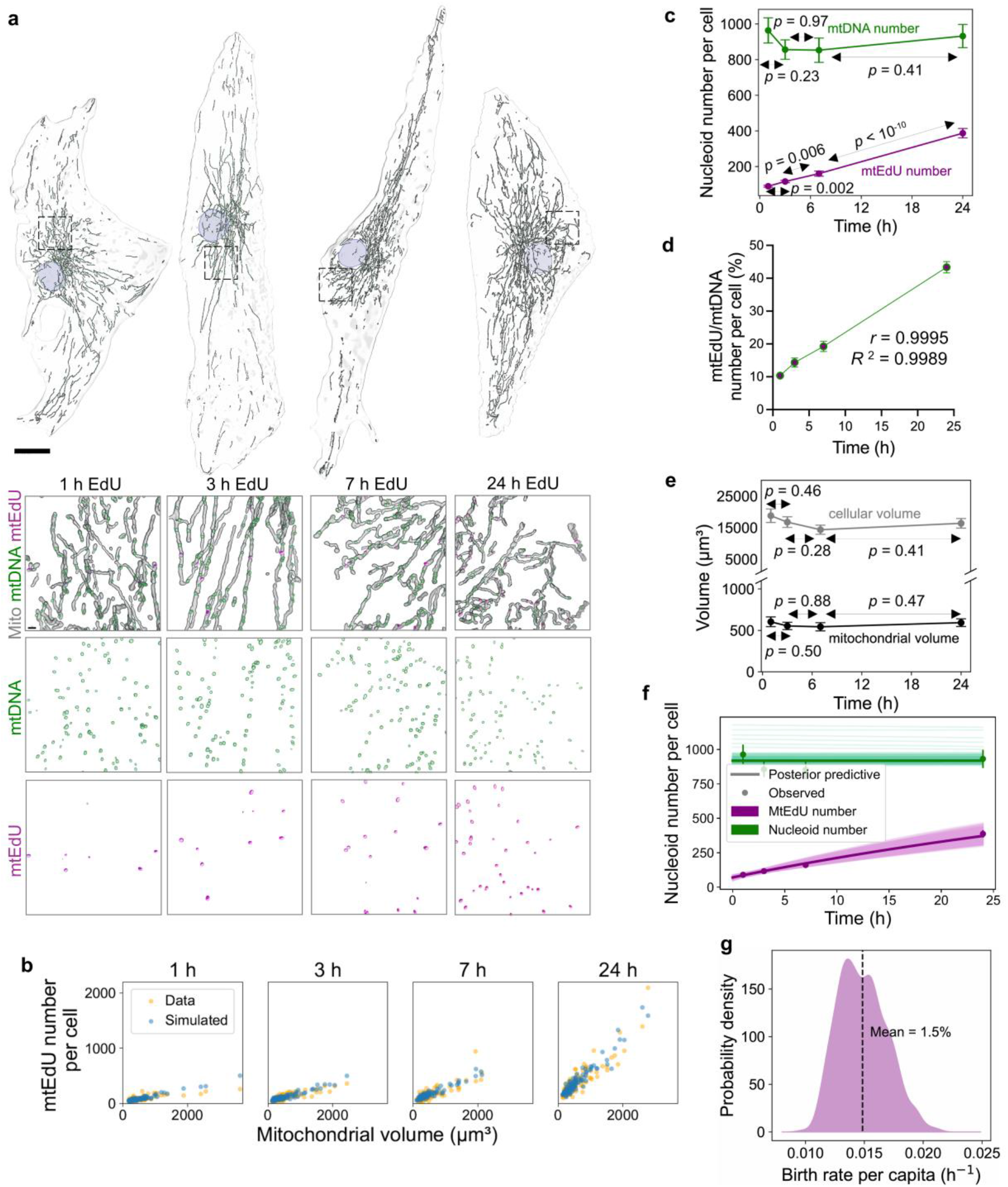
Mitochondrial DNA molecules replicate at a per-capita rate of 1.5% per hour in quiescent primary human cells. **a**, Representative lattice SIM renderings of quiescent primary human fibroblasts treated with 10 µM EdU for 1, 3, 7, and 24 h, prior to fixation and immunostaining. *Top:* Renderings of whole cells (transparent), showing nuclei (blue), mitochondria (gray), mtDNA (green), and mtEdU (magenta). Scale bar = 20 µm. *Bottom:* insets of cells at different time points showing details of mitochondria, mtDNA, and mtEdU. Scale bar = 1 µm. **b**, mtEdU levels scale linearly with mitochondrial network volume, indicating a global rate which scales linearly with total mitochondrial volume, or a constant (on average) per-capita rate. **c–e**, MtEdU levels significantly increase over time, while cellular and mitochondrial network volumes as well as total mtDNA remain stable. **f**, MtEdU and mtDNA over time with model fit. Dots and error bars are observed mean mtEdU and mtDNA. Lines indicate simulated trajectories from our model (posterior predictive distribution), with thick lines indicating the mean of all model simulations (posterior predictive mean). **g**, Posterior probability distribution of the per-capita replication and degradation rate, showing a mean of 1.5% per h. Data reflect 3D analysis of 381 cells from 3 independent experiments, respectively. Data are mean ± s.e.m. Statistical tests: Pearson correlation coefficient (*r*) with *p*-values from Welch’s *t*-test are shown (c, e).

Given the stability of the nucleoid number both at the single-cell and population level (Fig. 3b–d, 4c, f, and Extended Data Fig. 3a–c), the rate of mtDNA replication within the quiescent fibroblasts must be balanced by an equivalent rate of degradation. To directly assess the levels of mtDNA degradation in whole quiescent single cells, we used 3D lattice SIM to image nucleoids enveloped by LC3-positive vesicles (Fig. 5a and Extended Data Fig. 5a), representing an early-to-intermediate phase of autophagic clearance^60^. First, we found that the number of nucleoids being degraded significantly exceeded what we would expect from random overlap between nucleoids and LC3 (*p* = 0.001, Fig. 5b, Methods). Next, we found a significant correlation (*r* = 0.485, *p* = 0.019) between the number of nucleoids being degraded and mitochondrial network volume (Fig. 5c), confirming that the global degradation rate also scales with mitochondrial network volume, with an average of 14.22 ± 3.13 nucleoids in the process of degradation (Fig. 5d), equivalent to 1.7 ± 0.4% per cell (Fig. 5e). LC3-positive mtDNA number also significantly correlated with total nucleoid number per cell (Extended Data Fig. 5b). From the alignment between the number of nucleoids actively degrading (14.22 ± 3.13) and the number of replication events per hour in the average cell (13.5 ± 2), it directly follows that the number of actively degrading nucleoids and the number of degradation events per hour also align (since the latter matches the number of replication events per hour). This implies, assuming all nucleoids degrade through this pathway, that each nucleoid takes approximately one hour to degrade, roughly in accord with prior work on rat kidney cells^61^; by contrast, assuming not all nucleoids degrade through this pathway skews this estimate to be unreasonably large (Methods, Supplementary Information 5.3). This prompts us to conclude that the majority of nucleoids degrade through this pathway, in agreement with observations of selective enrichment of nucleoids in autophagosomes in neurons^62^ and different cell lines^63^.

**Figure 5:**
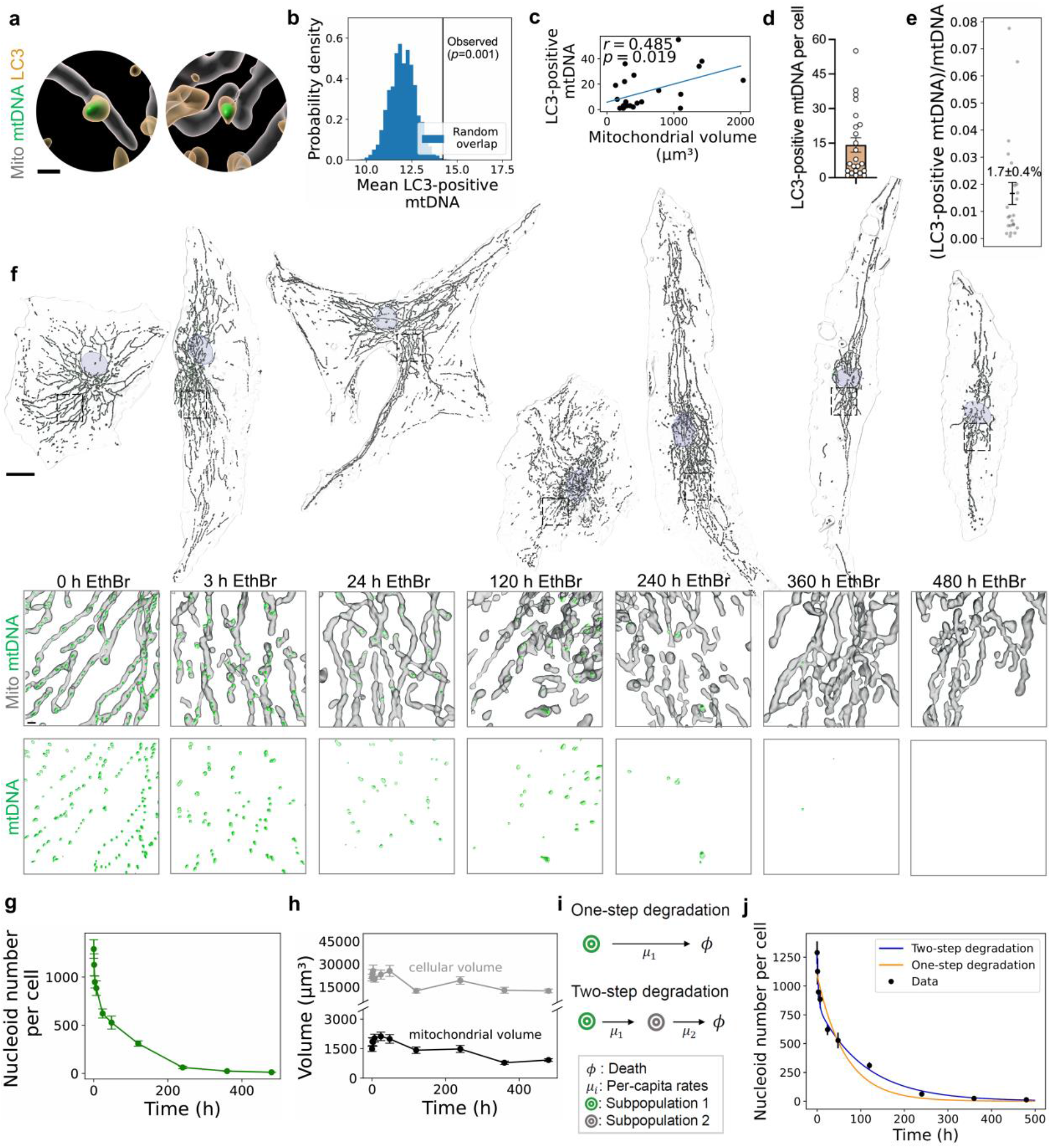
A potential two-step degradation mechanism governs mtDNA turnover. **a**, Representative lattice SIM renderings of mitochondria (gray) in quiescent primary human fibroblasts, showing mtDNA molecules (green) enveloped by LC3 puncta (light orange). Scale bar = 1 µm. **b**, Mean number of LC3-positive mtDNA molecules significantly exceeds the distribution expected under random overlap between nucleoids and LC3 (Methods). **c**, Number of LC3-positive mtDNA molecules scales linearly with mitochondrial network volume. **d**, Number of LC3-positive mtDNA molecules per cell. **e**, Percentage of LC3-positive mtDNA molecules per cell. **f**, Representative lattice SIM renderings of quiescent primary human fibroblasts treated with 500 nM ethidium bromide up to 480 h, prior to fixation and immunostaining. *Top:* Renderings of whole cells (transparent), showing nuclei (blue), mitochondria (gray), and mtDNA (green). Scale bar = 20 µm. *Bottom:* insets of cells at different time points showing details of mitochondria and mtDNA. Scale bar = 1 µm. **g, h**, MtDNA, cell volume, and mitochondrial network volume following inhibition of mtDNA replication, with mtDNA depletion resembling an exponential decay. **i**, Diagram illustrating one- and two-step degradation models. **j**, Decrease in mtDNA consistent with a two-step degradation model. One-step degradation, i.e., an exponential decay, gives a degradation rate of 1.55 ± 0.3%. Data reflect 3D analysis of 23 (a–e) and 472 (f–j) cells from 3 independent experiments, respectively. Data are mean ± s.e.m. Statistical tests: Pearson correlation coefficient (*r*) with *p*-value from two-tailed Student’s *t*-test are shown (c).

Exploring the kinetics of mtDNA degradation further, we completely inhibited mtDNA synthesis in quiescent primary human fibroblasts using ethidium bromide (Extended Data Fig. 5c) and monitored nucleoid levels over time (Fig. 5f). A precipitous drop in nucleoid number occurred in the first 24 hours, followed by a gradual disappearance over the course of 20 days, as CN approached zero (Fig. 5g, h). Fitting an exponential decay to this curve gives an mtDNA degradation rate of 1.55 ± 0.3%, matching that of replication; however, this curve is more consistent with a two-step degradation mechanism whereby a subpopulation of mtDNA degrades rapidly, while the rest of the mtDNA molecules slowly move into this subpopulation before degrading (Fig. 5i, j).

To corroborate these findings in a different cell type, species, and physiological contexts, we assessed primary murine macrophages with and without lipopolysaccharide (LPS) stimulation (Extended Data Fig. 6a, b). LPS stimulation resulted in a rapid increase in cellular as well as total mitochondrial volumes (Extended Data Fig. 6c, d), followed by an increase in nucleoid number (Extended Data Fig. 6e), consistent with previous reports^64^. Fitting a deterministic birth-death model (see Methods, Supplementary Information 5.1) to the mtEdU curves (Extended Data Fig. 6f, g) revealed a basal per-capita rate of 0.9 ± 0.2% per hour (Extended Data Fig. 6h). Following LPS stimulation, the per-capita birth rate increased to 10 ± 3.9% per hour before exponentially decreasing within the 26-hour period of monitoring (Extended Data Fig. 6i, j). Together, these single-cell data indicate that mtDNA molecules in markedly different cell types from different species replicate at similar rates under baseline conditions, and these rates are subject to physiological stimuli.

### Spatiotemporal segregation in mtDNA replication and degradation

The results of the ethidium bromide experiment (Fig. 5f, g) suggest that there are distinct subpopulations of mtDNA with unique replication and degradation properties (Fig. 5i, j); we therefore decided to probe more deeply and at scale. Since the ethidium-bromide-induced nucleoid decay curve could be explained by a non-constant degradation rate, we performed a series of pulse–chase experiments in which we pulsed the quiescent fibroblasts with EdU for 24 hours, followed by a chase with thymidine for different durations (Fig. 6a). From the start of the experiment, a replicating nucleoid will incorporate EdU into a single strand of each daughter mtDNA molecule (Fig. 6b). If either of these daughters undergoes an additional round of replication, a new set of daughters will emerge, one with both strands tagged by EdU, the other with still a single strand tagged, and so on. Removal of EdU during the chase reverses this process, so, for example, a doubly tagged mtDNA molecule would create two singly tagged daughters, etc.

**Figure 6:**
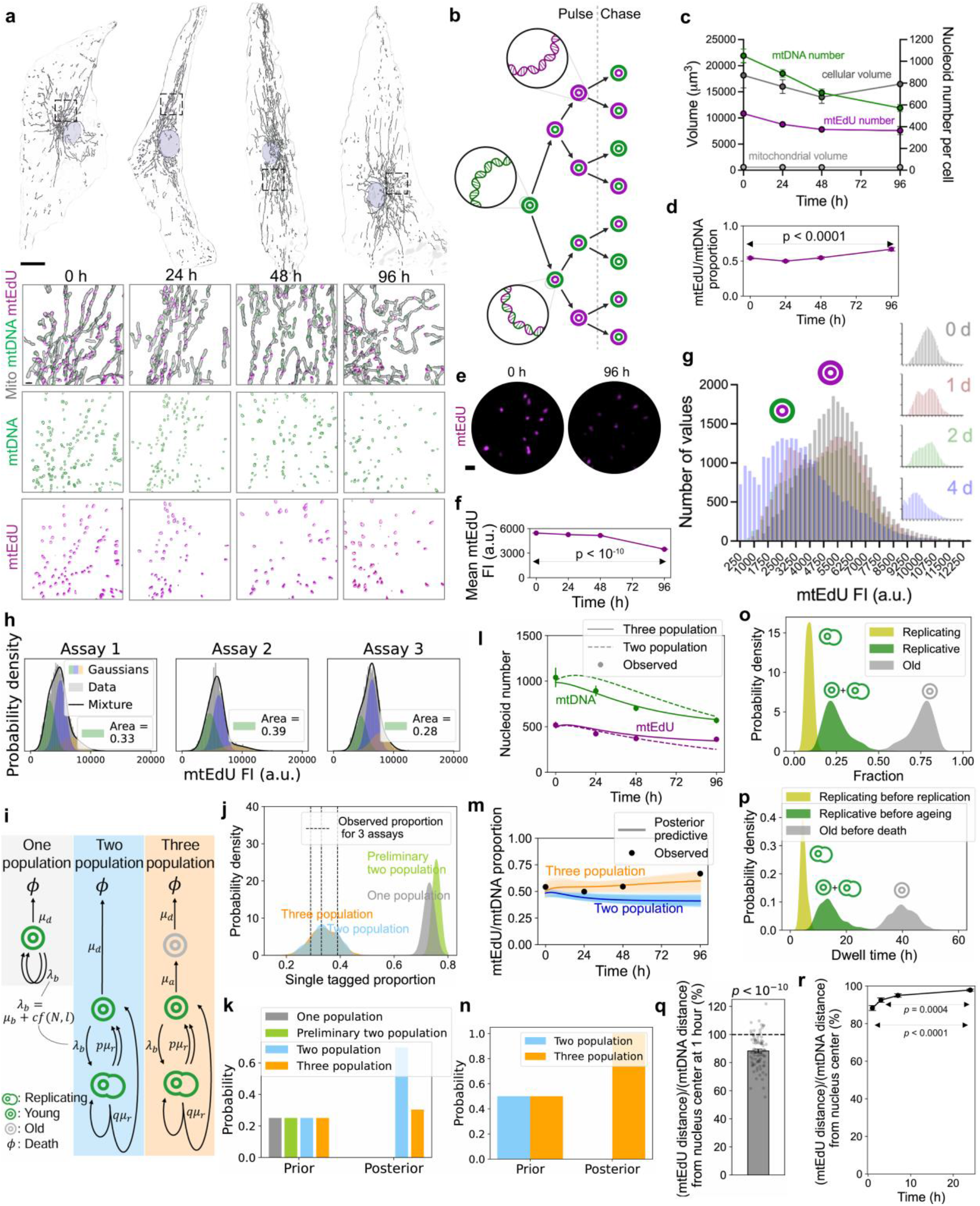
Newly synthesized mtDNA molecules are closer to the nucleus, undergo preferential replication, and are less likely to degrade. **a**, Representative lattice SIM renderings of quiescent primary human fibroblasts treated with 10 µM EdU for 24 h, followed by a chase with 1 mM thymidine for 0, 24, 48, and 96 h, prior to fixation and immunostaining. *Top:* Renderings of whole cells (transparent), showing nuclei (blue), mitochondria (gray), mtDNA (green), and mtEdU (magenta). Scale bar = 20 µm. *Bottom:* insets of cells at different time points showing details of mitochondria, mtDNA, and mtEdU. Scale bar = 1 µm. **b**, Cartoon of EdU incorporation into and clearance from replicating mtDNA molecules during a pulse–chase experiment. Green and magenta circles represent untagged and EdU-tagged mtDNA strands, respectively. **c**, Total mtEdU and mtDNA number per cell decreases slowly over 96 h, but mtEdU decreases at a slower rate than mtDNA. **d**, Following the chase, proportion of mtEdU to total mtDNA significantly increases over time. **e, f**, Representative 3D lattice SIM images of mtEdU molecules (e), showing significant decrease in fluorescence intensity (FI) (f) over time. Scale bar = 1 µm. **g**, Frequency distribution of FIs of individual mtEdU molecules from different time points, displaying two prominent peaks, showing the time-dependent change of mtDNA molecules harboring EdU in both strands (right peak) to only a single strand (left peak). **h**, Gaussian mixture model fits to each assay of the 0-day FI data. Area under the first Gaussian indicates that a mean of 33 ± 3% mtEdU molecules are singly tagged at this time point. Second and third Gaussians represent doubly and more than doubly tagged nucleoids, respectively. **i**, Diagrammatic representations of the one-, two-, and three-population models. Arrows indicate events moving a molecule from one subpopulation to another, with label being the per-capita rate of those events. Two arrows indicate doubling, and each arrow denotes the destination of one of the two daughter molecules. **j**, Posterior predictive probabilities of singly tagged mtDNA proportion according to each model. Only the models with preferential replication of newly synthesized mtDNA fit the observed proportions. **k**, ABC model selection following fitting to the pulse and FI data, selecting the two models with preferential replication. **l, m**, Three-population model with newly replicated molecules spared from degradation fits mtEdU/mtDNA (l) and mtEdU, mtDNA (m) data better than the two-population model. **n**, ABC model selection following fitting to the pulse–chase and FI data, selecting the three-population model, indicating newly replicated molecules are less likely to degrade. **o**, Population proportion posteriors from the three-population model, predicting a mean of 75 ± 8% mtDNA in the old population, 9 ± 2% actively replicating, and 25 ± 8% replicative—in either the young or actively replicating population (mean ± standard deviation of the posteriors). **p**, Dwell time posteriors from the three-population model, predicting a mean of 14.4 ± 4.4 h in the replicative populations before ageing, 40.0 ± 4.6 h in the old population before degrading, and 4.8 ± 1.0 h undergoing a single replication event (mean ± standard deviation of the posteriors). **q**, Newly replicated mtDNA molecules are closer to the nucleus than the general population of mtDNA. **r**, Over time, newly replicated mtDNA molecules are distributed to the cellular periphery. Data reflect 3D analysis of 265 (a, c–h, j–n) cells from 3 independent experiments, including 106,176 mtEdU puncta (g, h), and maximum-intensity-projection analysis of 381 cells from 3 independent experiments (q, r). *P*-value (q) was determined through a one-tailed Student’s *t*-test, while *p*-values (r) were determined using a one-way ANOVA with Tukey’s multiple comparison test. Data are mean ± s.e.m.

In keeping with our earlier findings, the prolonged absence of FBS led to a decrease in nucleoid number during the chase (Fig. 6c). This did not occur during the pulse phase, leading to an imbalance in replication and degradation that we exploited to probe the dynamics of degradation. During the chase, although the nucleoid numbers dropped steeply, the levels of mtEdU dropped only slightly (Fig. 6c), which resulted in the proportion of EdU-tagged molecules to total nucleoids significantly increasing over time (Fig. 6d). Consistent with the pattern of nucleoid clearance following ethidium bromide treatment (Fig. 5f–j), this dynamic suggests different subpopulations of mtDNA undergo degradation at different rates.

Given that EdU labels molecules that have replicated recently, we hypothesized that new mtDNA molecules are less likely to degrade than non-recently replicated, or *old*, molecules. To explore this further, we examined the fluorescence intensities (FIs) of the mtEdU-tagged puncta at each time point. Following the chase, we found that the average FI of mtEdU puncta significantly decreased over time (Fig. 6e, f), and the full distribution of mtEdU-tagged molecules at each time point, representing a total of 106,176 mtEdU puncta, contained two peaks. The left peak was more prominent in the latter time points, but the right peak was more prominent in the early time points (Fig. 6g). Given our use of 3D super-resolution microscopy, it is unlikely that this phenomenon was due to unresolved nucleoids, and if these peaks instead reflected nucleoids with one versus two mtEdU puncta, we would not expect to see the right peak be prominent in any time point, since nucleoids with two mtDNA molecules are the minority of the population^11^ (Supplementary Information 3.1.3). The most parsimonious explanation for these observations, therefore, is that they reflect singly and doubly labelled mtEdU molecules. During the chase phase, the average FI decreases as the doubly tagged mtEdU molecules replicate again, making a larger proportion of singly tagged genomes. Consistent with this, the prominence of the right peak at the first time point indicates that the majority of mtEdU molecules were doubly tagged following the pulse and thus had replicated at least twice. We confirmed this interpretation using Gaussian mixture modelling (Extended Data Fig. 7a–d), estimating that 33 ± 3% of mtEdU molecules were singly tagged following the pulse (Fig. 6h). Based on these findings, we hypothesized that newly formed nucleoids had a greater tendency to continue replicating compared to an aged population, which had not recently replicated.

To test this independently, we constructed four stochastic mechanistic models for nucleoid population control of increasing complexity (Fig. 6i). We called our null model the *one-population* model, consisting of a constant per-capita death rate *μ*_*d*_ and a per-capita birth rate *λ*_*b*_ = *μ*_*d*_ + *f*, where *f* is a functional feedback term embodying the need (Fig. 3) to ensure nucleoid count per unit of mitochondrial volume is controlled^65^ (see Methods and Supplementary Information 1.1). The *preliminary two-population* model added a replicating subpopulation, and the *two-population* model expanded on the preliminary model by assigning a probability *q* = 1 − *p* that one of the daughters of a molecule that has just replicated (i.e., exited the replicating population) begins replicating again (i.e., re-enters the replicating population). The *three-population* model extends the two-population model by adding an old subpopulation, which molecules must age into before undergoing degradation.

Only the two- and three-population models (Fig. 6i), which incorporate preferential replication of newly replicated molecules, fitted data from the pulse phase of the experiment where most nucleoids were doubly tagged (Fig. 6j), a result confirmed by model selection (Fig. 6k). Subsequently fitting these two models to the chase phase of the experiment showed that only the three-population model, which also incorporated the preferential degradation of old molecules, could explain the observed mtEdU drop (Fig. 6l, m), which we again confirmed with a model selection (Fig. 6n). All models were fit on two experimental assays, the three-population model was validated on a third, and measurement error was taken into account (see Methods and Supplementary Information Sections 3.1.1, 4.6). Overall, these findings provide strong evidence for unique subpopulations of mtDNA preferentially replicating and degrading relative to their ages.

Having selected the three-population model, we were able to probe the model fit to gain new biological insights and predictions. The model predicts a mean of 75 ± 8% of mtDNA molecules are in the old population, 9 ± 2% are actively replicating, and 25 ± 8% of molecules are *replicative*—in either the young or actively replicating population (Fig. 6o). Following a replication event, one of the daughter mtDNA molecules has a probability q = 0.67 ± 0.07 of recommencing replication. Moreover, we estimate that mtDNA molecules spend a mean of 14.4 ± 4.4 hours in the replicative populations before entering the quiescent phase (moving to the old population), 40.0 ± 4.6 hours in the old population before degrading, and 4.8 ± 1.0 hours undergoing a single replication event (Fig. 6p) (see Supplementary Information Section 8).

Finally, we asked whether the replicating, young, and old subpopulations of mtDNA are in any way spatially segregated in quiescent primary human fibroblasts. We measured the distance of EdU-tagged nucleoids from the cell nucleus, comparing this to the general population of nucleoids 1 hour after the addition of EdU. Replicating mtDNA molecules were 11.7 ± 1.3% closer to the nucleus, consistent with observations in PC12^66^ and COS-7^31^ cells (Fig. 6q and Extended Data Fig. 7e). At later time points, the mtEdU diffused, or was transported, away from the nucleus (Fig. 6r, Extended Data Fig. 7e). Altogether, these findings highlight a mechanism of spatiotemporal segregation at work within the mtDNA life cycle, where younger mtDNA molecules preferentially replicate closer to the nucleus and are spared from degradation, whereas older, non-replicating genomes migrate to the cellular periphery and are eventually turned over.

## Discussion

The majority of mtDNA mutations are thought to originate from replication errors by Polγ, the mtDNA polymerase^67–70^, and their accumulation is strongly linked to a decline in mitochondrial function, a hallmark of numerous human diseases, including cancer^71^ and neurodegeneration^72,73^. Pathogenic mtDNA mutations are prevalent in the general population^74,75^, but understanding how they build up over time, particularly in postmitotic cells, which are most relevant for aging, requires detailed knowledge of the kinetics by which mitochondrial genomes are copied and degraded; nevertheless, this critical information has been lacking in non-proliferative cells.

Our finding that 1.5% of the cell’s total nucleoid content turns over each hour indicates that for a typical nondividing cell with 1,000 mtDNA molecules, every day involves approximately 360 unique replication events. Given Polγ’s error rate^20,76^ and the 16,569 base pairs of the human mitochondrial genome^4^, this would amount to 12 de novo mtDNA mutations, daily, per cell. Although these mutations would be minority species and thus would be less likely to clonally expand through random drift^77^, a three-to fivefold increase in the number of point mutations, as seen in Polγ mutant mice, is associated with a severe premature aging phenotype and decreased lifespan^78^, and an average of two mutations per mtDNA molecule is sufficient to elicit signs of premature aging and death^22,79,80^. Our findings therefore provide an empirical framework for how these mutations naturally accumulate over the course of an entire human lifespan, contributing to the deterioration in cellular function linked to many age-related diseases and potentially to the aging process itself^81–83^.

It is important to note that the mtDNA flux that we observe in whole non-dividing human fibroblasts using 3D SIM aligns with estimates of the mtDNA replication rate in segments of proliferating human fibroblasts obtained by 2D STED^26^, suggesting that, while the spatial resolution of SIM is slightly less than that of STED, it is nevertheless sufficient for accurately quantifying nucleoid population dynamics. By focusing on quiescent primary human fibroblasts, we provide evidence that newly replicated mtDNA molecules not only represent a spatiotemporally distinct subpopulation but reflect diverging fates of individual mitochondrial genomes based on their ages. While evidence exists for heterogeneous TFAM enrichment at the scale of individual nucleoids^26–29^, affecting local gene expression, there was limited understanding of the duration of mtDNA replication and the rate of interchange between replicative and non-replicative states (see Supplementary Information Section 1.5). Here we provide concrete, model-based estimates of the dwell time, interchange frequencies, pulsatile (i.e., burst-like) replication, and relative sizes of each mtDNA subpopulation, shedding light on the dynamic life cycle of the mitochondrial genome in non-replicative cells.

While mtDNA is thought to have limited capacity for surveillance and repair^8^, the privileged replication and persistence of younger mtDNA molecules hints at a novel quality-control axis: for example, newly replicated mitochondrial genomes will have hadlless time to be exposed to environmental toxins, ionizing or ultraviolet radiation, and reactive oxygen species^84^, ubiquitous byproducts of the OXPHOS machinery embedded in nearby cristae membranes^85^. Simultaneously sparing younger mtDNA molecules from degradation and promoting their replication might curtail the risk of copying damaged or mutated genomes. Temporal sorting of this kind helps explain why most mtDNA mutations stem from proofreading errors, which would not be mitigated by this quality-control mechanism.

Such spatiotemporal regulation of the mitochondrial genome has interesting implications for the physiology of large, highly polarized cells, like neurons, where distal dendrites and synapses may harbor older mtDNA molecules, potentially impacting local mitochondrial function under a range of environmental stresses. Charting the subcellular distribution of young and old nucleoids within individual neuronal architecture could provide additional valuable insights into functional differences of mitochondrial populations already visible in the energetic landscape of the brain^86^.

It is important to note, as well, that a smaller replicative population creates a sustained subcellular genetic bottleneck when compared to the total cellular mtDNA content. This would accelerate the rate at which mutations accumulate under neutral drift, exposing them to selective forces operating at the organelle or cellular level, and thus removing them from the population. Similar mechanisms exist within the germline^87^, and here we show such a process could support mtDNA integrity across the lifespan of the organism.

Although nuclear somatic mutations and mosaicism represent critical drivers of aging^88^, mtDNA mutation is orders of magnitude faster. The absolute kinetics of the mtDNA life cycle in non-replicative cells, highlighted here, are key determinants of the speed at which mtDNA mutations accumulate, crucial for understanding how age-dependent mitochondrial dysfunction underlies cellular decline and death.

## Supporting information

Mathematical Modelling Supplement

## Acknowledgements

We thank the Cambridge Advanced Imaging Centre (CAIC), especially Dr. Martin O. Lenz, for assistance with Airyscan 2 and lattice light-sheet microscopy.

## Author contributions

D.M.W., N.S.J., and P.F.C. conceptualized the study and designed experiments. D.M.W. conducted experimentation. D.M.W. and M.S. performed image analysis. E.M. performed mathematical modelling and inference. A.M.C. generated murine BMDMs. N.B. isolated murine hepatocytes. D.M.W., R.F., M.P.M., J.P., N.S.J., P.F.C. provided resources. D.M.W. and E.M. made the figures and wrote the manuscript, with feedback from P.F.C, N.S.J., and all other authors. The work was supervised and funded by N.S.J. and P.F.C.

## Resource availability

All data are available in the manuscript or the supplementary materials. All materials generated in this study are readily available from the authors. All data supporting the findings of this study are available from the corresponding authors upon request. For the purpose of open access, the authors have applied a Creative Commons Attribution (CC BY) license to any Author Accepted Manuscript version arising.

## Supplementary information

Materials and Methods

Extended Data Fig. 1–7

Supplementary Videos 1 and 2

Mathematical Supplement (including Supplementary Fig. 8–32)

**Figures**

## Methods

### Cell culture

Cells were cultured at 37 °C, 5% CO_2_, and 95% relative humidity. Cell lines were obtained from American Type Culture Collection (ATCC). Primary adult dermal human fibroblasts (PCS-201-012), HeLa, and COS-7 cells were cultured in Dulbecco’s Modified Eagle’s Medium (DMEM), supplemented with 25 mM HEPES (Gibco, 15630056). Quiescent primary human fibroblasts were cultured in HEPES-containing DMEM without FBS.

### Generation of bone-marrow-derived macrophages (BMDMs)

Male C57BL/6 J mice aged 10–18 weeks old were euthanized by cervical dislocation, and death was confirmed by exsanguination. Bone marrow was harvested from the tibia and fibula. Cells were pelleted by centrifugation at 425 × g (1,500 rpm) for 5 min and red blood cells (RBCs) were lysed using a hypotonic RBC lysis buffer (Cambridge Bioscience). The remaining cells were pelleted by centrifugation at 425 × g (1,500 rpm) for 5 min and filtered through a 70-*μ*m nylon mesh. Obtained monocytes were differentiated in DMEM containing 20% L929 supernatant, 10% FBS (Gibco) and 1% penicillin/streptomycin (pen/strep) (Gibco) for 3 days in 10-cm dishes, at which point, additional L929 supernatant (10%) was added, and the cells were differentiated for a further 3 days. BMDMs were scraped, a sample was stained with Trypan blue, counted using an automated cell counter (ThermoFisher) and then 200,000 cells in DMEM containing 10% L929 supernatant, 10% FBS, and 1% pen/strep were plated on glass coverslips in a 24-well plate and left overnight to adhere.

### Isolation of primary hepatocytes from murine liver

Hepatocytes were isolated from 20–25-weeks-old C57BL/6J mice by liberase (Merck, 5401119001) perfusion. Following euthanasia by dislocation of the neck, the liver was perfused *in situ* at 37 °C via the portal vein using a peristaltic pump at a flow rate of 4 mL/min; first with 25 mL of HBSS no Ca^2+^ no Mg^2+^ (ThermoFisher, 14175095) + 0.5 mM EDTA (ThermoFisher, 15575020) + 25 mM HEPES (ThermoFisher, 15630080) followed by 50 mL of HBSS +Ca^2+^ +Mg^2+^ (ThermoFisher, 14025092) + 25 *μ*g/mL liberase (Merck, 5401119001) + 25 mM HEPES (ThermoFisher, 15630080). The liver was then dissected and the gall bladder eliminated. Under sterile conditions, the hepatocytes were released gently from the digested liver by cutting the liver sack and resuspended in cold plating media (DMEM low glucose, ThermoFisher, 11054020 + 5% FCS). After filtration in a 70-µm cell strainer, the cell suspension was spun at 50 × g (with low brake and acceleration to minimize trauma to hepatocytes) for 2 min at 4 °C. To separate the viable hepatocytes from dead hepatocytes and cell debris, the cell pellet was resuspended in 10 mL of plating media and 10 mL of 90% Percoll solution (100% Percoll, Cytiva, 17-0891-01, diluted in 10× PBS, ThermoFisher 70011044) was added. This suspension was mixed thoroughly and spun at 200 × g for 10 min at 4 °C (no brake). The pellet containing live purified hepatocytes was gently resuspended in plating media and spun at 50 × g for 2 min at 4 °C. After resuspension in plating media, the hepatocytes were counted and plated on wells coated with 0.1 mg/mL rat tail collagen I (Merck, C3867). After 3–4 h the dead cells were removed by washing with warm 1× PBS and maintenance media (William’s E media, 32551020) was added. The hepatocytes were kept in maintenance media in a humidified 5% CO_2_ incubator at 37 °C for downstream analyses. Animals used in this study were obtained from Charles River, U.K. This research has been conducted under the Animals (Scientific Procedures) Act 1986, Amendment Regulations 2012, following ethical review by the University of Cambridge Animal Welfare and Ethical Review Body (AWERB).

### Cellular treatments

10 µM carbonyl cyanide-p-trifluoromethoxyphenylhydrazone (FCCP) was used to depolarize mitochondria. Dimethyl sulfoxide (DMSO) (VWR Chemicals, N182) was used as a vehicle in control conditions. Lipopolysaccharide (LPS) derived from *Escherichia coli*, serotype EH100 (Alexis), was kept at 1 mg/mL in phosphate-buffered saline (PBS) at 4 °C, sonicated for 5 min, and used to stimulate macrophages at a final concentration of 100 ng/mL. 500 nM UltraPure Ethidium Bromide (ThermoFisher, 15585011) or 10 µM 2’,3’-Dideoxycytidine (ddC) (Merck, D5782) were used to generate ρ^0^ cells. DMEM containing ethidium bromide or ddC was replaced every 2 days. DPBS, calcium, magnesium (ThermoFisher, 14040133) was used for starvation experiments. Cells were washed 3× with DPBS and to remove residual DMEM at the start of the assay.

### Immunofluorescence

Cells were fixed in prewarmed 5% paraformaldehyde (PFA) in PBS for 15 min at 37 °C and washed 3× with PBS. Cells were subsequently treated with 50 mM ammonium chloride in PBS to quench PFA-induced background fluorescence, washed 3× with PBS, and permeabilized using 0.1–0.5% Triton X-100 (Sigma-Aldrich, T8787) in PBS for 15 min, followed by 3 additional washes with PBS. Cells were blocked with 10% FBS in PBS and incubated overnight at 4 °C with the indicated primary antibodies in a 5% FBS/PBS solution. Cells were then washed 3× with a 5% FBS/PBS solution before incubation for 1 h at room temperature (RT) with Alexa Fluor-conjugated secondary antibodies diluted to 1:1000 in a 5% FBS/PBS solution. Tinfoil was used to cover cells during incubation with secondary antibodies. Cells were then washed 3× with PBS. Cells were mounted using ProLong Diamond Antifade Mountant (ThermoFisher, P36961) and stored at 4 °C prior to imaging.

### Antibodies

To label mitochondria, mtDNA molecules, plasma membrane, and LC3-positive puncta, anti-TOMM20 (Abcam, ab232589 or Santa Cruz Biotechnology, sc-17764), at a dilution of 1:1000, anti-DNA (Sigma-Aldrich, CBL186), at a dilution of 1:500, anti-Sodium Potassium ATPase (Abcam, ab76020), at a dilution of 1:200, and anti-LC3B (Cell Signaling Technology, 83506), at a dilution of 1:500, primary antibodies were used, respectively. Fluorescent secondary antibodies used were Alexa Fluor 555 conjugate (Cell Signaling Technology, 4413S), Alexa Fluor 488 (ThermoFisher, A-21042), Alexa Fluor 647 (ThermoFisher, A-21244), Alexa Fluor 568 (ThermoFisher, A-21144), and Alexa Fluor 647 (ThermoFisher, A-21241).

### Click chemistry

Click-iT Plus EdU Cell Proliferation Kit for Imaging, Alexa Fluor 647 dye (ThermoFisher, C10640) was used to monitor *in situ* DNA synthesis in quiescent primary human fibroblasts. Cells were incubated with EdU for different lengths of time and then fixed in 5% PFA for 15 min. After washing 3× with PBS and 2× with 3% BSA (Sigma-Aldrich, A4378) in PBS, cells were incubated with 0.5% Triton X-100 (Sigma-Aldrich, T8787) in PBS for 20 min at RT. Permeabilization buffer was then removed, and cells were washed 2× with 3% BSA in PBS. After removing wash solution, Click-iT Plus reaction cocktail was added to cells, covered by tinfoil, and incubated on a rocker for 30 min. Reaction cocktail was then removed, and cells were washed 1× with 3% BSA in PBS. Immunofluorescence staining was subsequently carried out to label mitochondria and the total population of mtDNA molecules. Hoechst was then used to stain nuclei for 1 h, followed by 3× washes with PBS.

### Pulse–chase

Quiescent primary human fibroblasts were incubated with 10 *μ*M EdU for 24 h and then washed 3× with DMEM, excluding EdU. Cells were then incubated with 1 mM thymidine (ThermoFisher, A11493.06) to compete with residual EdU. Subsequently, cells were fixed immediately (0 h) to establish a baseline value and at 24, 48, and 96 h to monitor clearance of EdU-labelled mtDNA molecules over time. Following fixation, cells underwent click chemistry, immunofluorescence staining, and super-resolution imaging.

### Fluorescent probes

CellMask Plasma Membrane Stain (ThermoFisher, C10046) was used at a dilution of 1:1000 in live-cell imaging experiments to stain the plasma membrane. SPY620-actin (Spirochrome) was used at 1:1000 dilution to label whole cells during live-cell imaging. Tetramethylrhodamine, methyl ester (TMRM) (Abcam, ab275547) was used at a final concentration of 15 nM to monitor total mitochondrial volume as well as ΔΨ_m_. TMRM remained in imaging media for the duration of experiments. SYBR Green (ThermoFisher, S7563) was used at a dilution of 1:3×10^5^ to label nucleoids during live-cell imaging experiments. SYBR Green was incubated with cells for 10 min before washing 3× with DMEM. Hoechst 33342 (ThermoFisher, 62249) was used at dilution of 1:1000, incubated for 1 h, and washed 3× with PBS following immunostaining to label nuclei.

### Lattice structured illumination microscopy (SIM)

Lattice SIM images were acquired using a Zeiss Elyra 7 lattice SIM system (Carl Zeiss Microscopy) equipped with two sCMOS cameras. Imaging was performed using a 13- or 15-phased imaging protocol across all channels. Fluorophores were excited with 405-, 488-, 561-, and 642-nm lasers, and emissions were collected using an alpha Plan-Apochromat 40×/1.4 or 63×/1.4 Oil DIC objective. Z-stack images were manually set from the bottom to top of cells to ensure whole cell volumes were captured. Raw lattice SIM z-stacks were processed using the standard deconvolution setting in Zen Black (Carl Zeiss Microscopy).

### Airyscan 2 microscopy

Airyscan 2 microscopy was performed using a Zeiss LSM 900 microscope (Carl Zeiss Microscopy). Fluorophores were excited using 405- and 640-nm lasers and imaged with a Plan-Apochromat 40×/1.3 Oil DIC (UV) VIS-IR M27 objective. Raw Airyscan 2 images were processed using the SR deconvolution setting in Zen Blue (Carl Zeiss Microscopy).

### Lattice Llightsheet 7 microscopy

Lattice light-sheet microscopy was conducted using a Zeiss Lattice Lightsheet 7 microscope equipped with a sCMOS camera. Fluorophores were excited using 488-, 561-, and 640-nm lasers and a 13.3×/0.4 water objective. Emissions were detected with a 44.83×/1.0 water objective. Images were acquired at 15-min intervals for 3 h or at 2-h intervals for 24 h. Raw lattice light-sheet images were processed using standard deskewing and deconvolution with strength level 6 in Zen Blue (Carl Zeiss Microscopy).

### Live-cell imaging

Cells were plated in *μ*-Slide 8 well glass bottom imaging dishes (Ibidi, 80827) and incubated overnight and subsequently during live-cell imaging at 37 °C, 5% CO_2_, and 95% relative humidity.

### Image analysis

Z-stacks of whole individual cells were cropped in Fiji (ImageJ) and converted to Imaris (.ims) file types using ImarisFileConverter 10.2.0 (Oxford Instruments). Subsequently, objects in each channel in .ims files were segmented and rendered in 3D using the surfaces function in Imaris 10.2.0 (Oxford Instruments). Segmentation setup included smoothing with surfaces detail of 0.0986 *μ*m and background subtraction (local contrast) of 0.370 *μ*m and 0.1 *μ*m were used for TOMM20 and nucleoid channels, respectively. For whole-cell markers, smoothing with surfaces detail of 1 *μ*m was used. For nucleoid channels, split touching objects (region growing) was used with an intensity-based seed points diameter of 0.1 *μ*m. A surfaces filter was then applied to select objects above ten voxels to eliminate noise. The Imaris machine-learning module was used for segmenting and rendering nuclear and cell-volume markers. In cells lacking a bona fide cell marker, machine learning was used to recognize cytoplasmic background from combined channels as a proxy for cellular boundaries. 3D shape descriptors were used to evaluate mitochondrial morphology, including oblate and prolate ellipticity, indicating disk- and cigar-shaped spheroids, respectively, as well as sphericity, indicating how spherical an object is. Object statistics from Imaris surfaces were subsequently compiled in Microsoft Excel. For 2D analysis, z-stacks were converted to maximum-intensity projections in ImageJ, and mitochondria, nucleoids, and nuclei were subjected to a manual threshold before segmentation of regions of interest (ROIs) and quantification. With the x–y coordinates of nucleoids and the centroid of the nucleus, Euclidean geometry was used to determine the distances of mtEdU and mtDNA molecules from the centre of the nucleus.

### Statistical analysis

Data were subjected to normality tests in GraphPad Prism, and statistical significance was determined using one- or two-tailed Student’s *t*-tests or one-way Analysis of Variance (ANOVA) to evaluate differences between groups, followed by a Tukey’s multiple comparison test. Specific *p*-values are given in figures, and *p* < 0.05 is considered statistically significant. Pearson correlation coefficient (*r*) was used to measure linear correlation between two variables, and the coefficient of determination (*R*^2^) was used to assess the proportion of variance in one variable that is predictable from the other.

### Stochastic birth-death models

All stochastic models, and the necessity driven by the data for each modelling choice, are discussed in full detail in Supplementary Information Sections 1 and 2. All stochastic models (one-population model, preliminary two-population model, two-population model, three-population model) were simulated using the Gillespie algorithm^89^. Every model had a per-capita birth rate of the form *λ*_*b*_ = *μ*_*b*_ + *f*(*N, l*)^65^, where *N* is the nucleoid number, *l* is the mitochondrial volume, *μ*_*b*_ is a constant. *f* is a control term which ensures that the steady-state mtDNA copy number scales linearly with mitochondrial volume, which we assumed to only depend on *l*, not on *V* (cellular volume), due to the conclusions drawn from our conditional independence tests and Granger-causality analysis. We considered three possibilities for the function *f*: *f*(*N, l*) = *c*(*N*_*opt*_ − *N*) (differential control), 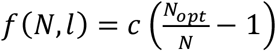 (ratiometric control), 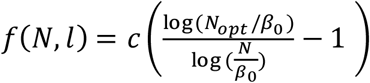 (logarithmic control), as well as one other arising from a hypothetical biological mechanism (see Supplementary Information Section 6.1), selecting logarithmic control due to its heteroskedastic properties (see Supplementary Informat ion Section 6.2). Here, *c* is the strength of the control, and *N*_*opt*_ = *β*_0_ + *β*_1_*l* is the linear relationship between *N* and *I*, with *β*_0_, *β*_1_ constant. All other parameters other than *λ*_*b*_ are constant. The one-, preliminary two-, two-, and three-population models contain parameters (*λ*_*b*_, *μ*_*d*_, *c*), (*λ*_*b*_, *μ*_*d*_, *c, μ*_*r*_), (*λ*_*b*_, *μ*_*d*_, *c, μ*_*r*_, *p*), (*λ*_*b*_, *μ*_*d*_, *c, μ*_*r*_, *p, μ*_*a*_), respectively, along with (*β*_0_, *β*_1_). Steady-state requirements were computed to be *μ*_*b*_ = *μ*_*d*_ for the one- and preliminary two-population models, *μ*_*b*_ = *pμ*_*d*_ for the two-population model, and *μ*_*b*_ = *pμ*_*a*_ for the three-population model, and models were initialized using the nucleoid count observed in cells, with subpopulation proportion set to their computed value under a deterministic treatment (see Supplementary Information Section 2). Existing models in the literature are discussed in Supplementary Information Section 1.5.

### Approximate Bayesian computation

ABC^90^ was used to fit all stochastic models, using assays 1 and 2 for training, and assay 3 for validation (see Supplementary Information Section 4.6 for validation). Every model was fit from 500,000 prior draws with an acceptance rate of 0.1%, leaving 500 posterior draws. We fit separately to two data sets: the *pulse cells* (the data of Fig. 4, as well as the inferred proportion of singly tagged molecules at the first time point, displayed in Fig. 6h) and the *pulse–chase cells* (the data of Fig. 6, as well as the inferred proportion of singly tagged molecules at the first and last time points). Priors for the pulse cells were mostly log-uniform, and steady state was assumed. For the pulse–chase cells, we assumed a change in parameterization in between the pulse and chase phases since steady state is no longer a valid assumption for the chase portion. We assumed that the parameters (*μ*_*b*_, *μ*_*a*_) changed for the three-population model, and (*μ*_*b*_, *μ*_*d*_) for the two-population model, with all other parameters remaining the same for both portions. We also show the fit when (*μ*_*b*_, *μ*_*d*_, *p*) are allowed to vary in the two-population model, showing that the three-population model is still selected even in this case, in Supplementary Information Sections 3, 4. We used a perturbation of the posteriors inferred for the pulse cells as priors for the pulse phase parameters of the pulse–chase cells, while the chase phase parameters were given log-uniform priors. See Supplementary Information Section 3 for full details of the ABC, including priors, summary statistics, distance function, and posteriors for each model.

### Posteriors and posterior predictive plots

Fig. 4b, f, g display estimates of the three-population model, fit using ABC to the data of Fig. 4a, b. Fig. 4b is the result of a single simulation using the estimated posterior mode, computed using a nearest neighbors algorithm, Fig. 4f is a posterior predictive plot using all 500 posterior draws, and Fig. 4g is the posterior distribution of the derived quantity *μ*_*r*_ *f*_*rep*_, where *μ*_*r*_ is the rate at which molecules leave the replicating population (finish replicating), and *f*_*rep*_ is the analytically computed fraction of molecules in the replicating population (computed in Supplementary Information Section 2). Fig. 6j, l, m are posterior predictive plots from the 500 posterior draws of each model, though Fig. 6l only displays the posterior predictive means. Fig. 6o, p are posterior distributions of the derived population proportions and derived mean dwell time (both computed in Supplementary Information Section 2), respectively. Fig. 6k, n display model posteriors, computed by combining all prior simulated draws from every model labelled by the model they came from and setting the rejection rate to only accept 500 total posterior parameter draws. Full details of ABC model selection, posteriors, posterior predictive plots, posterior interpretations, and validation are given in Supplementary Information Section 4. Existing replication rate estimates in the literature are discussed in Supplementary Information Section 4.3.2.

### Causal modelling

Conditional independence tests were implemented using R package “GeneralisedCovarianceMeasure” for GCM^45^, “CondIndTests” for KCIT^46,91^, both on CRAN, and the python package “tigramite” for CMknn^46^. Full table of test statistics and *p*-values are displayed in Supplementary Information Section 7. For linear Granger causality, lag was determined through a 19-cross validation with each fold removing the data from one of the 19 cells, and *p*-values were determined through an *F*-test. Full table of *p*-values is displayed in Supplementary Information Section 7. For neural Granger causality, we used an LSTM (Long Short-Term Memory)^92^ as our network architecture, as this was found to perform better under small data constraints^48^. We chose a context length of 6, and hidden layer size of 5, but ensured our model was robust to different choices (see Supplementary Information Section 7.1.2). ||*W*_*i* → *j*_|| refers to the *l*^2^-norm of the subset of neural network parameters involved in predicting time series *j* from past values of time series *i* (see Supplementary Information Section 7.1.2). All data were normalized to have mean 0 and variance 1 prior to analysis for the conditional independence tests and linear Granger causality. Data were normalized via a min-max normalization for the neural Granger causality due to evidence of better performance in prior studies^93^, along with the fact that we found that the causal networks inferred were more stable.

### Gaussian mixture modelling

Gaussian mixture models were fit using maximum likelihood estimation. Two and three peaks were determined to be better than one based on BIC score (see Supplementary Information Section 3.1.3). Two peaks were fit for all but the first time point in line with our singly and doubly tagged interpretation. Fitting three rather than two peaks for the first time point was done on the following grounds. First, biologically, we expect there to be a third peak due to the presence of replicating nucleoids incorporating more than two strands of EdU from the pulse portion of the experiment. Second, agreeing with this biological reasoning, fitting with two peaks leads to estimates which are inconsistent with the remaining time points (see Supplementary Information Section 3.1.3), whereas three peaks lead to a consistent singly tagged proportion estimate across assays, and consistency across time points (Extended Data Fig. 7). For full details on the Gaussian mixture modelling, see Supplementary Information Section 3.

### LC3 data modelling

The conclusion for the majority of nucleoids degrading through autophagy is illustrated here. If we assume every nucleoid degrades through this pathway, we can estimate that a nucleoid is detectable within an LC3-positive vesicle for 57 ± 16 min, roughly in accord with prior work on rat kidney cells^61^. This arises from the following logic: the rate of nucleoids exiting the vesicles should be equal to the rate of nucleoids entering the vesicles (the global replication rate, if we are assuming steady state, and that every nucleoid degrades through this pathway). Hence, the per-capita rate of nucleoids exiting the vesicles should be the global replication rate divided by the number of mtDNA in the vesicle, and the average time a nucleoid spends in a vesicle should hence be the inverse of this quantity: 57 ± 16 min. Assuming a lower percentage of nucleoids degrades through this pathway skews the estimate to be unreasonably large; for instance, if only 50% of nucleoids are degraded by LC3-positive routes, we estimate degradation takes a less credible estimate of ~2 hours. Full reasoning and modelling is given in Supplementary Information Section 5.3. The test of Fig. 5d is as follows. Let *V, V*_*L*_, *l, N, N*_*L*_ denote the cellular volume, total LC3 volume, mitochondrial volume, nucleoid number, and number of nucleoids enveloped by LC3, of a particular cell, respectively. Assuming that LC3 is distributed uniformly throughout the cell, the volume of LC3 overlapping with the mitochondrial network is given by 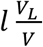, which as a fraction of the mitochondrial volume is given by 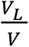. Hence, assuming the number of nucleoids is distributed uniformly across the mitochondrial network, the number of nucleoids randomly overlapping with LC3 is given by 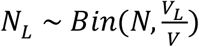. Computing the mean of this quantity over every observed cell and running a Monte-Carlo simulation allows us to generate a null distribution, which we then compare to the true observed mean.

### Other mathematical modelling

Primary mouse macrophages were fit using a deterministic version of the two-population model, assumed to be in steady state for the non-LPS stimulated cells and out of steady state for LPS stimulated cells, using nonlinear least squares (see Supplementary Information Section 5.1). For non-LPS stimulated cells, we found that the data were consistent with the two-population model with *μ*_*b*_, *μ*_*d*_ = 0, but nonzero *μ*_*r*_, *p*, allowing the replicating subpopulation to deplete while disallowing any new birth and death events. Ethidium bromide degradation data were fit using a deterministic version of the three-population model (two-step degradation), and the two-population model (one-step degradation), assuming the replicating subpopulation was empty, using nonlinear least squares (see Supplementary Information Section 5.2). All quoted errors (i.e., x ± y) refer to standard deviations of the posterior for Bayesian inferences, and standard errors for frequentist inferences.

## Extended Data Figures

**Extended Data Figure 1:**
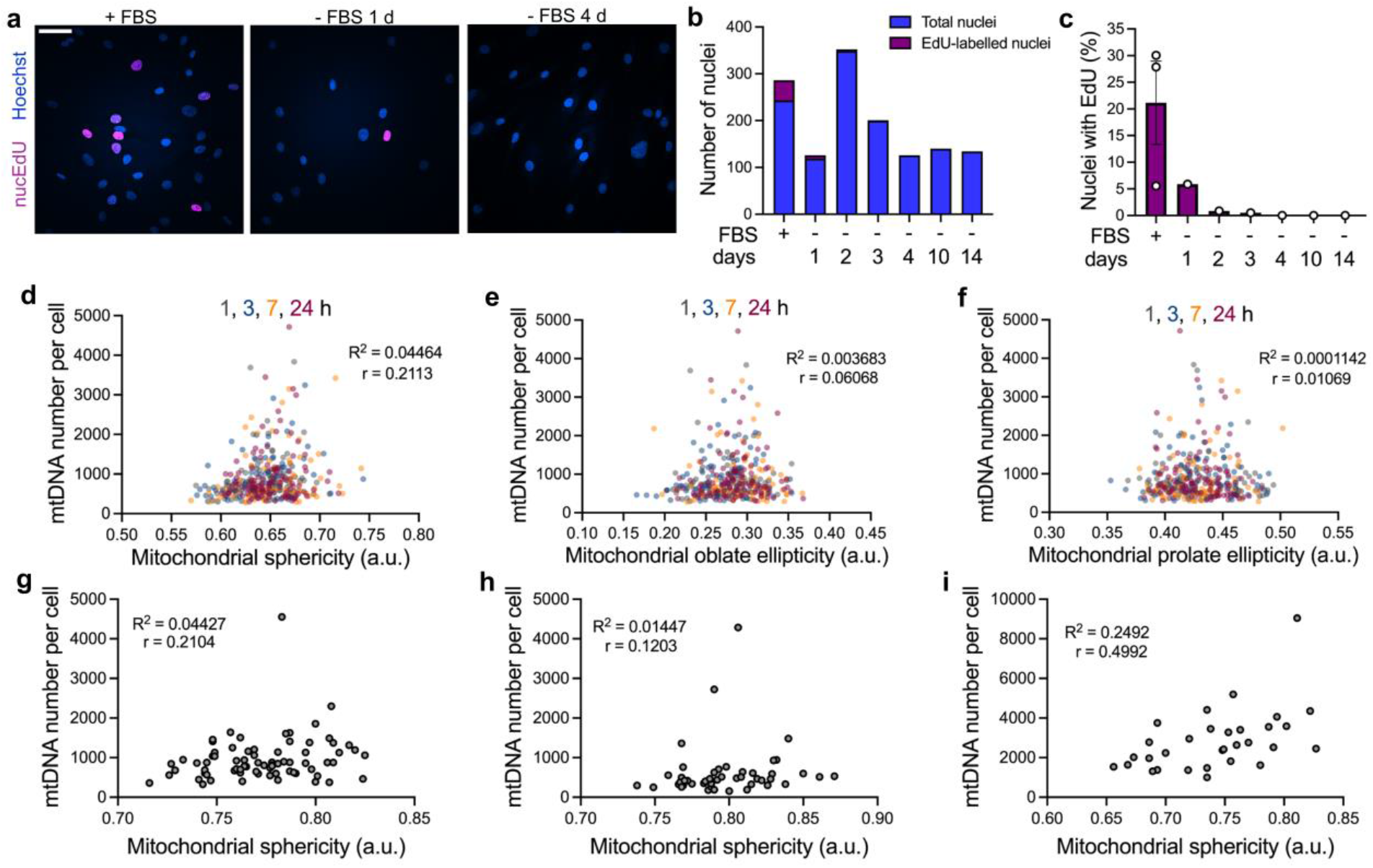
Approach for inducing quiescence in primary human fibroblasts. **a**, Representative Airyscan 2 images of fixed primary human fibroblasts with and without foetal bovine serum (FBS). Incorporation of EdU (magenta) into nuclear DNA (blue) indicates cells in a proliferative state. Scale bar = 50 *μ*m. **b, c**, Withdrawal of FBS induces quiescence after 4 days. Data reflect analysis of 1,798 (a–c) cells from ≥ 3 independent experiments. **d**–**f**, In quiescent primary human fibroblasts, mtDNA CN does not correlate with mitochondrial morphology. Data reflect 3D analysis of 381 cells from 3 independent experiments. Statistical tests: Pearson correlation coefficient (*r*) and coefficient of determination (*R*^2^) are shown in panels. **g–i**, MtDNA CN is not correlated with mitochondrial morphology in HeLa (g), COS-7 (h) cells, or primary murine hepatocytes (i). Data reflect 3D analysis of 73 (g) and 48 (h) cells from 3 independent experiments; and 31 cells (i) from ≥ 3 biological replicates. Statistical tests: Pearson correlation coefficient (*r*) and coefficient of determination (*R*^2^) are shown in panels.

**Extended Data Figure 2:**
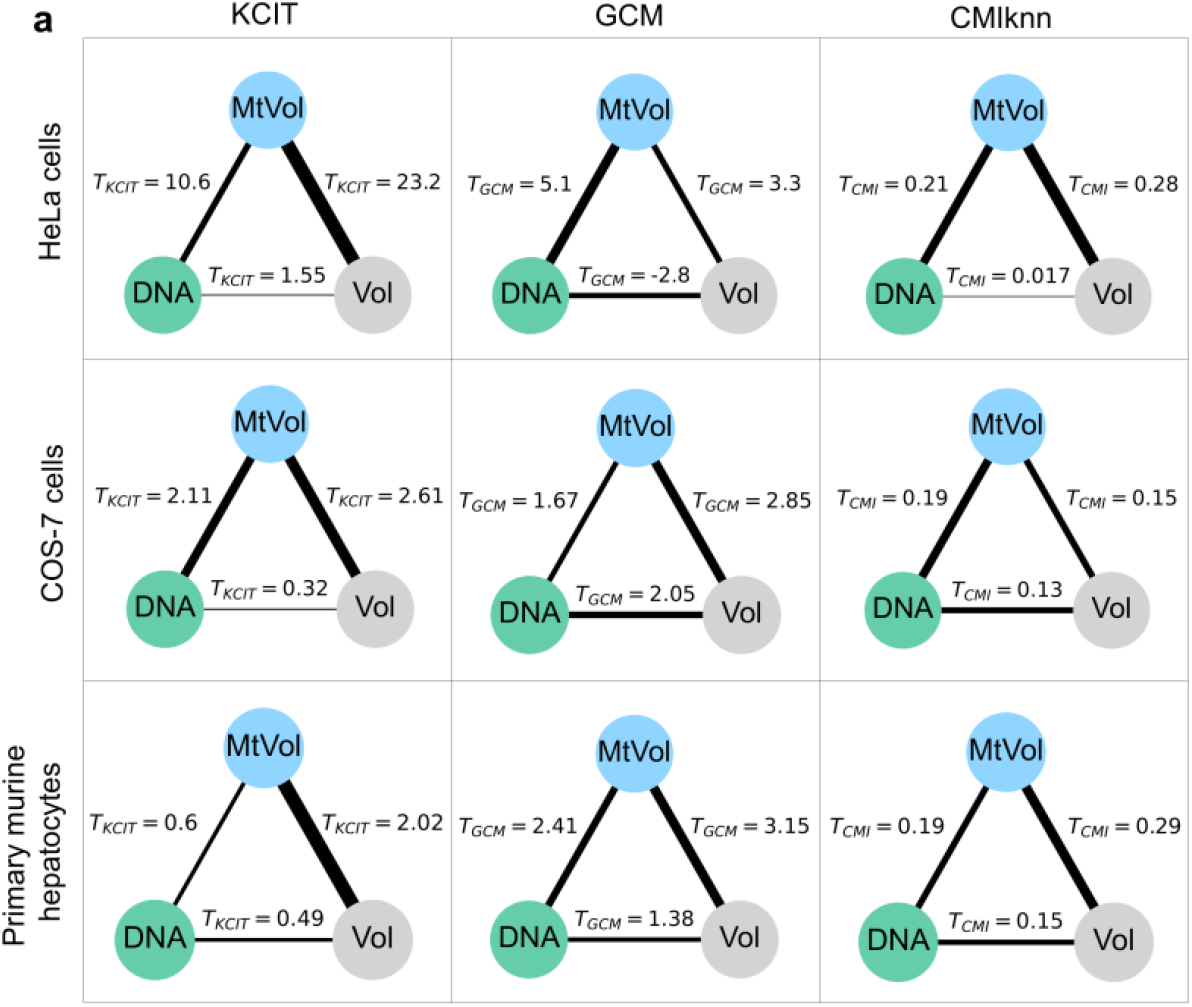
Conditional independence tests reveal a weaker connection between nucleoid number and cellular volume than other connections in a variety of cell types. **a**, Test statistic for conditional independence between nucleoid number and cellular volume is smaller in magnitude than that between nucleoid number and mitochondrial network volume, and between mitochondrial network and cellular volume for all cell types and tests other than GCM for the COS-7 cells.

**Extended Data Figure 3:**
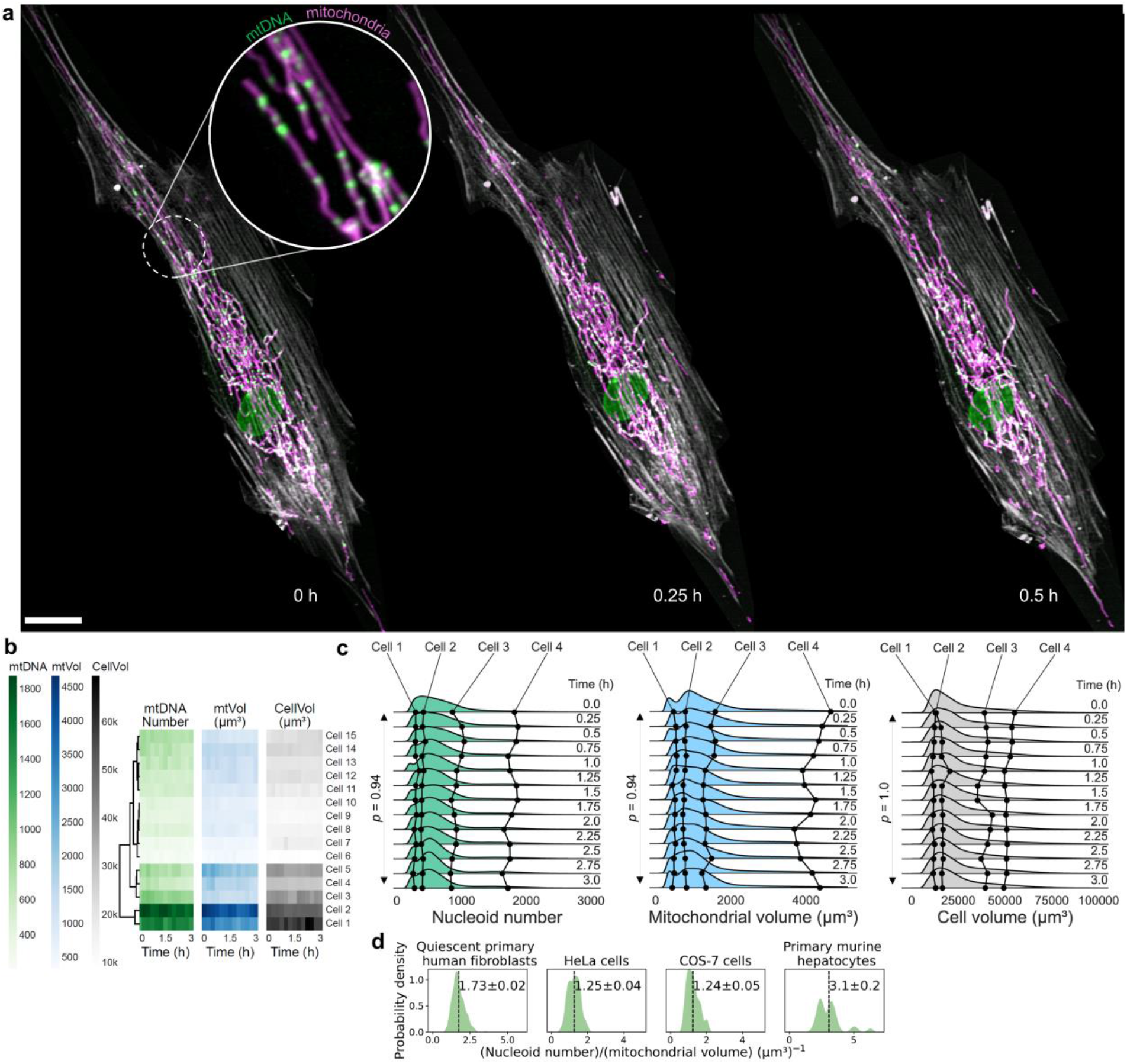
4D imaging of quiescent primary human fibroblasts at short time intervals indicates maintenance of steady state initially after cell-cycle arrest. **a**, Representative lattice light-sheet maximum-intensity projections of individual quiescent primary human fibroblast, showing cell boundaries (light gray), mitochondria (magenta), and mtDNA (green). Scale bar = 20 µm. **b**, Heatmaps of nucleoid number, mitochondrial network volume, and cellular volume for each cell measured in the 4D lattice light-sheet imaging, with dendrogram. **c**, Distribution of nucleoid number, mitochondrial network volume, and cellular volume over time from 4D lattice light-sheet imaging data, with plots from individual cells overlaid, demonstrating minimal changes in cellular volume, total mitochondrial volume, and nucleoid number. Data reflect 4D analysis of 15 cells from 3 independent experiments. **d**, Nucleoid density in a variety of mammalian cell types. Data reflect 3D lattice SIM analysis of 381 quiescent primary human fibroblasts, 78 HeLa and 48 COS-7 cells from 3 independent experiments, and 31 primary murine hepatocytes from ≥ 3 biological replicates.

**Extended Data Figure 4:**
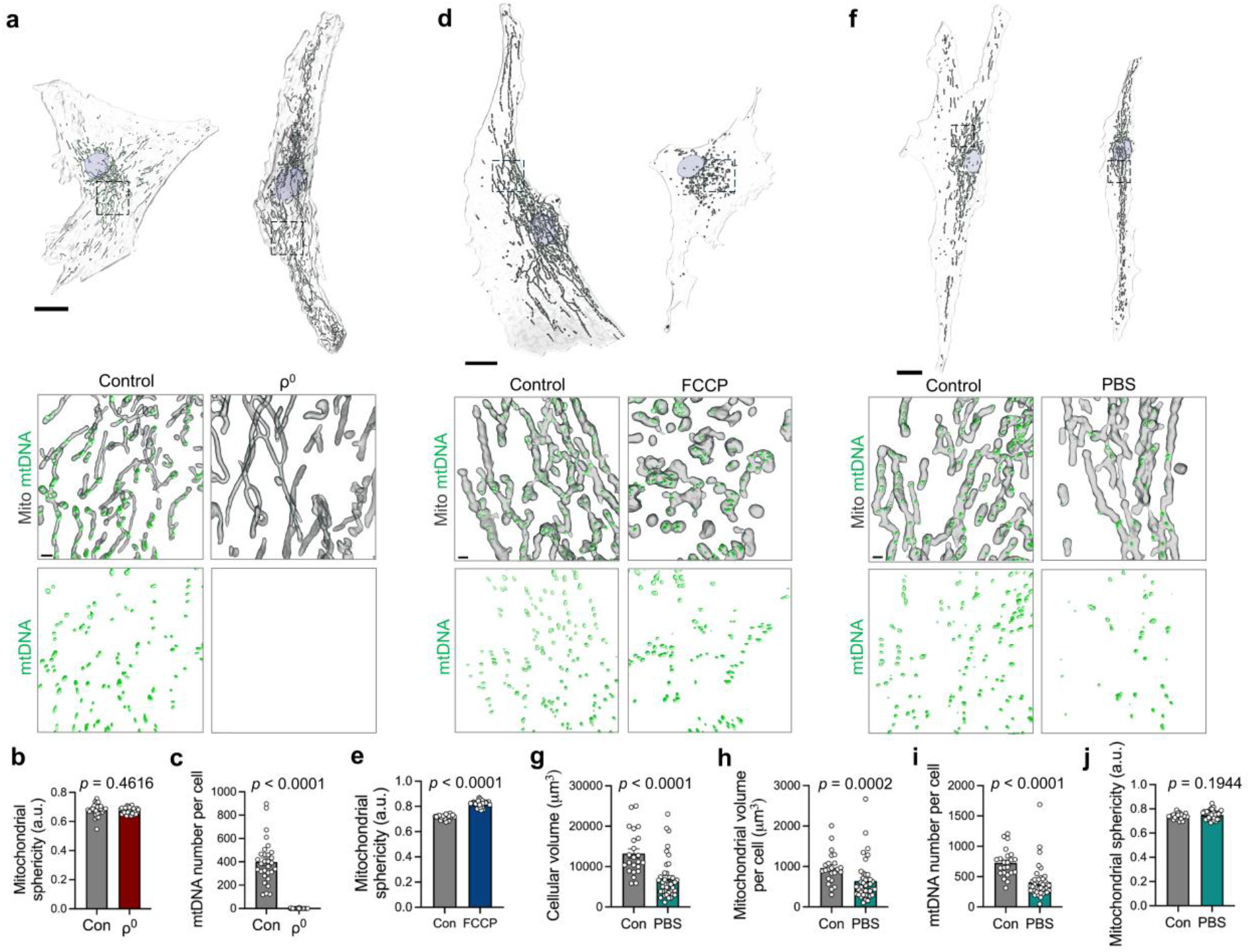
Cellular and mitochondrial network volumes, together with mtDNA copy number, are regulated by mitochondrial bioenergetics and nutrient status. **a**, Representative live-cell, lattice SIM renderings of quiescent primary human fibroblasts, with and without mtDNA. *Top:* Renderings of whole cells (transparent), showing nuclei (blue), mitochondria (gray), and mtDNA (green). Scale bar = 20 µm. *Bottom:* insets of cells with details of mitochondria and mtDNA. Scale bar = 1 µm. **b, c**, depletion of mtDNA (ρ^0^) does not affect mitochondrial morphology in quiescent primary human fibroblasts. **d**, Representative lattice SIM renderings of quiescent primary human fibroblasts, treated with vehicle (DMSO) versus 10 µM FCCP for 24 h, prior to fixation and immunostaining. *Top:* Renderings of whole cells (transparent), showing nuclei (blue), mitochondria (gray), and mtDNA (green). Scale bar = 20 µm. *Bottom:* insets of cells with details of mitochondria and mtDNA. Scale bar = 1 µm. **e**, Treatment with FCCP induces mitochondrial fragmentation. **f**, Representative lattice SIM renderings of quiescent primary human fibroblasts, grown in DMEM without FBS versus phosphate buffered saline (PBS) for 96 h, prior to fixation and immunostaining. *Top:* Renderings of whole cells (transparent), showing nuclei (blue), mitochondria (gray), and mtDNA (green). Scale bar = 20 µm. *Bottom:* insets of cells with details of mitochondria and mtDNA. Scale bar = 1 µm. **g–j**, Starvation induces quiescent primary human fibroblasts to shrink (g) and decreases total mitochondrial volume (h) and mtDNA copy number (i), while mitochondrial morphology (j) was not significantly affected. Data reflect 3D analysis of 67 (a– c) 54 (d, e), and 63 (f–j) cells from 3 independent experiments, respectively. *P*-values shown in panels were determined using two-tailed Student’s *t*-tests. Data are mean ± s.e.m.

**Extended Data Figure 5:**
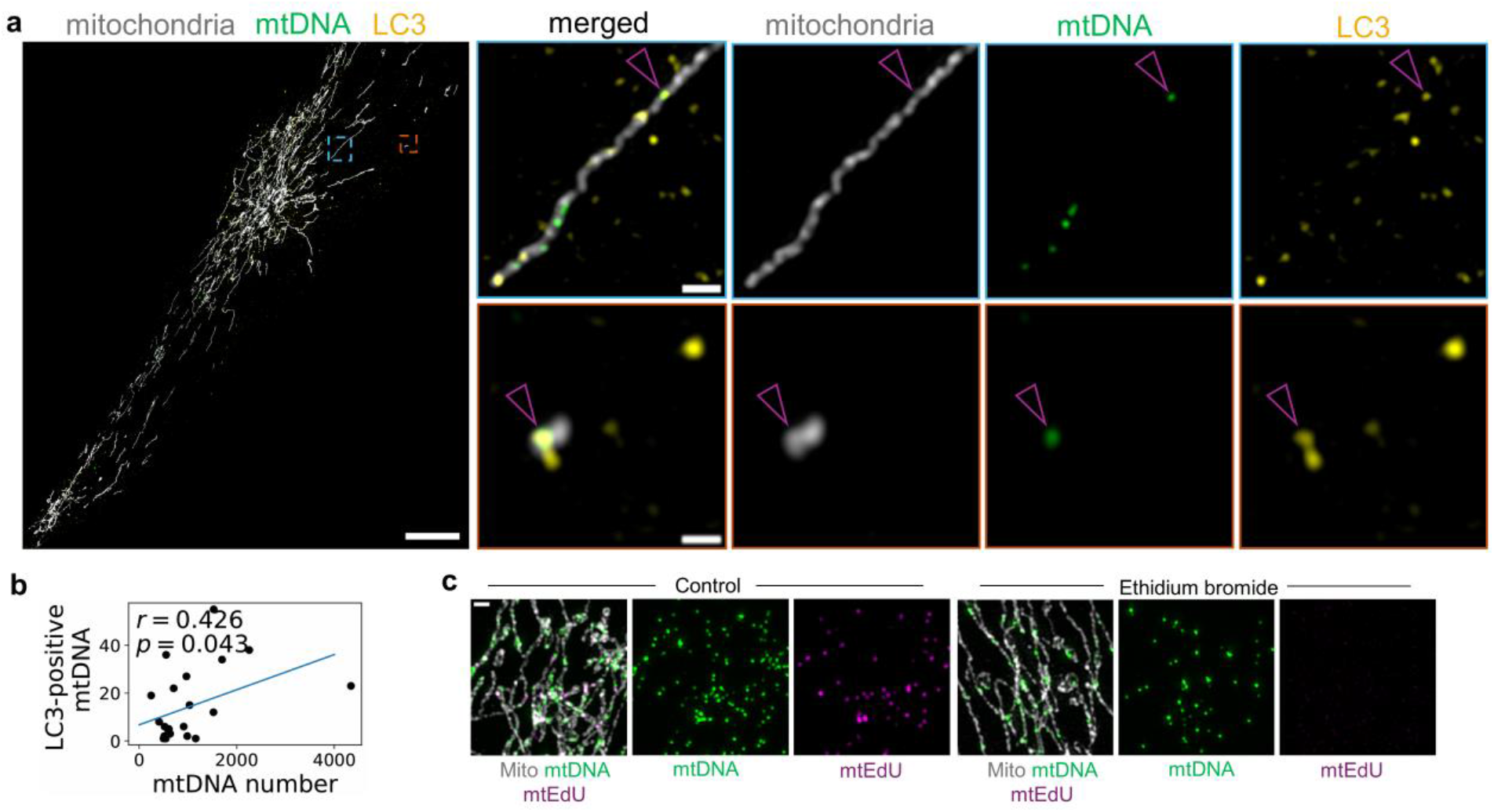
3D super-resolution imaging of LC3-positive nucleoids in quiescent primary human fibroblasts. **a**, Representative 3D lattice SIM maximum-intensity projections of quiescent primary human fibroblasts, immunostained with anti-TOMM20 labelling mitochondria (gray), anti-DNA labelling the whole population of mtDNA (green), and anti-LC3 labelling early-to-intermediate autophagosomes (yellow). Arrowheads show mtDNA colocalization with LC3. Whole cell (left) scale bar = 20 µm; top inset scale bar = 1 µm; bottom inset = 0.5 µm. **b**, LC3-positive mtDNA number scales with total mtDNA number per cell. **c**, Representative 3D lattice SIM maximum-intensity projections of quiescent primary human fibroblasts treated with 500 nM ethidium bromide for 28 h, in the presence of 10 µM EdU, prior to fixation and immunostaining with anti-TOMM20 (gray) and anti-DNA (green), together with EdU labelling replicating nucleoids (magenta). Scale bar = 1 µm. EdU was added 1 h after commencing ethidium bromide treatment. Data reflect 3D analysis of 23 (a, b) cells from 3 independent experiments. Statistical tests: Pearson correlation coefficient (*r*) with *p*-value from two-tailed Student’s *t*-test are shown (b).

**Extended Data Figure 6:**
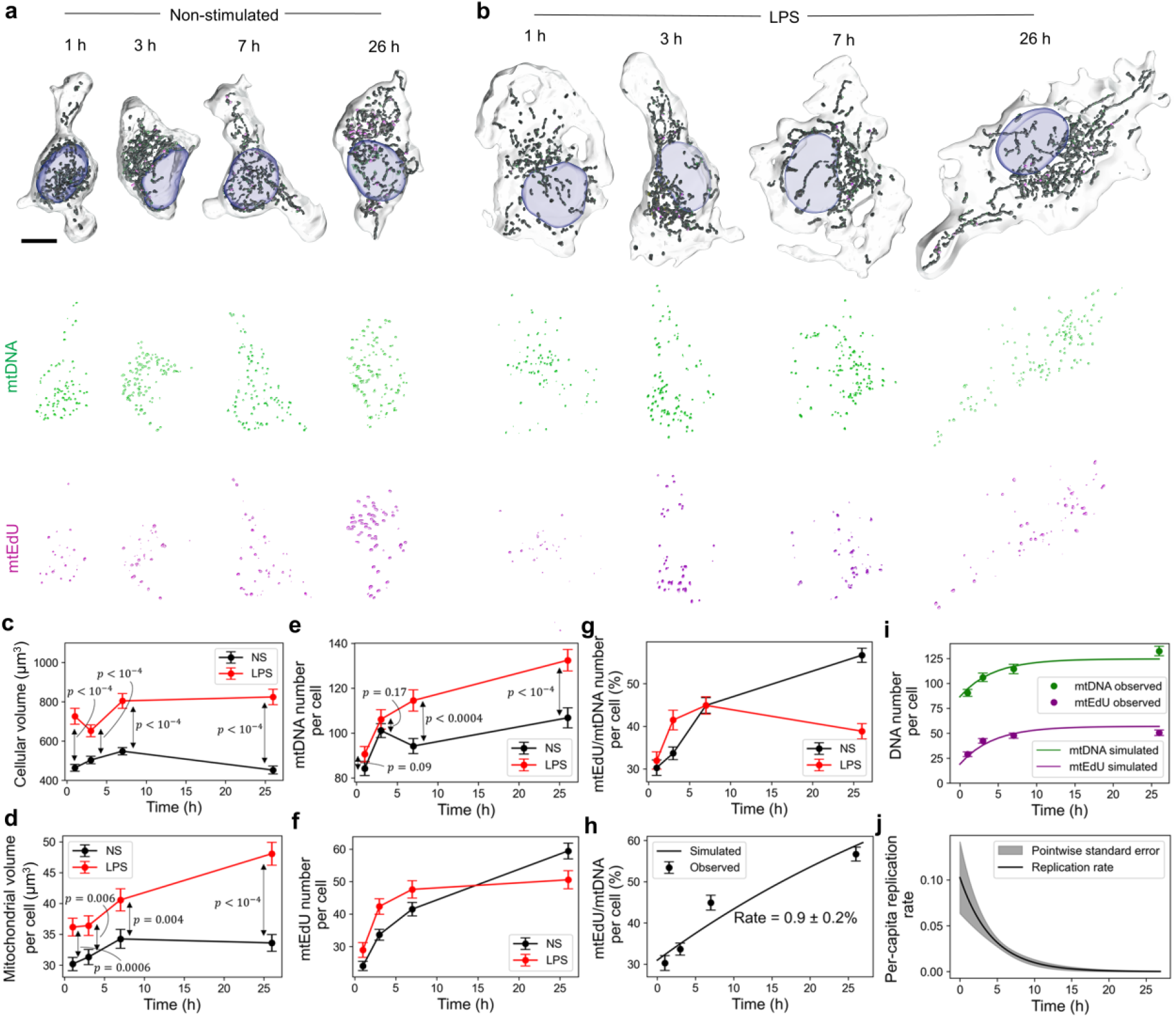
Lipopolysaccharide (LPS) stimulation of primary murine macrophages increases cellular and mitochondrial volumes together with mtDNA copy number. **a, b**, Representative lattice SIM renderings of non-stimulated (NS) (a) and LPS-stimulated (b) primary murine macrophages, treated with 50 µM EdU for 1, 3, 7, and 26 h, prior to fixation and immunostaining. *Top:* Renderings of whole cells (transparent), showing nuclei (blue), mitochondria (dark gray), mtDNA (green), and mtEdU (magenta). Scale bar = 5 µm. *Bottom:* mtDNA and mtEdU at different time points. **c, d**, Cellular (c) and total mitochondrial (d) volumes increase upon LPS stimulation. **e**, MtDNA levels increase following LPS stimulation. **f, g**, MtEdU levels increase in both LPS-stimulated and NS cells. **h**, Deterministic two-population model fit indicates NS cells replicate at a rate of 0.9 ± 0.2% per h. **i**, Deterministic two-population model fit to LPS-stimulated cells. **j**, LPS-stimulated cells are consistent with a deterministic two-population model with an mtDNA replication rate that increases initially to 10 ± 3.9% per h after LPS stimulation and decreases exponentially after 26 h. Data are mean ± s.e.m. of 3 biological replicates, reflecting 3D analysis of 284 (NS) and 223 (LPS) cells. *P*-values shown in panels were determined using one-tailed Student’s *t*-tests.

**Extended Data Figure 7:**
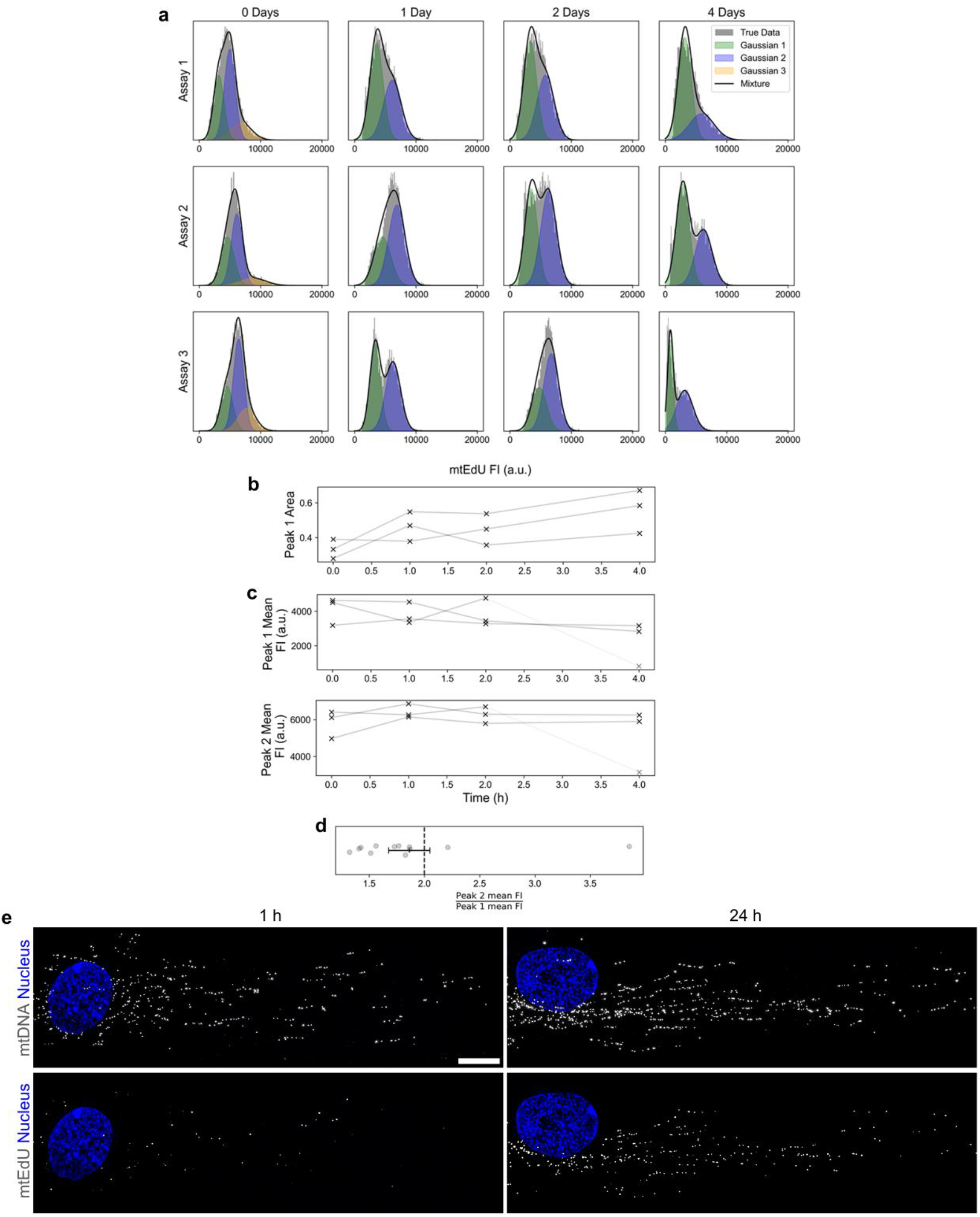
Gaussian mixture models infer two major FI peaks, corresponding to singly and doubly tagged mtEdU, across all assays and time points. **a**, Frequency distribution of FIs, with Gaussian mixture model fits, for every assay and time point of the chase portion of the experiment. Two prominent peaks are seen, corresponding to singly and doubly tagged mtEdU, as we**ll** as a third minor peak in the first time point for replicating molecules consisting of nucleoids with more than two tagged strands. **b**, Area of the first Gaussian is consistent across assays and increases over time, interpreted as more singly tagged mtEdU emerging and less doubly tagged, indicating active replication. **c**, Mean FI of first (*top*) and second (*bottom*) Gaussians are stable over time and consistent across assays, other than the anomalous 4-day assay 3, indicating that the average FI of mtEdU which is singly or doubly tagged is not changing over time. **d**, The mean FI of the second peak is double the mean FI of the first, across al**l** assays and time points, indicating that doubly tagged mtEdU is twice the FI of singly tagged mtEdU. Horizontal bar indicates mean ± s.e.m. Data reflect 3D analysis of 265 cells from 3 independent experiments, including 106,176 mtEdU puncta. **e**, Representative 3D lattice SIM maximum-intensity projections of quiescent primary human fibroblasts treated with 10 µM EdU for different times, prior to fixation and immunostaining. Nuclei (blue), mtDNA (white), and mtEdU (white). Scale bar = 10 µm. Newly replicated mtDNA molecules are closer to the nucleus than the general population of mtDNA (1 h) and, over time (24 h), are distributed to the cellular periphery. Data reflect maximum-intensity-projection analysis of 381 cells from 3 independent experiments.

