## Supplementary material for "Single-nucleoid imaging in whole cells defines the dynamics of the mtDNA life cycle": Mathematical Modelling Supplement: mtDNA_Replication_Dynamics_Mathematical_Supplement_BioRxiv.pdf

**8 Discussion** **44**

**Appendix A Glossary of Key Terms and Symbols** **48**

**Appendix B Inhibition Model Equilibrium** **49**

**Appendix C Colocalisation Error** **50**

**Appendix D Stochastic Systems** **51**

Code for all mathematical modelling is available at https://github.com/Wendaroo/mtDNA-Replication-Dynamics.git.

### 1 Data Insights and Modelling Choices

The mathematical models we construct are at their heart informed by aspects of the data we see. We have five nonproliferative fibroblast datasets which we draw information from:

- **Pulse data**. Three assays consisting in total of 382 cells. Each cell was measured at a time point, either 1, 3, 7, or 24 hours, its measured quantities being mitochondrial volume, cell volume, nucleoid number, and mtEdU number, with EdU having been inserted into the system at 0 hours.
- **Pulse-chase data**. Three assays consisting in total of 265 cells, having been exposed to EdU for 24 hours like the pulse data (which we will refer to as the 'pulse portion' of the pulse-chase data), but at this 24 hour mark, all EdU is removed from the system (which we refer to as the 'chase portion' of the pulse-chase data. Each cell was designated a time point, either 24, 48, 72, 120 hours (i.e., 0, 1, 2, or 4 days following EdU removal), at which point its mitochondrial volume, cell volume, nucleoid number, and mtEdU number were measured.
- **EdU intensity data**. The intensity of every EdU tagged nucleoid from the pulse-chase data was measured, for each time point.
- **4D Imaging data**. 19 individual cells were tracked for 24 hours, with 2 hour increments between datapoints. Mitochondrial volume, cell volume, and nucleoid number were measured.
- **Degradation data**. Replication was halted in a total of 472 cells through the use of ethidium bromide. Each cell was measured at one of the following time points: 0, 1, 3, 7, 24, 48, 120, 240, 360, 480 hours, its measured quantities being mitochondrial volume, cell volume, and nucleoid number.

This section is structured to illustrate how our models grew in complexity to account for an increasing number of data aspects. For a more in-depth mathematical analysis of each model, see Section 2. For the remainder of this document, let $N, l, V$ denote the nucleoid number, mitochondrial volume, and cell volume of a general cell, respectively.

#### 1.1 Linear Equilibrium and Copy Number Control

The starting point for our modelling is a stochastic model in which we have a population of nucleoids each subject to two Poisson point processes: a birth process with rate $\lambda_b$, and a death process with rate $\lambda_d$. If a birth event occurs, the nucleoid doubles, while if a death event occurs, the nucleoid is removed from the system. A-priori, we do not know if $\lambda_b, \lambda_d$ are constant or functions of the cellular state. On this point, our first insight comes from the degradation data, in which we see a clear exponential-like decay when replication is halted, or in other words, when $\lambda_b \equiv 0$ (Supplementary Fig. 1a). This behaviour is consistent with what we would expect under a constant per capita degradation rate, so we may assume $\lambda_d \equiv \mu_d$ is constant. The fact that the decay is not precisely exponential will later be accounted for with the three-population model (see Section 5.2), still with a constant degradation rate.

Next, main text Figures 3d, 4c, and Extended Data Fig. 3 clearly demonstrate that in the pulse and 4D-imaging data, nucleoid number stays approximately constant over time, meaning that the cells maintain copy number control. This necessitates a birth rate which is, on average, equal to the death rate: $\langle \lambda_b \rangle = \mu_b$, with $\mu_b = \mu_d$, at least for this simple model with only one population of nucleoids (Supplementary Fig. 1b). For more complex models, we will need different conditions on $\mu_b$ to maintain control (see Section 2), and we will define $\mu_b$ more carefully. Rather than each cell having the same global birth rate, we see directly from the data that each cell has the same *per capita* birth rate. Specifically, we see in the data that tagged nucleoid number scales linearly with mitochondrial volume (main text Figure 4b), and hence also with total nucleoid number (as established in main text Figure 2), from which it follows that the global replication rate must be directly proportional to the nucleoid number, or constant per-capita. This justifies our choice of $\lambda_b, \lambda_d$ being per capita rates rather than global rates.

Next, Fig. 2 of the main text demonstrates that the cell seeks to maintain an optimal copy number $N_{opt}$ which scales linearly with mitochondrial and cellular volume. This fact necessitates a non-constant per-capita birth rate that varies based on the cellular state. For instance, if a cell is overpopulated with nucleoids, it must be the case that the birth rate drops to push the cell back to its linear equilibrium. Likewise, if a cell is underpopulated, the birth rate must rise, assuming that the degradation rate is always constant per-capita. We are thus led to adopt a functional form for $\lambda_b$ which satisfies[1,2] (Supplementary Fig. 1c):

$$\lambda_b(N, l, V) = \mu_b + f(N, l, V), \tag{1}$$

where:

$$\begin{cases} f \geq 0, & N < N_{opt} \\ f = 0 & N = N_{opt} \\ f \leq 0 & N > N_{opt}. \end{cases} \tag{2}$$

With the addition of the feedback term $f$, we ensure that the birth rate $\lambda_b$ is larger than the death rate $\mu_d$ (recall that $\mu_b = \mu_d$ for now) when the cell is underpopulated ($N < N_{opt}$), and smaller than the death rate when the cell is overpopulated ($N > N_{opt}$). Finally, the results of the conditional independence testing, Granger and neural Granger causality (main text Fig. 3a, 3e, 3g), suggest that $N$ should not be a direct function of $V$, but rather be a function of $l$, which might in turn be a function of $V$. This motivates the simplification of dropping the dependence on $V$ in $\lambda_b$: $\lambda_b = \mu_b + f(N, l)$, and we may write $N_{opt}$ as being linear with respect only to $l$, not $V$:

$$N_{opt} = \beta_0 + \beta_1 l, \tag{3}$$

for some constants $\beta_0, \beta_1$ (Supplementary Fig. 1d). There are many choices of functional forms $f$ which satisfy Equation 2. In our analysis, we consider the following three:

$$f(N, l) = c(N_{opt}(l) - N) \text{ (Differential control)} \tag{4a}$$

$$f(N, l) = c\left(\frac{N_{opt}(l)}{N} - 1\right) \text{ (Ratiometric control)} \tag{4b}$$

$$f(N, l) = c\left(\frac{\log(N_{opt}(l)/\beta_0)}{\log(N/\beta_0)} - 1\right), \text{ (Logarithmic control)} \tag{4c}$$

where $c$ is some control strength parameter, along with a further hypothetical mechanism arising from a more microscopic model (Section 6). With any of these mechanisms, we implicitly enforce the condition that $\lambda_b \geq 0$ by simulating based on $\max(0, \lambda_b)$ to ensure that the birth rate is non-negative. We select logarithmic birth out of these birth rates to use in our models, as we found that this control law enforces similar heteroscedasticity in the simulated cells as in the real cells. For a detailed comparison and discussion of such birth rates, see Section 6.

It should be noted that in many bioenergetic and physiological situations linearity is the exception, for instance, mammalian basal metabolic rate scales not linearly to body mass ($M$), but proportional to $M^{2/3}$. To ensure our assumption of linearity between $N$ and $l$ is sound therefore, we used nonlinear least squares to fit constants $b_0, b_1, \alpha$ to the equation $N = b_0 + b_1 l^\alpha$ using all assays of the pulse data, obtaining estimate $\hat{\alpha} = 0.9 \pm 0.04$. Given such a small discrepancy from 1, model parsimony strongly suggests fitting a simple linear model.

These insights give rise to our first real model, whereby we have a per capita birth rate $\lambda_b(N, l)$ of the form discussed above (Equation 4c), and a constant per capita death rate $\mu_d$, both modulating Poisson processes acting on a population of nucleoids. We term this the *one-population model* (Supplementary Fig. 2a).

#### 1.2 A Replicating Subpopulation

An inspection of the pulse data at the 1 hour time point (Supplementary Fig. 2b) reveals that there is an unusually rapid accumulation of EdU between 0 and 1 hour before the accumulation slows significantly. To explain this accumulation of EdU, we make the intuitive hypothesis that at time $t = 0$ of the pulse experiment, some fraction of nucleoids in each cell are already mid-replication. This replicating subpopulation will begin accumulating EdU as soon as it is added to the system, leading to this initial rapid acculation we see in the pulse data.

This leads us to update our one-population model into the *preliminary two-population model* (Supplementary Fig. 2c), in which we separate nucleoids into two populations: replicating nucleoids and nonreplicating nucleoids. In this model, each nonreplicating nucleoid can undergo a birth event with rate $\lambda_b$, which does not cause it to double (as in the one-population model), but to move into the replicating subpopulation. We introduce a new 'replication-termination' Poisson process with constant rate $\mu_r$ which each replicating nucleoid undergoes, where an event causes a replicating nucleoid to double, and both daughter nucleoids to move to the nonreplicating population once more. Like in the one-population model, the nonreplicating molecules can die with constant rate $\mu_d$. In effect, this model extents the one-population model from a panmictic pool to a pool with relative compartmentalisation. Enforcing that the replicating subpopulation becomes EdU tagged as soon as EdU is added to the system leads to the same behavior in the models as in the data.

#### 1.3 Newly Replicated Nucleoids are Likely to Replicate Again

We now use the EdU intensity data to draw further insights. In the presence of EdU, when an untagged mtDNA replicates once, its two mtDNA strands split, and both gather an EdU tagged strand, creating two singly tagged molecules. When a singly tagged molecule replicates, its two strands split, and both its tagged strand and its untagged strand will gain a new tagged strand, resulting in one double tagged molecule, and one singly tagged molecule being created. Likewise, a doubly tagged molecule replicating will create two doubly tagged molecules. It follows that a mtDNA must replicate twice to become doubly tagged, and that there are two types of mtEdU: singly and doubly tagged.

Extended Data Fig. 7 demonstrates how we can infer the proportions of mtEdU which are singly and doubly tagged in all assays and time points using Gaussian mixture modelling, by interpreting the first peak as the distribution of singly tagged molecules, and the second peak as the distribution of doubly tagged molecules. We find that for the pulse-chase data, in between the pulse and chase phases, a mean of 33 $\pm$ 3% of mtEdU were singly tagged across the three assays, meaning the the majority of mtEdU were doubly tagged, not singly tagged (Supplementary Fig. 2d). Since a doubly tagged mtEdU must have replicated twice, this data suggests that recently replicated molecules have a tendency to begin replicating again.

This observation necessitates another model update. We allow, with some probability $q = 1 - p$, that if a replication-termination event occurs with rate $\mu_r$, one of its daughter nucleoids remains in the replicating subpopulation, while the other moves to the nonreplicating subpopulation. With probability $p$, both daughter nucleoids move to the replicating subpopulation. In other words, with probability $q$, a molecule which has just finished replicating will begin replicating one of its daughters again. Here, $p$ can be interpreted as the probability the replication machinery diffuses away from the nucleoid it has just replicated, rather than beginning to replicate it again. We term this the *two-population model* (Supplementary Fig. 2e). With this new model upgrade, in order to keep copy number control, it is no longer the case that $\mu_b = \mu_d$, but rather $\mu_b = p\mu_d$. Details are given in Section 2.

#### 1.4 Newly Replicated Nucleoids are Less Likely to Degrade

We now use the pulse-chase data to draw our final insight. In the pulse-chase data, the nucleoid number is dropping, meaning that the cells are no longer in equilibrium, and the birth rate must be lower than the death rate on average. We are exploiting the fact that the cells are beginning to starve after 24 hours in an FBS free environment. It is notable, however, that the mtEdU number does not drop as quickly as the nucleoid number, particularly between the third and fourth time points (Supplementary Fig. 2f). This leads us to the hypotheses that EdU tagged molecules are less likely to degrade than untagged molecules. Since EdU does not fundamentally alter the chemistry of mtDNA, this must imply that newly replicated molecules are less likely to degrade.

To account for the hypothesis that newly replicated molecules are less likely to degrade, we upgrade our two-population model into the *three-population model* (Supplementary Fig. 2g). We introduce a third population of newly replicated molecules which are not able to degrade, with some rate of 'ageing' $\mu_a$ from the newly replicated population to the 'old' population, which is able to degrade. More details regarding this model are given in Section 2.

Further evidence for this three-population model is given in the degradation data. In the one- and two-population models, if we halt replication, all nucleoids become nonreplicative, and only the death Poisson process remains, which would lead to an exponential decay in the nucleoid number $N(t) = N_0 e^{-\mu_b t}$. In the three-population model, the nucleoid number drop would not be precisely exponential, but would have a more complex form (see Section 5.2). This more complex form comes about since the old population would rapidly degrade, while the young population would need some time to age before degrading, leading to a steeper nucleoid number slope initially before becoming more shallow, matching what we observe in the data.

This section has served as motivation necessitated by the data for the construction of our stochastic models. To confirm our hypotheses, we will need to rigorously formulate and fit these models to our data, and perform a model selection; the focus of Sections 2, 3, and 4.

#### 1.5 MtDNA Models in the Literature

Numerous mtDNA proliferation models exist in the literature. Hoitzing et al. give an overview and analytic descriptions of many models[1], and Johnston and Jones discuss numerous control mechanisms satisfying Equation 2[2]. Insalata et al. extend these models in a spatial manner to account for the proliferation of mutations without a replicative advantage[3]. These models all work within a one-population context however, where every mtDNA molecule has identical replication and degradation properties, a feature which our models must necessarily extend to account for our observed data.

Evidence for models with more than one subpopulation also exists in the literature. Isaac et al. observe through sequencing that nucleoids exist in either a compactified, transcriptionally inaccessible state, or in an accessible state, and provide evidence suggesting that those in an inaccessible state tend to undergo less replication[4]. They find that the majority of nucleoids lie in this inaccessible, nonreplicative state, consistent with our inferred 'old' subpopulation size (main text Fig. 6o), although they do not attempt to fit a quantitative mathematical model. Brüser et al. posit a quantitative two-population model with a 'fast' population (which both replicates and degrades) and 'slow/inactive' population (which replicates and degrades at a much slower rate), motivated by EdU incorporation dynamics that could not be explained through a one-population model[5]. They also provide evidence that this fast population is preferentially transcribed, in agreement with Isaac et al[4]. Although this model has the advantage of quantitative fit to data, and concludes that the majority of nucleoids are 'inactive', agreeing with our modelling (see main text Fig. 6o), critically, the models are fit to proliferating cells. They endeavour to account for this by assuming that the nucleoid count rises over time as the cell progresses through its cycle, meaning their model has no equilibrium, however, they do not attempt to model the effect of the mitosis event itself, and the sharing of nucleoids between the two daughter cells therein. Confounding effects from the cell cycle therefore urge us to interpret any modelling results with care. Our modelling focusses on non-proliferative cells, and therefore avoids these constraints. Two critical pieces of information also missing from this model that we include in ours are the evidence of repeated replication from our EdU intensity data, and evidence of preferential degradation of old molecules from our pulse-chase data. This model is also deterministic, and so is not able to select between different control mechanisms[2] (Equation 2), for example, which our stochastic model can.

Cote-L'Heureu et al. also posit a two-population model, motivated by the observation that certain transversion mutations do not accumulate with age in mice[6,7]. This model introduces a 'stem' population (analagous to our replicating and young populations), which actively proliferates, and a 'worker' population (analagous to our old population), which is transcribed and not actively replicating. They do not quantitatively fit the model to data however, and so give no estimates on the relative subpopulation sizes or turnover rate, as we do in our modelling. Zhang et al. model mouse oocyte development using a two-population model whereby only a fixed-size subpopulation of mtDNA is able to replicate, yielding linear growth in oocyte copy-number, and explore selective mechanisms to explain observed mutation propagation[8]. This model is stochastic, and is quantitatively fit to non-proliferative cells. However, the model is by design fit to out-of-equilibrium oocyte cells, and so replication rate estimates are not applicable to post-mitotic human cells, for example. Degradation is also not embedded into their model, so preferential degradation is not explored. Finally, Davis and Clayton observe that, in PC12 cells, replicating mtDNA foci tend to be found closer to the nucleus, suggesting subpopulations of mtDNA that are spatially segregated[9].

In short, precedent for multiple subpopulations of mtDNA already exist in the literature. However, none embed both preferential replication of young mtDNA and preferential degradation of old mtDNA, as our model does, and few give quantitative estimates of such subpopulations and associated replication rates. While our work points to subpopulations, our timescales indicate they are not long lived (see main text Fig. 6p). Further, other than Zhang et al.[8], which works with out-of-equilibrium oocytes, all papers mentioned here work with proliferative cells, which makes isolating the effect of mtDNA replication technically challenging from a modelling perspective. Our work therefore not only extends on these models in a biologically meaningful way, but is fit to non-proliferative cells, ensuring there is no confounding from the cell cycle.

#### 1.6 Nucleoids and MtDNA

We use nucleoid count as a proxy for mtDNA count throughout. While nucleoids typically contain one mtDNA, the average number of mtDNA per nucleoid is $\sim 1.4$[10]. Here we discuss how this may affect our inference.

A simple way of incorporating this into our modelling is to multiply all observed cellular nucleoid and EdU tagged nucleoid numbers by 1.4 prior to performing our stochastic modelling fits. Since this leaves the ratio of mtDNA to EdU tagged mtDNA unchanged, this should not effect our inference of the per-capita replication rate, nor should it change the inferred values of any other Poisson rate parameters ($\mu_d, \mu_a$, etc.), since these are all per-capita rates. The only parameters which will change are the linear regression parameters $\beta_0, \beta_1$, and potentially the control strength $c$. $\beta_0, \beta_1$ in particular would necessarily become larger, however, the relationship between mitochondrial volume and mtDNA copy number will remain linear. Critically, since the inference of the Poisson rate parameters remains unchanged, our inference of the vast majority of quantities of interest should also remain unchanged. This includes, for instance, the quoted replication rate in the main text ($1.5 \pm 0.2\%$), subpopulation size estimates, dwell time estimates, and probability of repeated replication $q$. This justifies our use of nucleoid count as a proxy for mtDNA count.

It should be noted, however, that although here we have argued that the distinction between nucleoids and mtDNA does not affect our parameter inference in our stochastic models, the distinction becomes critical when probing for causality - at least in some extreme scenarios (Section 7). Seel et al. observe that in budding yeast cells, inducing a mutation which reduces mitochondrial volume but holds cellular volume the same results in a reduction of nucleoid number, but no change in mtDNA copy number[11]. This suggests that nucleoid number is linked to mitochondrial volume, whereas mtDNA number is, at least in some extreme scenarios, linked to cellular volume. Indeed, this agrees with our causal analysis, which looks only at nucleoid number, and concludes that it is linked to mitochondrial volume (Section 7). The results of this section therefore must be taken to be for nucleoids in a physiological context, rather than mtDNA in the context of nuclear mutation.

### 2 Stochastic Systems

Section 1 introduced us to four main models: the one-population model, the preliminary two-population model, the two-population model, and the three-population model. In this section, we will give full mathematical details of these models necessary for simulation, incorporating EdU as in the pulse phase of pulse and pulse-chase data, and removing EdU as in the chase phase of the pulse-chase data. Since the preliminary two-population model is a special case of the two-population model where $p = 1$, it will not be referred to for the remainder of this section.

#### 2.1 One-Population Model

##### 2.1.1 Equilibrium Dynamics

Let $(N, l, V)$ be the nucleoid number, mitochondrial volume and cell volume of a cell respectively, and let $n$ refer to a single nucleoid. As discussed in Section 1, the one-population model assumes that each nucleoid in the system is subject to two Poisson processes - a birth process and a death process, with rates $\lambda_b, \mu_d$ respectively. Whenever a nucleoid undergoes a replication event, it doubles, and whenever a nucleoid undergoes a death event, it is removed from the system. These equilibrium dynamics are encapsulated within the following events, and in Supplementary Fig. 2a:

$$\begin{aligned} n &\xrightarrow{\lambda_b} 2n \\ n &\xrightarrow{\mu_d} \phi, \end{aligned} \tag{5}$$

where $\phi$ indicates degradation, and $n$ represents a single nucleoid. As discussed in Section 1, the death rate $\mu_d$ is constant, and the birth rate $\lambda_b \equiv \lambda_b(N, l)$ is a function that varies to maintain copy number control, which in general, has the form $\lambda_b(N, l) = \mu_b + f(N, l)$, where $f$ is some feedback law satisfying Equation 2. $N_{opt}$ of Equation 2 must satisfy $N_{opt}(l) = \beta_0 + \beta_1 l$ for constants $\beta_0, \beta_1$ (Equation 3), since we observe a linear relationship between $N$ and $l$ in the data. This definition of $\lambda_b$, $\mu_b$, $f$, and $N_{opt}$ will remain the same for the two- and three-population models (Sections 2.2 and 2.3). As mentioned in Section 1.1, the birth rate is not a function of cell volume $V$, motivated by evidence concerning the causal relationship between $(N, l, V)$ explored in Section 7. Richer models could define $l$ itself as a stochastic process - we consider an example richer model in Section 6.4

By definition, we must have $f(N_{opt}, l) = 0$, so $\lambda_b(N_{opt}, l) = \mu_b$. At this optimal copy number we should have cellular equilibrium, which for the one-population model necessitates the condition $\mu_b = \mu_d$. This condition will be different for the two- and three-population models. Again, as mentioned in Section 1.1, and covered in Section 6, we choose logarithmic birth (Equation 4c) to be our birth rate function $\lambda_b$, due to its heteroscedastic properties.

###### 2.1.2 EdU Incorporation (Pulse Phase)

Let $N^{(0)}, N^{(1)}, N^{(2)}$ be the number of untagged nucleoids, singly tagged, and doubly tagged respectively, so that $N = N^{(0)} + N^{(1)} + N^{(2)}$, and let $n^{(0)}, n^{(1)}, n^{(2)}$ refer to one member of the untagged, singly tagged, and doubly tagged populations, respectively. Then our events with EdU incorporation factored in become:

$$\begin{aligned} &n^{(0)} \xrightarrow{\lambda_b} 2n^{(1)} \\ &n^{(0)} \xrightarrow{\mu_d} \phi \\ &n^{(1)} \xrightarrow{\lambda_b} n^{(1)} + n^{(2)} \\ &n^{(1)} \xrightarrow{\mu_d} \phi \\ &n^{(2)} \xrightarrow{\lambda_b} 2n^{(2)} \\ &n^{(2)} \xrightarrow{\mu_d} \phi. \end{aligned} \tag{6}$$

See also Supplementary Fig. 3a. That is, whenever an untagged molecule replicates, it becomes two singly tagged molecules; a singly tagged molecule becomes one singly tagged and one doubly tagged molecule, and a doubly tagged molecule becomes two doubly tagged molecules. This follows from the genetics of the system: the two strands of the mtDNA split, and each gains a tagged strand to accompany it.

###### 2.1.3 EdU Removal (Chase Phase)

During the chase portion of the pulse-chase data, i.e., at the 24 hour time point of the pulse-chase data, EdU incorporation flips. That is, whenever a double tagged molecule replicates, it becomes two single tagged molecules; whenever a single tagged molecule replicates, it becomes one single tagged and one untagged molecule; and whenever an untagged molecule replicates, it becomes two untagged molecules. This gives the exact same diagrams as in Supplementary Fig. 3a, but with green representing an EdU tagged strand, and magenta representing an untagged strand, and is represented by the following stochastic systems equations:

$$\begin{aligned} &n^{(2)} \xrightarrow{\lambda_b} 2n^{(1)} \\ &n^{(2)} \xrightarrow{\mu_d} \phi \\ &n^{(1)} \xrightarrow{\lambda_b} n^{(1)} + n^{(0)} \\ &n^{(1)} \xrightarrow{\mu_d} \phi \\ &n^{(0)} \xrightarrow{\lambda_b} 2n^{(0)} \\ &n^{(0)} \xrightarrow{\mu_d} \phi. \end{aligned} \tag{7}$$

#### 2.2 Two-Population Model

###### 2.2.1 Equilibrium Dynamics

The two-population model assumes that there is a subpopulation of replicating nucleoids $\mathcal{N}_r$ containing $N_r$ nucleoids, and a subpopulation of nonreplicating ('old') nucleoids $\mathcal{N}_o$ containing $N_o$ nucleoids, so that $N = N_r + N_o$. Let $n_r, n_o$ denote one member of the replicating and nonreplicating population respectively. The birth rate $\lambda_b$ now modulates the rate at which nucleoids begin replicating, i.e., move from population $\mathcal{N}_o$ to $\mathcal{N}_r$, and we have a new constant replication-termination rate $\mu_r$ which modulates how quickly a nucleoid finishes replicating. Whenever a nucleoid finishes replicating, there is some probability $p$ that the replicating machinery diffuses away, and the two daughter molecules move to the nonreplicating population $\mathcal{N}_n$, and a probability $q = 1 - p$ that the replicating machinery does not diffuse away, but begins to replicate one of the daughter molecules. Finally, nucleoids in the nonreplicating population can degrade with constant rate $\mu_d$. This gives rise to the following stochastic systems equations:

$$\begin{aligned} n_o &\xrightarrow{\lambda_b} n_r \\ n_r &\xrightarrow{p\mu_r} 2n_o \\ n_r &\xrightarrow{(1-p)\mu_r} n_r + n_o \\ n_o &\xrightarrow{\mu_d} \phi \end{aligned} \tag{8}$$

See also Supplementary Fig. 2c. By definition, $\lambda_b(N_{opt}, l) = \mu_b$ (Section 2.1.1). Unlike in the one-population model, it is not the case that $\mu_b = \mu_d$ in order to maintain copy number control. To find the new requirement on $\mu_b$ at equilibrium, we suppose that we are in the steady state, with $N = N_{opt}$, and therefore $\lambda_b = \mu_b$. We further assume that in the steady state, the number of nucleoids in each of the subpopulations $N_o$ and $N_r$ are constant over time, and so the rate at which molecules leave each population equals the rate at which molecules enter. For population $\mathcal{N}_o$, writing the net flow out of the population on the left hand side, and the net flow into on the right, this gives us:

$$\begin{aligned} \mu_b N_o + \mu_d N_o &= (1-p)\mu_r N_r + 2p\mu_r N_r \\ \implies (\mu_b + \mu_d) N_o &= (1+p)\mu_r N_r \end{aligned} \tag{9}$$

Similarly for population $\mathcal{N}_r$:

$$p\mu_r N_r = \mu_b N_o \tag{10}$$

These simultaneous equations can be readily solved by dividing:

$$\frac{\mu_b + \mu_d}{\mu_b} = \frac{(1+p)}{p}$$

which leaves us with:

$$\mu_b = p\mu_d \tag{11}$$

We therefore see that $\mu_b$ must be a constant $p$ multiple of the death rate $\mu_d$ in order for the system to have a stable copy number. From the above simultaneous equations also follows the ratio of molecules in each subpopulation:

$$\begin{aligned} &N_r : N_o \\ &= N_r : \frac{p\mu_r}{\mu_b} N_r \\ &= \mu_d : \mu_r, \end{aligned} \tag{12}$$

where in the first equality we have substituted the steady state equation for $\mathcal{N}_r$, and in the second we have used the fact that $\mu_b/p = \mu_d$. From here the fraction of molecules in each subpopulation easily follows:

$$\begin{aligned} f_r &:= \frac{N_r}{N} = \frac{\mu_d}{\mu_d + \mu_r} \\ f_o &:= \frac{N_o}{N} = \frac{\mu_r}{\mu_d + \mu_r}. \end{aligned} \tag{13}$$

These proportions will be important in order to initialise these models when it comes to parameter inference (Section 3).

##### 2.2.2 EdU Incorporation (Pulse Phase)

Like in the one-population model, let $N_o^{(0)}, N_o^{(1)}, N_o^{(2)}$ denote the number of nonreplicating untagged, single tagged and double tagged molecules, and let $N_r^{(0)}, N_r^{(1)}, N_r^{(2)}$ denote the number of replicating untagged, single tagged and double tagged molecules, so that $N_\alpha = \sum_{i=0}^{3} N_\alpha^{(i)}$, for $\alpha = r$ or $o$. Let $n_\alpha^{(i)}, \mathcal{N}_\alpha^{(i)}$ denote a single member of the population, and the whole population, respectively, for $\alpha = r$ or $o$; $i \in \{0, 1, 2\}$. Like in the one population case, whenever a replicating molecule (i.e., in $\mathcal{N}_r$) finishes replicating, its two strands split, and each gains a tagged strand to accompany it. The added complexity with this two-population model is that there is a probability $q = 1 - p$ that one of these two daughter molecules is immediately chosen to replicate again (and either daughter is equally likely to be chosen). This leads to the more complex stochastic diagram shown in Supplementary Fig. 3b, with the full systems given in Equation 64 in Appendix D.

##### 2.2.3 EdU Removal (Chase Phase)

EdU removal for the two-population model behaves analogously to the one-population model, where EdU incorporation flips (see Section 2.1.3). Again, this leads to the same stochastic system as in Supplementary Fig. 3b, but with green representing an EdU tagged strand, and magenta representing an untagged strand. With the two-population model however, there is another added complexity: the replicating population $\mathcal{N}_r$ from the pulse portion will have already incorporated EdU before the beginning of the chase phase, where EdU is removed. These pulse 'replicating remnants' will therefore behave differently than molecules which begin to replicate during the chase phase.

It is not obvious precisely how a population which begins replicating with EdU in the system but finishes with no EdU in the system will behave. To model this population, we assume that nucleoids in this population spent the first half of their replication cycle within the pulse portion of the experiment exposed to EdU, and the second half of their replication cycle within the chase phase of the experiment, not exposed to EdU. According to the most up-to-date models of mtDNA replication, both strands of the mtDNA are not replicated at the same time, but rather one strand begins accumulating bases first, and once 2/3 of this strand has been replicated, only then does the other strand begin to replicate[12]. In our model therefore, our assumption that half of the time spent replicating is in the pulse portion, and half is in the chase phase can be interpreted as assuming that one of the strands of the mtDNA is replicated during the pulse phase, whereas one is replicated during the chase phase. Hence, one strand will gather an EdU tagged strand to accompany it, while the other will gather an untagged strand to accompany it.

Mathematically, we account for this behavior by introducing the 'remnant replicating' population $\mathcal{N}_{rem}$, which consists of nucleoids which were in the population $\mathcal{N}_r$ (which includes both nucleoids that are being newly replicated and those which have been recycled with probability $1 - p$) at the end of the pulse portion of the experiment. When a replication event occurs, a nucleoid in $\mathcal{N}_{rem}$ will split into 2 daughter nucleoids which get tagged according to the dynamics just discussed. For instance, if an untagged remnant nucleoid replicates, one possible event is $n_{rem}^{(0)} \xrightarrow{p\mu_r} n_o^{(1)} + n_o^{(0)}$, encapsulating the assumption that one of its daughter particles gathers a tagged strand, whereas the other does not. Here, as usual $n_{rem}$ denotes one nucleoid in $\mathcal{N}_{rem}$. If one of the daughter molecules is chosen to keep replicating, they move to the replicating population $\mathcal{N}_r$ (the ordinary replicating population, not the remnant replicating population $\mathcal{N}_{rem}$), as they have now began replicating during the chase portion of the experiment. For this untagged example, this consists of the two events: $n_{rem}^{(0)} \xrightarrow{(1-p)\mu_r/2} n_r^{(1)} + n_o^{(0)}$, $n_{rem}^{(0)} \xrightarrow{(1-p)\mu_r/2} n_o^{(1)} + n_r^{(0)}$, which occur with equal probability as each daughter nucleoid is equally likely to be chosen to replicate again. Full equations for this model are given in Appendix D Equation 65.

It should be noted that hypothetically, we could have chosen to model the introduction of EdU at the 0 hour time point of the pulse phase similarly; that is, add a remnant replicating population of molecules which begin replicating in an environment with no EdU and finishes in an environment with EdU, modelling these molecules as producing one singly tagged and one untagged molecule. We instead chose to keep the model simpler by assuming these replicating molecules produce two singly tagged molecules upon completion, as is typical in previous models[5]. It is not clear which of these modelling assumptions is more biologically accurate. We note, however, that since this population will decay rapidly, and we do not use the proportion of singly tagged molecules at the 0 hour time point during inference (Section 3.2), the effect of this modelling choice on inference should be minimal.

#### 2.3 Three-Population Model

##### 2.3.1 Equilibrium Dynamics

The three-population model adds one more population to the two-populations $\mathcal{N}_r$, $\mathcal{N}_o$ of the two-population model: the 'young' population $\mathcal{N}_y$, which is shielded from degradation (see Section 8). The model now has four classes of events that can occur: birth events, replication-termination events, ageing events, and death events. From left to right, a birth event with rate $\lambda_b$ causes a young nucleoid in $\mathcal{N}_y$ to begin replicating, and hence move into the population $\mathcal{N}_r$. A replication-termination event with rate $\mu_r$ causes a nucleoid in $\mathcal{N}_r$ to finish replicating producing two nucleoids. With probability $p$ both daughter nucleoids will move to $\mathcal{N}_y$, and with probability $q = 1-p$, one will remain in $\mathcal{N}_r$ to keep replicating. An ageing event with rate $\mu_a$ will cause a nucleoid in the young population $\mathcal{N}_y$ to move to the old population $\mathcal{N}_o$, where it is permitted to degrade with rate $\mu_d$. Critically, this embeds into the model the idea that when a nucleoid has just finished replicating, there is a purgatory period where it will remain in the young population, where it is not permitted to degrade, before either replicating again or ageing. This captures the intuition of Section 1.4 that newly replicated molecules cannot degrade. We obtain the following stochastic system:

$$\begin{aligned} &n_y \xrightarrow{\lambda_b} n_r \\ &n_r \xrightarrow{p\mu_r} 2n_y \\ &n_r \xrightarrow{(1-p)\mu_r} n_r + n_y \\ &n_y \xrightarrow{\mu_a} n_o \\ &n_o \xrightarrow{\mu_d} \phi. \end{aligned} \tag{14}$$

This adds one more parameter $\mu_a$ to the two-population model, increasing the complexity by one degree of freedom. Like in the two-population model, we can compute the condition on the parameters necessary for the existence of a steady state. Notice that Equations 14 and 8 are identical in structure for the first four equations up to relabelling of variables $n_y \sim n_o$, $n_o \sim \phi$, $\mu_a \sim \mu_d$ (where the left hand side is three-population label, and the right is the two-population label). It immediately follows that enforcing steady state on populations $\mathcal{N}_r$, $\mathcal{N}_y$ will lead to identical equations to Equations 9 and 10, up to relabelling, i.e.:

$$\begin{aligned} &(\mu_b + \mu_a)N_y = (1+p)\mu_r N_r \\ &p\mu_r N_r = \mu_b N_y \end{aligned} \tag{15}$$

We have already solved this equation in Section 2.2.1, and we obtain:

$$\mu_b = p\mu_a \tag{16}$$

Interestingly, the birth rate $\mu_b$ and the death rate $\mu_d$ are no longer directly coupled, but rather it is the birth rate and the ageing rate $\mu_a$ which are coupled for copy number control. Enforcing steady state on the remaining population $\mathcal{N}_o$ gives us:

$$\mu_a N_y = \mu_d N_o. \tag{17}$$

From here, like in Section 2.2, we may compute the ratio of molecules in each subpopulation at equilibrium:

$$\begin{aligned}
&N_r : N_y : N_o \\
=&N_r : \frac{p\mu_r}{\mu_b} N_r : \frac{\mu_a}{\mu_d} N_o \\
=&N_r : \frac{p\mu_r}{\mu_b} N_r : \frac{\mu_a}{\mu_d} \frac{p\mu_r}{\mu_b} N_r \\
=&1 : \frac{\mu_r}{\mu_a} : \frac{\mu_r}{\mu_d} \\
=&\frac{1}{\mu_r} : \frac{1}{\mu_a} : \frac{1}{\mu_d},
\end{aligned} \tag{18}$$

where in the above computation we liberally use the fact that $\mu_b = p\mu_a$, as well as the steady state Equations 15 and 17. Despite the seeming complexity of the model, we obtain a remarkably simple equation for the population ratios, which can be directly compared to the analogous Equation 12 in the two-population model (rewritten slightly): $N_r : N_o = \frac{1}{\mu_r} : \frac{1}{\mu_d}$. This is not surprising, as one can think of the two-population model as the limit of the three-population model as $\mu_d \to \infty$, up to some relabelling. From Equation 18, one can easily write the fraction $f_\alpha$ of nucleoids in each subpopulation $\mathcal{N}_\alpha$ at steady state, for $\alpha = r, y, o$, which will be used to initialise the model when simulating.

For the three-population model, we also detail how we can compute dwell times in each subpopulation. Let $T_r$, $T_d$, $T_a$ be the mean time it takes for a nucleoid to replicate once given that it is in $\mathcal{N}_r$ (note: this is not the time it takes for the nucleoid to leave $\mathcal{N}_r$ fully, as repeated more subsequent replication events may occur), to die given that it is in $\mathcal{N}_o$, and to age given that it is in $\mathcal{N}_r \cup \mathcal{N}_y$, respectively. The dwell times in the replicating and the old populations are exponentially distributed with means $1/\mu_r$, $1/\mu_d$ respectively, and so it follows easily that:

$$\begin{aligned}
T_r &= \frac{1}{\mu_r} \\
T_o &= \frac{1}{\mu_o}.
\end{aligned} \tag{19}$$

Computing $T_a$ is more difficult, since molecules can move between $\mathcal{N}_y$ and $\mathcal{N}_r$ many times before ageing into $\mathcal{N}_a$. To this end, let $Q$ be the generator matrix, so that $Q_{ij}$ is the Poisson rate of a single nucleoid transitioning from state $i$ to $j$, if $i \neq j$, and $Q_{ii} = -\sum_{j\neq i} Q_{ij}$ is the negative of the total outward flux from state $i$. To be clear, here, we are indexing based on the different subpopulations, with $i, j \in \{0, 1, 2\}$, and $0, 1, 2$ referring to $\mathcal{N}_r$, $\mathcal{N}_y$, $\mathcal{N}_o$, respectively. Under this framework, events where the nucleoid duplicates must be treated with care. If a nucleoid duplicates, we choose at random one of the daughter nucleoids to continue tracking as the 'original' nucleoid. With this treatment, the rate of movement from $\mathcal{N}_r$ to $\mathcal{N}_y$ is given by:

$$Q_{01} = p\mu_r + (1-p)\mu_r/2 = \frac{\mu_r}{2}(1+p)$$

The first term, $p\mu_r$, arises from the rate at which molecules replicate and both daughters move to $\mathcal{N}_y$. The second term $(1-p)\mu_r/2$ arises from the rate at which molecules replicate and only one daughter moves to $\mathcal{N}_y$, multiplied by the probability (1/2) that we select the daughter which moves to keep tracking as the 'original' nucleoid. The remainder of the matrix $Q$ is easily populated:

$$Q = \begin{pmatrix} -\frac{\mu_r}{2}(1+p) & \frac{\mu_r}{2}(1+p) & 0 \\ \mu_b & -(\mu_b + \mu_d) & \mu_d \\ 0 & 0 & 0 \end{pmatrix}, \tag{20}$$

where here we treat $\mathcal{N}_o$ as an absorbing state, and since we are in equilibrium, $\lambda_b = \mu_b$. Now, let $T_a^i$ be the expected time until ageing for a molecule in state $i$. Then, since the expected dwell time in state $i$ is $\frac{1}{-Q_{ii}}$, and the nucleoid has probability $\frac{Q_{ij}}{-Q_{ii}}$ of moving to state $j$ when an event occurs, we can write the following recursion relation, for $i = 0, 1$:

$$
\begin{aligned}
T_a^i &= \frac{1}{-Q_{ii}} + \sum_{j \neq i} \frac{Q_{ij}}{-Q_{ii}} T_a^j \\
\implies -1 &= \sum_j Q_{ij} T_a^j \\
\implies \tilde{Q} \boldsymbol{T}_a &= -I,
\end{aligned}
\tag{21}
$$

where $\boldsymbol{T}_a = [T_a^0, T_a^1]^T$, $\tilde{Q}$ is the $2 \times 2$ submatrix of $Q$ of transient states $0, 1$, and $I$ is the $2 \times 2$ identity matrix. This matrix equation can be simply solved by inverting:

$$
\boldsymbol{T}_a = -\tilde{Q}^{-1} = \frac{2}{\mu_r \mu_a} \begin{pmatrix} \mu_b + \mu_a + \frac{1+p}{2}\mu_r \\ \mu_b + \frac{1+p}{2}\mu_r \end{pmatrix}. \tag{22}
$$

Finally, given that a nucleoid is in $\mathcal{N}_r \cup \mathcal{N}_y$, at equilibrium, it has probability $p_r = \frac{1/\mu_r}{1/\mu_r + 1/\mu_a}, p_y = \frac{1/\mu_a}{1/\mu_r + 1/\mu_a}$ of being in $\mathcal{N}_r, \mathcal{N}_y$, from the subpopulation proportions we have just computed (Equation 18). Letting $\boldsymbol{p} = [p_r, p_y]^T$, we can finally write the expected ageing time from the quantities we have already computed:

$$
T_a = \boldsymbol{T}_a \cdot \boldsymbol{p}. \tag{23}
$$

###### 2.3.2 EdU Incorporation (Pulse Phase)

EdU incorporation works in the same way for the three-population model as in the two-population model (see Section 2.2.2). We adopt analogous notation where $\mathcal{N}_\alpha^{(i)}$ denotes the subpopulation of $\mathcal{N}_\alpha$ which has $i$ strands tagged; $n_\alpha^{(i)}$ denotes a member of this population; and $N_\alpha^{(i)}$ denotes the total number of elements of this population. Stochastic systems equations are given in Equation 66 in Appendix D, and is visualised in Supplementary Fig. 3c.

###### 2.3.3 EdU Removal (Chase Phase)

Again, the chase portion of the experiment works in the same way as in the two-population model, with EdU incorporation flipping and the new replicating remnant population $\mathcal{N}_{rem}$ being introduced (see Section 2.2.3). Stochastic systems equations are given in Equation 67 in Appendix D.

#### 2.4 Measured EdU Intensities of Replicating Molecules

In the pulse data and the pulse-chase data, we have a certain number of molecules that are EdU tagged. In the intensity data, the EdU tagged molecules are further broken down into singly tagged and doubly tagged molecules (see Section 1.3 and Section 3.1.3). We do not attempt to generate the full intensity distributions we observe in the biological data from our models, rather, we assign to each subpopulation a 'measured EdU-level': the number of EdU tags the population would be measured as having, and assume that the observed intensity distributions arise from noise both in the amount of EdU accumulated and from measurement. For every subpopulation but the replicating and remnant replicating subpopulations (the subpopulation of molecules which began replicating in the pulse and finished in the chase, see Section 2.2.3) $\mathcal{N}_r$, $\mathcal{N}_{rem}$, the question of how many tags the nucleoid has is trivial: $\mathcal{N}_\alpha^{(i)}$ is measured as having $i$ tagged strands, for $\alpha \neq r, rem$. For $\mathcal{N}_r^{(i)}$

or $\mathcal{N}_{rem}^{(i)}$ however, the question becomes more complex, as these populations are in the process of incorporating EdU (depending on whether we're in the pulse or chase phase of the experiment). We first focus on what this looks like during the pulse portion of the experiment.

##### 2.4.1 Pulse Portion

During the pulse portion of the experiment a nucleoid in population $\mathcal{N}_r$ is one which is currently in the process of incorporating EdU (and the population $\mathcal{N}_{rem}$ does not exist). It thus is not clear what EdU intensity (single, or double, or more) would actually be measured. As discussed in Section 1.2, we assume that the replicating population $\mathcal{N}_r^{(0)}$ gets tagged immediately when EdU is introduced into the system to account for the large EdU number in the pulse data at the 1 hour time point, and so all three populations $\mathcal{N}_r^{(i)}$ are measured as EdU tagged, for $i = 0, 1, 2$.

What EdU intensity would these replicating populations be measured as having? Focusing on $\mathcal{N}_r^{(0)}$, biologically, an mtDNA in this population is one which was untagged, and is currently splitting its two strands, gathering up two new EdU tagged strands to accompany each of its previous strands. At the beginning of its replication therefore, this nucleoid would be measured as having 0 tagged strands, but at the end, it would be measured as having 2 tagged strands. For mathematical simplicity, since we are assuming this population gets tagged immediately, we will be assuming that this population is measured as having 2 tagged strands. With the same logic, $\mathcal{N}_r^{(1)}$ and $\mathcal{N}_r^{(2)}$ would be measured as having 3 and 4 tagged strands, respectively, which we put under the blanket category of '> 2 tagged strands'.

Even if this approximation is not precise, it ultimately does not affect our inference significantly, for two reasons. The first is that, as we will see in Section 4, the proportion of molecules in this replicating population $\mathcal{N}_r$ is small: roughly 10%, so the contributions of these molecules to the overall singly tagged population (which is the population we use in our inference) is minimal. Secondly, our Gaussian mixture modelling estimate of the singly tagged proportion in the data (Section 3.1.3) is also only meant to be a rough estimate of the number of nucleoids that are single tagged. In other words, we are not relying on these estimates being precise: only that they capture the broad behaviour of the system. This rough behaviour is enough to distinguish between the one-population model and the two-/three-population models, as we will see in Section 4.

To conclude this subsection, for all three of the one-, two-, and three-population models, we define $E^{(1)}, E^{(2)}, E^{(>2)}$ to be the number of molecules measured as single tagged, double tagged, and more than double tagged, respectively, and $E = E^{(1)} + E^{(2)} + E^{(>2)}$ to be the number of tagged molecules in general. As per the discussion of this section, for the two- and three-population models, we have:

$$\begin{aligned} E^{(1)} &= \sum_{\alpha \neq r} N_\alpha^{(1)} \\ E^{(2)} &= \sum_{\alpha \neq r} N_\alpha^{(2)} + N_r^{(0)} \\ E^{(>2)} &= N_r^{(1)} + N_r^{(2)}. \end{aligned} \tag{24}$$

For the one-population model, the population $\mathcal{N}_r$ does not exist, so the above equations become trivial: $E^{(1)} = N^{(1)}$, $E^{(2)} = N^{(2)}$, $E^{(>2)} = 0$.

##### 2.4.2 Chase Portion

The logic of the chase portion is similar, except now the population $\mathcal{N}_r$ is not accumulating EdU. This is therefore simpler than the pulse portion: $\mathcal{N}_r^{(i)}$ should be measured as having $i$ tagged strands. The remnant replicating population $\mathcal{N}_{rem}$ however is an artifact left from the pulse section which we are assuming incorporates EdU into one of its strands and does not incorporate EdU in the other (see Section 2.2.3). Thus, we assume that $\mathcal{N}_{rem}^{(i)}$ is measured as having $i+1$ tagged strands, as it is only incorporating one more tagged strand. For the chase portion of the experiment, we therefore have, for the two- and three-population models:

$$\begin{aligned} E^{(1)} &= \sum_{\alpha \neq rem} N_{\alpha}^{(1)} + N_{rem}^{(0)} \\ E^{(2)} &= \sum_{\alpha \neq rem} N_{\alpha}^{(2)} + N_{rem}^{(1)} \\ E^{(>2)} &= N_{rem}^{(2)}. \end{aligned} \tag{25}$$

The one-population model is, again, more simple: $E^{(1)} = N^{(1)}$, $E^{(2)} = N^{(2)}$, $E^{(>2)} = 0$.

### 3 Approximate Bayesian Computation

All parameters of our stochastic systems models were fit using approximate Bayesian computation (ABC)[13]. This technique is a simulation based inference which aims to emulate a traditional Bayesian analysis in the case where the likelihood function is unknown. Briefly, we choose a prior $\pi(\boldsymbol{\mu})$ on our parameters $\boldsymbol{\mu}$, and draw some number $n$ prior parameter samples $\{\boldsymbol{\mu}_i\}_{i=1}^{n} \overset{iid}{\sim} \pi$. For each parameter sample $\boldsymbol{\mu_i}$, we simulate our model, which can be treated as a likelihood function, obtaining some output $\hat{Y}_i \sim l(\cdot|\boldsymbol{\mu_i})$. If $Y$ is our true data, then we accept a parameter sample $\boldsymbol{\mu}_i$ if $d(S(\hat{Y}_i), S(Y)) < \epsilon$, for some small $\epsilon > 0$, distance function $d$, and summary statistics $S$. This leaves us with some collection $\{\boldsymbol{\mu}_{k_i}\}_{i=1}^{m}$ of 'good parametrisations', and, in fact, this collection is a sample from an approximate posterior distribution. For a more detailed discussion of ABC, see Sisson et al[14].

We fitted our models separately to the pulse cells and to the pulse-chase cells to account for slightly different experimental conditions. In the pulse portion of the pulse and pulse-chase cell experiments, copy number control is maintained, and thus we assumed that the system is in equilibrium. Under this assumption, recall that Equations 11 and 16 hold ($\mu_a = \mu_b/p$ (three-population), $\mu_b = p\mu_d$ (two-population), $\mu_b = \mu_d$ (one-population)). Therefore, a degree of freedom is dropped for each model, and so there is one less parameter we need to infer. For the chase portion of the pulse-chase data however, copy number control is not maintained, meaning that the parametrisation of our models must have changed in-between the two phases of the experiment. Section 3.5.2 details how we account for this. When fitting our models, we only used data from assays 1 and 2 of the pulse cells and pulse-chase cells, leaving the third assay for validation (Section 4.6).

We fitted all three models (one-, two-, and three-population) to the pulse cells. In Section 4 we will see that only the two- and three-population models fit this data adequately, and so we only fit the two- and three-population models to the pulse-chase data, where we will subsequently see that only the three-population model fits adequately. The remainder of this section details precisely how this ABC inference was performed. In this section, it becomes necessary to distinguish between measured and simulated data more carefully. To that end, we will denote a simulated quantity with a hat: $\hat{X}$, and a measured quantity without: $X$.

#### 3.1 Pre-ABC Data Analysis

##### 3.1.1 Measurement Error

When doing our inference, we have to account for any possible sources of systematic error within the pulse and pulse-chase data. We identified two. First, if the brightness of a nucleoid is lower than a set brightness threshold, the software does not detect and count the nucleoid. This leads to a systematic undercounting of nucleoids in each cell. To account for this, we manually counted the number of tagged and untagged nucleoids in 28 cells, and compared this figure to the number counted automatically by the software. We observed that the percentage of tagged and untagged molecules missed fit an exponential distribution, with means 3.41%, 3.83%, for the tagged and untagged populations respectively, as is shown by the Q-Q plots of Supplementary Fig. 4. As these errors are so close, we model this as a single error of 3.6% for all nucleoids. We take this into account in our models by drawing a random error $e \sim Exp(0.036)$ for each simulated cell and multiplying the final nucleoid number output (of any subpopulation) by $1 - e$. Although we account for these errors, practically speaking, they are small enough to not have any significant effect on our analysis.

Second, another source of 'undercounting' noise comes from colocalisation of nucleoids. If two nucleoids are sufficiently close, they are indistinguishable from the perspective of the measuring apparatus. However, since our microscope has such high resolution (between $50nm$ and $100nm$ in diameter), we found by a simple calculation that this form of measurement error should be negligible (Appendix C).

###### 3.1.2 $\beta_0, \beta_1$ Inference

Recall the parameters $\beta_0, \beta_1$ from Equation 3. To fit $\beta_0$ and $\beta_1$, we need not do a full ABC, but rather we can simply perform a regression. This allows the ABC to perform more efficiently as it has less parameters it is required to fit. To adjust this inference for the fact $N$ is slightly undercounted, we divide the nucleoid number in each cell by $1 - 0.036 = 0.964$, i.e., the mean error from the previous section, and we will refer to this as the adjusted nucleoid number $N_{adj} = N/0.964$. As previously mentioned, we do this inference separately for the pulse cells and the pulse-chase cells to account for any experimental differences. For the pulse cells, we use every cell from all 4 time points of assays 1 and 2. For the pulse-chase cells, we also use assays 1 and 2, however, we can only use the first time point $t = 24$, as the system is not in equilibrium for subsequent time points.

We use a Bayesian linear regression in order to be more consistent with the ABC. As a preliminary analysis, we performed an OLS regression to the pulse cells, and did a subsequent OLS regression without intercept on the absolute residuals (Supplementary Fig. 5a). We found that the regression without intercept fit the absolute residuals well, indicating linear heteroscedasticity in the data. Due to this heteroscedasticity, we use the methodology of Startz which is robust to this[15]. Let $\boldsymbol{\beta} = (\beta_0, \beta_1)$, $X = [\mathbb{1}, \boldsymbol{l}]$ be the data matrix, where $\boldsymbol{l} = \{l_i\}_i$ is all of the measured mitochondrial volumes and $\mathbb{1}$ is the vector of ones. Let $Y = \{N_{adj,i}\}_i$ be the response vector. We model our data as:

$$\begin{aligned} N_{adj} &= \beta_0 + \beta_1 l + \epsilon \\ \epsilon &\sim N(0, (l\sigma)^2). \end{aligned} \tag{26}$$

We put a diffuse prior on $\boldsymbol{\beta}$: $\boldsymbol{\beta} \sim N(0, V)$, with $V \to \infty$, and an inverse-Gamma prior on $\sigma^2$: $\sigma^2 \sim I\Gamma(\frac{\nu_0+2}{2}, \frac{\sigma_0^2\nu_0}{2})$, where $\sigma_0, \nu_0$ are hyperparameters we will specify. With this choice of priors, we obtain the conditional posteriors:

$$\begin{aligned} \boldsymbol{\beta}|\sigma^2, \boldsymbol{l} &\sim N(\bar{\boldsymbol{\beta}}, \bar{V}) \\ \sigma^2|\boldsymbol{\beta}, \boldsymbol{l} &\sim I\Gamma(\frac{\alpha_1}{2}, \frac{\delta_1}{2}), \end{aligned} \tag{27}$$

where:

$$\begin{aligned} \bar{\boldsymbol{\beta}} &= (X^TX)^{-1}X^TY \\ \bar{V} &= (X^TX)^{-1}X^T\Sigma X(X^TX) \\ \alpha_1 &= \nu_0 + 2 + n \\ \delta_1 &= \sigma_0^2\nu_0 + \hat{e}^T\hat{e}, \end{aligned} \tag{28}$$

and $\hat{e}_i = \frac{Y_i - X_i\beta}{l_i}$, $\Sigma = \text{diag}(\sigma^2 l_i^2)$ is the diagonal covariance matrix of the residuals, $n$ is the number of datapoints. With these conditional posteriors, we can then use Gibbs sampling to sample from the joint posterior $(\boldsymbol{\beta}, \sigma^2)|\boldsymbol{l}$.

Letting $\pi_{\sigma^2}$ denote the prior of $\sigma^2$, we have $\mathbb{E}[\pi_{\sigma^2}] = \sigma_0^2, \text{Var}[\pi_{\sigma^2}] = \frac{2(\sigma_0^2)^2}{\nu_0 - 2}$. Our OLS regression gave us an absolute parameter estimate of $\sigma = 0.25$, so we set this to be the mean of our prior $\pi_{\sigma^2}$: $\sigma_0 = 0.25$, for both the pulse and the pulse-chase data. We set $\nu_0 = 1$ in order to have a reasonably large prior variance, again for both datasets. Notice that modelling the heteroscedasticity present in the data does not impact the conditional posterior mean of $\boldsymbol{\beta}$: $\bar{\boldsymbol{\beta}} = (X^T X)^{-1} X^T Y$ is the same formula as in a frequentist OLS regression, or a standard Bayesian treatment with diffuse priors. Thus, this more sophisticated analysis only affects the posterior variance $\bar{V}$ in the $\boldsymbol{\beta}$ inference. We obtain posterior means of $\beta_0 = 222$, $\beta_1 = 1.28$, $\sigma^2 = 0.12$ for the pulse cells, and $\beta_0 = 261$, $\beta_1 = 1.37$, $\sigma^2 = 0.10$ for the pulse-chase cells, with full posteriors shown in Supplementary Fig. 5b. Gibbs sampling trace plots are displayed in Supplementary Fig. 5c, showing good mixing, and subsequent posterior predictive distributions are shown in Supplementary Fig. 5d.

Although we now have already inferred $\boldsymbol{\beta}$, we will still pass these posteriors through the ABC methodology in order to propagate the uncertainty. Specifically, we will use the posteriors inferred in this subsection as ABC priors for $\boldsymbol{\beta}$ in subsequent sections to ensure that we are accounting for the uncertainty in $\boldsymbol{\beta}$ when inferring all other parameters in our models.

##### 3.1.3 Singly Tagged Proportions

As discussed in Section 1.3, our main source of evidence for preferential replication of recently replicated molecules (and hence the two- or three-population model over the one-population or preliminary two-population model) is the EdU intensity data coming from the chase portion of the pulse-chase data, and the large implied proportion of double tagged molecules therein. When looking at the distribution of EdU intensities, many are noticeably bimodal, with the second mode occuring at roughly double the intensity of the first (Extended Data Fig. 7). We hypothesise that the first peak corresponds to those molecules that have only one strand tagged, the second peak corresponds to molecules that have both strands tagged, and any higher intensities correspond to replicating nucleoids which may have more than one tagged strand contained within them. To estimate the proportion of tagged molecules that are single tagged at the 24 hour hour time point of the pulse data and pulse-chase data, and at the 120 hour time point of the pulse-chase data, we fit Gaussian mixture models to all 3 assays for these two time points. Although we only used these two time points as our ABC summary statistics, as we will discuss in Section 3.2, we fit every time point for the sake of completeness.

Two other interpretations of this data need to be considered. The first is that the bimodal distribution represents not singly and doubly tagged mtEdU, but rather nucleoid clustering which our microscope was not able to resolve. Under this interpretation, the first peak consists of single nucleoids, while the second consists of two or more nucleoids clustered. This, however, is unlikely. The resolution of our lattice structured illumination microscopy is $110nm$ in diameter, approximately the same diameter of a nucleoid as measured in human fibroblast cells[16], a resolution high enough to be able to resolve individual nucleoids. The second interpretation is that the two peaks could represent differences in nucleoid packaging. Some nucleoids will contain one mtDNA, and some will contain two (or more), with an average of 1.4[10]. The second peak could thus simply be nucleoids which contain two or more mtEdU molecules, while the first peak consist of nucleoids which contain only one. This, however, does not match what is observed in the data. Assuming nucleoids contain either one or two mtDNA, an average of 1.4 implies that approximately 60% of nucleoids contain one mtDNA and 40% contain two. Thus, the fraction of nucleoids containing two mtEdUs must be, at most, 40%, but likely smaller, and thus the second peak of the Gaussians must not consist of more area than 40%. We see in Extended Data Fig. 7 however that this is not the case, and the majority of Gaussians consist of a second peak encapsulating at least half of the density. Moreover, even if these second peaks did, in part, represent nucleoids with more than one mtEdU, rather than a single doubly tagged mtEdU, this would still likely represent mtDNA that have replicated more than once, thus, these represent the same phenomenon as far as the model is concerned.

Computing the BIC (Bayesian Information Criterion[17]) score for mixture models of different peaks, we clearly see that two and three peaks fit the data substantially better than one, after which there is a plateau (Supplementary Fig. 6a). We expect that for the 24 hour time point, the distribution should be best captured by three Gaussians due to the prevalence of the remnant replicating population which contains more than two tagged peaks (see Sections 2.4, 2.2.3, 2.3.3). For any subsequent time point, we should only need two Gaussians, as this remnant replicating population should have finished replicating and no longer exist. We may then interpret the area and mean of each Gaussian to be the proportion and mean intensity of the mtEdU in that population, respectively. If we do this, we obtain results that match our interpretation very well. First, we see agreement of the singly tagged estimate across assays for all time points, along with the singly tagged proportion gradually increasing across time points, indicating doubly tagged molecules replicating into singly tagged ones over the course of the chase (Supplementary Fig. 6b). We further find estimates for the singly and doubly tagged intensity which are stable over time and consistent across assays (Extended Data Fig. 7c), and the intensity of the second peak is double the first (Extended Data Fig. 7d). To fully justify our choice of three peaks rather than two for the first time point, we probed the fit with two Gaussians also, which we expect to give skewed results due to the presence of an uncounted for remnant replicating population. Indeed, when fitting two peaks, the singly tagged proportion estimate is larger than all subsequent time points (Supplementary Fig. 6c). This is biologically infeasible as the proportion of singly tagged molecules should increase over the course of the chase as doubly tagged molecules replicate. Along with this, the estimate is much less consistent across assays than when fitting three peaks, indicating misspecification. Overall, we take this as justification for choosing three peaks rather than two for the first time point. Although these choices were made on the above grounds, it should be noted that the precise proportion of molecules that are singly tagged at 24 hours is only important insofar as it is less than we expect under the one-population model. This can be seen intuitively from noting that most of the weight of the distributions are at 6000 arbitrary units at 24 hours, but at 120 hours, most are at 3000 arbitrary units (main text Fig. 6g), suggesting that most of the mtEdU are singly tagged. See Section 4.1 for a more detailed discussion on this point.

Let $P_i^t$ be the single tagged proportion at time point $t$ for assay $i$, and let $\bar{P}^t = \frac{1}{3}\sum_{i=1}^{3} P_i^t$ be the average across the three assays for this time point. We will use $\bar{P}^{24}$ as a summary statistic for both the pulse and the pulse-chase data, and $\bar{P}^{120}$ for just the pulse-chase data. Specifically we found that $P_1^{24} = 0.33, P_2^{24} = 0.39, P_3^{24} = 0.28$, and $P_1^{120} = 0.67, P_2^{120} = 0.58, P_3^{120} = 0.43$, so that $\bar{P}^{24} = 0.33$, $\bar{P}^{120} = 0.56$. The value $\bar{P}^{24} = 0.33$ in particular is striking in that it is far lower than we would suspect under the one-population and preliminary two-population models, as we will see in Section 4.1.

As a final point, it is worth noting that we are using only assays 1 and 2 of the pulse and pulse-chase data, with the purpose of saving the third for validation, but we are using all three assays of the intensity data. The reason for this is that since we only have three estimates per time point of the single tagged proportion, our data is limited and is thus very valuable. It would be counterproductive for the inference to ignore any one of these assays. On this note, due to the lack of data for this statistic, along with the potential for the Gaussian mixture models to give biased estimates (if the underlying distributions are not Gaussian, or we have chosen the incorrect number of peaks, for instance), we do not expect a high level of precision from the intensity data - its purpose rather is to give us a general sense of how many molecules are singly tagged. Even with this lack of precision, this summary statistic will be enough to differentiate between models, as we will see in Section 4.

#### 3.2 Summary Statistics

Let $C^t_{pulse} = (N^t_{pulse}, E^t_{pulse}, l^t_{pulse})$ be an observed cell at time $t$ from the pulse dataset, and let $C^t_{chase} = (N^t_{chase}, E^t_{chase}, l^t_{chase})$ be an observed cell at time $t$ from the pulse-chase dataset, containing the nucleoid number, EdU tagged number, and mitochondrial volume of the cell, respectively.

##### 3.2.1 Pulse Cells

We use the following summary statistics:

$$S_{pulse} = \{\bar{E}^1_{pulse}, \bar{E}^3_{pulse}, \bar{E}^7_{pulse}, \bar{E}^{24}_{pulse}, \bar{P}^{24}, S_{cs}\},$$

where:

- $\bar{E}^t_{pulse}$ is the mean EdU number over assays 1 and 2 at time $t$ in the pulse data.
- $\bar{P}^{24}$ is the mean singly tagged proportion over all 3 assays as inferred from the intensity data in Section 3.1.3 at 24 hours after the beginning of the experiment.
- $S_{cs} = 0$ is the *control strength statistic*. This statistic is a measure of how different the copy number control strength of a simulation is to the real data, and so the 'measured' version of this statistic is by definition 0, as the real data is identical to itself. This statistic is necessary for accurate inference of the control strength parameter $c$. The simulated version of this statistic, $\hat{S}_{cs}$, will be introduced in more detail in Section 3.3.

We do not include mean nucleoid number as a summary statistic, as our inference of $\beta_0$ and $\beta_1$ (Section 3.1.2) already ensures that copy number control is maintained in our simulations, meaning that this summary statistic adds no additional information to the inference. We found that including higher order moments did not lead to a qualitatively better fit, so we also did not include these.

##### 3.2.2 Pulse-Chase Cells

For the pulse-chase cells, we use the following summary statistics:

$$\begin{aligned} S_{chase} = \{&\bar{N}^{24}_{chase}, \bar{N}^{48}_{chase}, \bar{N}^{72}_{chase}, \bar{N}^{120}_{chase}, \\ &\bar{E}^{24}_{chase}, \bar{E}^{48}_{chase}, \bar{E}^{72}_{chase}, \bar{E}^{120}_{chase}, \\ &\overline{E^{24}_{chase}/N^{24}_{chase}}, \overline{E^{48}_{chase}/N^{48}_{chase}}, \overline{E^{72}_{chase}/N^{72}_{chase}}, \overline{E^{120}_{chase}/N^{120}_{chase}}, \\ &\bar{P}^{24}, \bar{P}^{120}, S_{cs}\}, \end{aligned} \tag{29}$$

- $\bar{E}^t_{chase}$ is the mean EdU number over assays 1 and 2 at time $t$ in the chase data.
- $\bar{N}^t_{chase}$ is the mean nucleoid number over assays 1 and 2 at time $t$ in the chase data.
- $\overline{E^t_{chase}/N^t_{chase}}$ is the mean fraction of EdU tagged nucleoids over assays 1 and 2 at time $t$ in the chase data
- $\bar{P}^t$ is the mean singly tagged proportion over all 3 assays as inferred from the intensity data in Section 3.1.3 at time $T$.
- $S_{cs} = 0$ is the control strength statistic, as introduced in Section 3.2.1 (for which the simulated version $\hat{S}_{cs}$ will be introduced in Section 3.3).

We did not include nucleoid number as a summary statistic for the pulse data. It must however be included for the pulse-chase data, as the system is no longer in equilibrium, and the rate at which the nucleoid number decreases informs us about how uncoupled the birth and death rates are. The fractions $E/N$ are not vital to include, but we found slightly better (qualitatively) posterior predictive plots when we did include them, as they incorporate information on how $E$ and $N$ covary. These division statistics are unnecessary to include for the pulse data since nucleoid number does not vary, and so the statistic $E/N$ includes no more information than the statistic $E$. Again, higher order moments were not deemed to be necessary to include.

##### 3.3 Initialisation, Burn-In, and $S_{cs}$

Let $C_\gamma^t = (N_\gamma^t, E_\gamma^t, l_\gamma^t)$ be a real cell from experiment $\gamma$ ($\gamma = pulse$, for the pulse cells, or $chase$ for the pulse-chase cells) measured at time $t$, consisting of nucleoid number, EdU tagged nucleoid number, and mitochondrial volume, respectively. For every cell $C_\gamma^t$, we simulate an analogous cell $\hat{C}_\gamma^t(s) = (\hat{N}_\gamma^t(s), \hat{E}_\gamma^t(s), \hat{l}_\gamma^t(s))$ over time $s \in (-t_b, t)$, where $t_b$ is some burn-in time (Section 3.3.2).

###### 3.3.1 Initialisation

For the pulse experiment, we use the state of the real cell $C_{pulse}^t = (N_{pulse}^t, E_{pulse}^t, l_{pulse}^t)$ to initialise the analogous simulated cell: $\hat{C}_{pulse}^t(-t_b) := (N_{pulse}^t, 0, l_{pulse}^t)$. That is, the cell starts with the same nucleoid number and mitochondrial volume as its associated cell, but with no mtEdU. For the pulse-chase experiment, this initialisation is not possible, as for time points $t > 24$ the nucleoid number $N_{chase}^t$ is out of equilibrium and is thus different to what the state of the cell was at the start of the experiment. Thus, we initialise based on the empirical distribution of $(N, l)$ inferred in Section 3.1.2. That is, if the simulation is being run with parameters $\beta_0, \beta_1, \sigma^2$, we initialise the simulated pulse-chase cell via $\hat{C}_{chase}^t(-t_b) = (\beta_0 + \beta_1 l_{chase}^t + N(0, \sigma^2 l_{chase}^t), 0, l_{chase}^t)$. In other words, the cell starts with the same mitochondrial volume as the real cell, no mtEdU, and a nucleoid number drawn from the inferred distribution of section 3.1.2.

For the two- and three-population models, we must also initialise the number of nucleoids in each subpopulation. For this we use the proportions $f_\alpha$ calculated previously (see Equations 12, 13 of Section 2.2.1 and Equation 18 of Section 2.3.1). That is, if a simulated cell has been initialised with nucleoid number $N$, then the number of nucleoids in subpopulation $\mathcal{N}_\alpha$ will be $N_\alpha = f_\alpha N$, where $\alpha = r, y, o$. Finally, we assume that the mitochondrial volume of the cells do not change over time, so that $\hat{l}_\gamma^t(s) \equiv l_\gamma^t$ is constant for a simulated cell and equal to the mitochondrial volume of its associated real cell.

###### 3.3.2 Burn-in

For $s \in [-t_b, 0]$, we do not allow EdU to accumulate in the simulated cells. In other words, for this time period, we simulate based on the dynamics of Equations 5, 8 or 14 for the one-, two-, and three-population models respectively. From this 'burn-in period', we determine the simulated control strength summary statistic $\hat{S}_{cs}$ mentioned in Section 3.2. This summary statistic is necessary specifically for accurate inference of the control strength statistic $c$. A value of $c$ which is too large will lead to the simulated cells to be too concentrated around the equilibrium line $N_{opt} = \beta_0 + \beta_1 l$ (Supplementary Fig. 7a). A value of $c$ which is too small will lead to the simulated cells to be distributed too far away from the equilibrium line (Supplementary Fig. 7a). We wish to build a summary statistic $\hat{S}_{cs}$ to quantify this behaviour. The statistic will have the property that $\hat{S}_{cs} > 0$ if $c$ is too small and the simulated cells are diffusing too far from the equilibrium line, $\hat{S}_{cs} < 0$ if $c$ is too large and the cells are too concentrated about the equilibrium line, and $\hat{S}_{cs} \approx 0$ if $c$ is appropriately specified.

To that end, let $\hat{R}(s) = \log[\hat{N}(s)] - \log[\beta_0 + \beta_1 \hat{l}]$ be the log-residual of a simulated cell at time $s$ (where we have dropped the sub and superscripts $\gamma$ and $t$). Due to the linear heteroscedasticity present in the data (see Section 3.1.2), applying logarithms is a well-known variance stabilising transformation in this case (see for example Bartlett [18]), and means that the quantity $\hat{R}(s)$ should be heteroscedastic (i.e., has constant variance regardless of $l$). Let $\mathcal{R}(s) = \{\hat{R}_i(s)\}_{i=1}^n$ be the set of log-residuals of each simulated cell (for either the pulse, or the pulse-chase experiment, depending on which inference we are performing). If $c$ is appropriately specified, then the sample variance of this quantity $\mathcal{R}(s)$ should be roughly constant through time, and in particular should not change much between the start and the end of the burn-in period: $\widehat{\text{Var}}[\mathcal{R}(-t_b)] \approx \widehat{\text{Var}}[\mathcal{R}(0)]$. It is reasonable to penalise a parametrisation equally if it leads to a factor of $k$ increase in this variance $\widehat{\text{Var}}[\mathcal{R}(-t_b)]/\widehat{\text{Var}}[\mathcal{R}(0)] = k$ vs a factor of $k$ decrease $\widehat{\text{Var}}[\mathcal{R}(-t_b)]/\widehat{\text{Var}}[\mathcal{R}(0)] = 1/k$, which leads us to the following choice of control-strength (cs) summary statistic:

$$\hat{S}_{cs} = \log \frac{\widehat{\text{Var}}[\mathcal{R}(-t_b)]}{\widehat{\text{Var}}[\mathcal{R}(0)]}$$

It is important to note that although we have labelled $t_b$ the 'burn-in time', the system does not need to be fully burned-in and decoupled from its initial conditions for the control strength summary statistic to be effective. Rather, we need only pick $t_b$ large enough in order for $\widehat{\text{Var}}[\mathcal{R}(-t_b)]$ to be sufficiently different from $\widehat{\text{Var}}[\mathcal{R}(0)]$ in the case where $c$ is misspecified. In other words, if $c$ is too small for example, the cells need time to diffuse away from the equilibrium line enough for it to become apparent. On the other hand, choosing $t_b$ too large becomes computationally expensive. We found that setting $t_b = 250$ was enough to be able to clearly see when $c$ was misspecified (Supplementary Fig. 7b), while being computationally feasible.

#### 3.4 Distance Function

We use a Euclidean distance function normalised by the variance of each measured summary statistic:

$$d(S_\gamma, \hat{S}_\gamma) = \sum_i \left( \frac{(S_\gamma)_i - (\hat{S}_\gamma)_i}{\sigma_i} \right)^2,$$

where $S_\gamma$, $\hat{S}_\gamma$ are the observed and simulated summary statistics respectively for $\gamma = pulse$ or $chase$, and $\sigma_i^2$ is the variance of the $i$'th observed summary statistic. For the summary statistics $\bar{E}^t_\gamma$, $\gamma = pulse$ or $chase$, $\bar{N}^t_{chase}$, $\overline{E^t_{chase}/N^t_{chase}}$, we set $\sigma_i^2$ to be the variance in the mean. That is, $\sigma_i^2 = \widehat{Var}[E^t_\gamma]/n_t$, $\sigma_i^2 = \widehat{Var}[N^t_{chase}]/n_t$, $\sigma_i^2 = \widehat{Var}[E^t_{chase}/N^t_{chase}]/n_t$, respectively, where $n_t$ is the number of observed cells at time point $t$. For the singly tagged proportion summary statistics $P^t$, we let $\sigma_i^2 = \widehat{Var_i}[P^t_i]$ be the variance of the inferred proportion over the three assays. We use this rather than the variance in the mean as we only have three datapoints, and we wish to incorporate into the ABC the idea that this summary statistic is not precise and only needs to be roughly correct.

For the control strength summary statistic $S_{cs}$, we do not have access to multiple realisations, so we must find alternate means to probe its variance. To do this, like in Section 3.3.2, we compute the log-residuals, but of the data itself, not of the simulated data, for each time point of the pulse cells $\widehat{Var}[\mathcal{R}^t_{pulse}]$. We then compute a 'realisation' of the $S_{cs}$ by computing $\log \frac{\widehat{Var}[\mathcal{R}^{t_1}_{pulse}]}{\widehat{Var}[\mathcal{R}^{t_2}_{pulse}]}$ for two time points in the data $t_1, t_2$. This gives us 16 'realisations' of $S_{cs}$, for which we compute the sample variance to set equal to $\sigma_i^2$. This estimate is necessarily imperfect; the true summary statistic is a function of the same cell at different time points, rather than of two different cells, for instance. This estimate, however, is the best proxy we have available to us in absence of this data. We do not use the pulse-chase cells to constrain $c$ as these are not in equilibrium.

#### 3.5 Priors, Posteriors and Simulation Pipeline

##### 3.5.1 Pulse Cells Fitting

Following the burn in (Section 3.3), at simulation time $s = 0$, we simulate adding EdU into the system by switching to simulating based on the dynamics of Equation 6, 64, or 66, depending on the model we are simulating. The simulation of a cell $\hat{C}^t_{pulse}$ is terminated at time $s = t$, and the summary statistics discussed in Section 3.2.1 are recorded.

We fit four models to the pulse data: the one-population model, the preliminary two-population model (where $p = 1$), the two-population model, and the three-population model. Priors were generally chosen to be uniform or log-uniform, and are displayed in Table 1. Notable exceptions include the replication rate $\mu_r$, for which we put an inverse uniform prior, with the interpretation that $1/\mu_r$ corresponds to the time it takes for an mtDNA to replicate, and $\beta_0, \beta_1$, for which we use as prior the posterior distributions inferred in Section 3.1.2. $500,000$ simulations were run for each model, and the acceptance criterion $\epsilon$ was chosen so that 0.1% of prior draws were accepted, leading to 500 posterior draws.

##### 3.5.2 Pulse-Chase Cells Fitting

Like in the pulse data fitting of the previous Subsection 3.5.1, at simulation time $s = 0$ we simulate adding EdU into the system. At time $t = 24$, we simulate removing EdU from the system by switching to the dynamics of Equations 65 or 67 for the two- or three-population model respectively. When $s = 24$, since the cells become out of equilibrium, we also change the parametrisation of the models to account for this change in behaviour. To be clear then, for the pulse-chase cells, there are three stages to the inference:

1. **Burn-in phase**: When the time $s \in [-t_b, 0]$, we simulate the model with no EdU, according to parametrisation $\boldsymbol{\mu}_{pulse}$.
2. **Pulse phase**: When the time $s \in [0, 24]$, we simulate the model with EdU incorporation dynamics, according to parametrisation $\boldsymbol{\mu}_{pulse}$.
3. **Chase phase**: When the time $s \in [24, 120]$, we simulate the model with EdU removal dynamics, according to parametrisation $\boldsymbol{\mu}_{chase}$.

The vectors $\boldsymbol{\mu}_{pulse}$, $\boldsymbol{\mu}_{chase}$ constitute different parameters and must both be inferred, although we assume there is significant overlap between them. Since only the two- population and three-population models fit the pulse data (see Section 4), we only fit these models to the pulse-chase cells. For the three-population model, we have parameters:

$$\boldsymbol{\mu}_{pulse} = (\beta_0, \beta_1, p, \mu_r, \mu_b, \mu_a, \mu_d, c),$$

but since $\mu_a = \mu_b/p$ for copy number control (Equation 16), $\mu_a$ does not need to be inferred. For the chase portion, we only assume that $\mu_b$ and $\mu_a$ changes:

$$\boldsymbol{\mu}_{chase} = (\beta_0, \beta_1, p, \mu_r, \mu_b^{chase}, \mu_a^{chase}, \mu_d, c, c^{chase} = 0).$$

Indeed, as there is no copy number control during the chase portion, and in particular the nucleoid number is decreasing, we must have $\mu_a^{chase} \geq \mu_b^{chase}/p$, so at least one of $\mu_a$, $\mu_b$ or $p$ must change between the pulse and chase portions of the experiment. We found that $\mu_a$ and $\mu_b$ was the smallest number of parameters we had to change before the three-population model achieved a good fit to the data. In fact, we found that the data could be well explained by simply setting $\mu_b^{chase} = \mu_a^{chase} = 0$, as we will see in Section 4.

For the two-population model, we have parameters:

$$\boldsymbol{\mu}_{pulse} = (\beta_0, \beta_1, p, \mu_r, \mu_b, \mu_d, c),$$

with the copy number control relationship $\mu_b = p\mu_d$ eliminating the need to infer $\mu_b$. For a direct comparison with the three-population model, we allow two parameters to vary between the pulse and chase portions for the two-population model: $\mu_b$ and $\mu_d$. Along with this, we also do an inference where we allow three parameters to vary: $\mu_b$, $\mu_d$ and $p$, and we will see that this model still does not explain the data as well as the three-population model in Section 4. We refer to the former model simply as the two-population model, and the latter as the *variable-p* two-population model:

$$\begin{aligned} \boldsymbol{\mu}_{chase} &= (\beta_0, \beta_1, p, \mu_r, \mu_b^{chase}, \mu_d^{chase}, c, c^{chase} = 0) \text{ (Two-population)} \\ \boldsymbol{\mu}_{chase} &= (\beta_0, \beta_1, p^{chase}, \mu_r, \mu_b^{chase}, \mu_d^{chase}, c, c^{chase} = 0) \text{ (Variable-p two-population)} \end{aligned} \tag{30}$$

In the main text we only display the two-population model. Here we will also display the variable-p two-population model. For the chase portion of the experiment, we set the control strength $c = 0$,

| | One-population | Preliminary two-population | Two-population | Three-population |
|---|---|---|---|---|
| $\beta_0$ | Pulse cells posterior from Section 3.1.2 | | | |
| $\beta_1$ | Pulse cells posterior from Section 3.1.2 | | | |
| $c$ | $LogU(10^{-5}, 10^{-1})$ | $LogU(10^{-5}, 10^{-2})$ | $LogU(10^{-5}, 10^{-2})$ | $LogU(10^{-5}, 10^{-1})$ |
| $\mu_b$ | $LogU(0.005, 0.1)$ | $LogU(0.005, 0.1)$ | $LogU(0.005, 0.1)$ | $LogU(10^{-5}, 1)$ |
| $\mu_r$ | N.A. | $1/U(0.1, 10)$ | $1/U(0.1, 10)$ | $1/U(0.1, 10)$ |
| $p$ | N.A. | N.A. | $U(0, 1)$ | $U(0, 1)$ |
| $\mu_d$ | N.A. | N.A. | N.A. | $LogU(5 \cdot 10^{-3}, 10^{-1})$ |

**Table 1:** Priors for pulse cells ABC. *LogU* denotes a log-uniform distribution, $1/U$ denotes the reciprocal of a uniform distribution.

so that the birth rate function $\lambda_b = \mu_b$ is constant, since the out of equilibrium behaviour of the cells means we have no expectation that copy number control is being maintained.

We expect that the inferred parametrisations for $\boldsymbol{\mu}_{pulse}$ for the pulse-chase cells should be similar to the parametrisation inferred for the pulse cells. Therefore, as prior to these parameters (other than $\beta_0, \beta_1$), we take the posteriors inferred from the pulse cells with a small perturbation $\xi$ added as priors to these parameters. This perturbation is necessary for two reasons. The first is that the parametrisation of the pulse-chase cells need not be identical to the pulse cells, due to different experimental conditions, for example. We thus should leave room for slightly different ABC posteriors. The second is that we are propagating not a full posterior distribution from the pulse fit, but rather 500 accepted posterior samples, which will give poor estimates of distributional tails. This discrete sample is thus smoothed into a continuous distribution with the addition of the perturbation $\xi$, similar to ABC-SMC, for example[19].

The way $\xi$ was added was dependent on the scale on which the parameter varied. For instance, $\mu_d$ varies on a log scale, so the perturbation was added to the log-parameter: $\mu_b' = exp(\log[\mu_b] + \xi)$. Here, $\mu_b$ is a draw from the posterior inferred from the pulse cells, and $\mu_b'$ is the perturbed parameter which is a draw from the prior of the pulse-chase cells. For a parameter $\mu$, varying on a scale $f$ so that $f(\mu)$ is the parameter transformed to the scale it varies on, $\xi$ was chosen to be normally distributed with variance equal to $0.1\sigma^2_{f(\mu)}$, where $\sigma^2_{f(\mu)}$ is the pulse cells posterior variance of $f(\mu)$. Performing this procedure rather than simply putting uniform or log-uniform priors allows us to use information already inferred from the pulse cells, and speeds up the ABC by reducing the complexity of parameter space.

Priors on $\beta_0, \beta_1$, like in the pulse inference, were given by the posteriors inferred in Section 3.1.2. Priors on the parameters that changed in the chase section, like $\mu_b^{chase}$ were generally chosen to be uniform or log-uniform, with any conditions such as $\mu_a^{chase} > \mu_b^{chase}/p$ for the three-population model built in. When fitting the three-population model, we found that posteriors for $\mu_a^{chase}$ and $\mu_b^{chase}$ were accumulating significant mass towards the values closest to zero. We thus opted to implement a spike and slab prior setup for these parameters. Full details of prior parameters are given in Table 2. Like in the pulse-cells, $500,000$ simulations were performed for each model, and 500 parameters were accepted.

### 4 Model Selection, Interpretation, and Validation

Having fitted each model to our data using ABC, we move to interpreting and validating the output of the fit, beginning with posterior predictive plots and ABC model selection, moving to parameter posterior interpretations and finishing with a validation using our withheld third assay. In this section, we finally make concrete the intuitions of Section 1, and clearly differentiate between models based on different aspects of the data.

| | Variable-p two-population | Two-population | Three-population |
|---|---|---|---|
| $\beta_0$ | Pulse-chase cells posterior from Section 3.1.2 | | |
| $\beta_1$ | Pulse-chase cells posterior from Section 3.1.2 | | |
| $c$ | $exp(\log[c'] + \xi)$ | | |
| $\mu_b$ | $exp(\log[\mu_b'] + \xi)$ | | |
| $\mu_r$ | $\lvert \frac{1}{1/\mu_r' + \xi} \rvert$ | | |
| $p$ | $min(\lvert p' + \xi \rvert, 1)$ | | |
| $\mu_d$ | N.A. | N.A. | $exp(\log[\mu_d'] + \xi)$ |
| $\mu_b^{chase}$ | $Ber(0.5) \cdot LogU(10^{-6}, 10^{-2})$ | | |
| $\mu_d^{chase}$ | $\frac{\mu_b^{chase}}{p^{chase}} + LogU(10^{-6}, 1)$ | $\mu_b^{chase} + LogU(10^{-6}, 1)$ | N.A. |
| $p^{chase}$ | $U(0,1)$ | N.A. | N.A. |
| $\mu_a^{chase}$ | N.A. | N.A. | if $\mu_b^{chase} \neq 0$ : $\mu_a^{chase} \sim \frac{\mu_b^{chase}}{p} + LogU(10^{-6}, 1)$ if $\mu_b^{chase} = 0$ : $\mu_a^{chase} \sim Ber(0.5) \cdot LogU(10^{-6}, 1)$ |

**Table 2:** Priors for pulse-chase cells ABC. $LogU$ denotes a log-uniform distribution, $1/U$ denotes the reciprocal of a uniform distribution. For some parameter $\mu$, $\mu'$ denotes this same parameter sampled from the 500 accepted pulse cells parameters, i.e., a draw from the pulse cells posterior. $\xi$ denotes a perturbation of $\mu'$, with distribution $\xi \sim N(0, 0.1Var[f(\mu')])$, where $f$ is the scale on which $\mu'$ varies. Specifically, $f(\cdot) = \log(\cdot)$ for $c', \mu_b'$, $f(\cdot) = 1/\cdot$, for $\mu_r'$, and $f(\cdot) = \cdot$, for $p'$. $Z$, in general, denotes a spike and slab prior. The prior for $\mu_a^{chase}$ should be interpreted as $\mu_a^{chase} \sim \frac{\mu_b^{chase}}{p} + LogU(10^{-6}, 1)$ if $\mu_b^{chase} \neq 0$, otherwise we put a spike and slab prior with probability 0.5 that $\mu_a^{chase} = 0$, and probability 0.5 that $\mu_a^{chase} \sim LogU(10^{-6}, 1)$. The conditions $\mu_d^{chase} > \frac{\mu_b^{chase}}{p^{chase}}$, $\mu_d^{chase} > \mu_b^{chase}$, $\mu_a^{chase} > \frac{\mu_b^{chase}}{p}$, implicit in the priors of the variable-p two-population, two-population, and three-population models respectively are necessary to ensure copy number drops during the chase, rather than rises.

.

#### 4.1 Bulk Posterior Predictive Plots

First, we construct and inspect posterior predictive plots for each model to gain an intuitive understanding of the model fits. We begin the with all three assays of the pulse cells, displayed in Supplementary Fig. 8. Inspecting the nucleoid trajectories (Supplementary Fig. 8a), it is clear that every model but the one-population model is able to fit the data adequately. The failure of the one-population is due to the fact that the mtEdU number must necessarily start at 0, while the other models have no such constraint due to their replicating subpopulation $\mathcal{N}_r$ that gets tagged immediately. Next, inspecting the posterior predictive distributions of the 24 hour singly tagged proportion, we see that only the two- and three-population models fit (Supplementary Fig. 8b). Focussing only on the two- and preliminary two-population models, the former fits this aspect, whereas the latter does not. The only difference between these two models is that in the two-population model, a recently replicated molecule has some probability $q = 1 - p$ of replicating again, whereas the preliminary two-population model has no such feature. This makes the intuition of Section 1 concrete: we must have preferential replication in order to explain the quantity of nucleoids which are doubly tagged in the data. Finally, we see that all models seem to fit the control-strength statistic well (Supplementary Fig. 8c), indicating that the ABC has selected values of $c$ that maintain linear copy number control effectively.

It is worth pausing here and considering the influence of our Gaussian mixture modelling summary statistic $\bar{P}^{24}$ (see Section 3.1.3), since the preferential replication insight is heavily reliant on this. Indeed, if we had chosen two peaks rather than three for the 24 hour time point, or if the underlying mixture is not Gaussian but some other distribution, this estimate would be different. However, it is evident from Supplementary Fig. 8b that the models with no preferential replication (preliminary two-population, one-population models) necessarily predict that around 75% of nucleoids are singly tagged after 24 hours. This is a constraint inherent in the models if we fix the replication rate by the mtEdU increase, regardless of $\bar{P}^{24}$. On the other hand, the models with preferential replication (two-, three-population models) have the freedom to decouple the predicted final singly tagged proportion from the replication rate simply by varying the parameter $p$. Therefore, as long as our estimate of $\bar{P}^{24}$ is significantly smaller than 75% (which we intuitively see by looking at the assays in bulk in main text Figure 6g), the preferential replication models will still display a far better fit than the non-preferential replication models, even if the true value is not 33%, as we estimate it. In other words, we argue here that the model selection should be robust to moderate deviations in this summary statistic.

Overall, we see that the two- and three-population models fit the pulse data well, while the preliminary two- and one-population models fail. We now inspect the posterior predictive distributions of the two-, variable p two-, and three-population models fit to the three assays of the pulse-chase cells. Beginning with the nucleoid trajectories (Supplementary Fig. 9a), we see that the two-population model predicts that the mtEdU number should keep dropping, disagreeing with the data, and overshoots the nucleoid number curve. When inspecting the fraction of nucleoids which are tagged over time, it becomes even more clear that the two-population model is a poor fit (Supplementary Fig. 9b). When allowing $p$ to vary, the variable-p two-population model looks visually as though it does better on the nucleoid trajectories (Supplementary Fig. 9a), but when comparing the ratio it is apparent that this model also fails to fit the data fully (Supplementary Fig. 9b). On the other hand, the three-population model fits both these aspects (Supplementary Fig. 9, a and b). In other words, even when allowing three parameters to change from the pulse to the chase phases, the two-population model still displays a poorer fit than the three-population model with only two parameters varying. Inspecting the other summary statistics, all of the models seem to fit the singly tagged proportions adequately (Supplementary Fig. 9c), but the two- and variable-p two-population models seem to output a biased posterior predictive for the control strength summary statistic, indicating that the model fit is pushing into less suitable areas of parameter space, hinting at model misspecification (Supplementary Fig. 9d). The three-population model, on the other hand, fits this statistic well also. Overall, we see that the three-population model, in which nucleoids must age before they die, fit the chase data better than the two-population model, concretising our intuitions from Section 1.

To create such posterior predictive plots, for the pulse cells, we simulated every cell for 24 hours, regardless of the time point it was actually measured, and recorded the nucleoid state every 15 simulation minutes. This is so that we could observe a smooth trajectory over the course of the full 24 hours. For a particular posterior draw, we ran this simulation, and averaged across all cells to obtain one sample from the average trajectory posterior predictive. We then did this for all 500 of our posterior samples to obtain a full average trajectory posterior predictive distribution, for which we took the mean across all samples to obtain the posterior predictive mean. This same procedure was done for posterior predictive plots for the pulse-chase data fitting, except in this case every cell was simulated for 120 hours. Posterior predictive distributions of the control strength statistic $S_{cs}$ and the single tagged proportion summary statistics $P^t$ are estimated simply from the 500 samples outputted from the posterior predictive procedure just mentioned.

#### 4.2 ABC Model Comparison

To formalise the informal model selection of the previous section, we performed a Bayesian model selection for the pulse and the pulse-chase fitting[20]. For this we took the $500,000$ prior parameters and corresponding simulated summary statistics for each model, and concatenated all of the summary statistics, labelling which model produced them. We then applied the same rejection algorithm as before, only accepting 500 total parameters. We then recorded which proportion of these accepted parameters came from each model, and recorded this as the posterior distribution on the models. Since each model had $500,000$ prior parameter draws, this constitutes a uniform prior on each model. Results are shown in Supplementary Fig. 10, and agree with our intuitions from the posterior predictive plots. Following the pulse data fitting, there is no posterior probability placed on the one- or preliminary two-population models. Slightly more is placed on the two-population model than the three-population model due to the simplicity of the two-population model in the absence of other data. Following the pulse-chase data, essentially all of the posterior probability is placed on the three-population model, leading us to select the three-population model as our final model. While the model selection indeed selects the three-population model, this is not a surprise: all of the key insights behind this selection are contained in Supplementary Fig. 1 and Supplementary Fig. 2 which motivate the stack of models that we designed.

#### 4.3 Interpretation of Posterior Distributions

Focussing on the three-population model fit to the pulse data first (Supplementary Fig. 11d), we see that there is quite some degeneracy in the parameter space, and many parametrisations fit using the pulse dataset only. However, the ABC is still evidently finding distinct regions in parameter space, as can be seen clearly in the two dimensional marginals (Supplementary Fig. 12d), and can be viewed as two interpretable modes (see discussion in Section 8). This is to be expected: the two-population model already sufficed to fit the pulse data, so adding in another parameter in the three-population model necessarily increased the number of degrees of freedom to one larger than is necessary. This degeneracy substantially reduces when subsequently fitting the model to the chase data (Supplementary Fig. 13c), where we see reasonably sharp posteriors for all of the pulse-portion parameters. In particular, the parameters $\mu_b$, $p$ and $c$ become far more constrained when the pulse-chase dataset is accounted for (Supplementary Fig. 11d and Supplementary Fig. 13c). Given that the evidence for the three-population model is in the chase portion of the experiment, not the pulse, this behaviour is expected. One aspect of the two dimensional three-population pulse-chase posterior marginal (Supplementary Fig. 14c) is worth noting: there is significant posterior weight on $\mu_b^{chase} = 0$, and indeed, on $\mu_a^{chase} = 0$, and in fact, even when these parameters are non-zero, there is significant posterior weight on very small, essentially zero values. This leads to a natural interpretation - the three-population model explains the chase well by halting the initiation of any more replication and ageing and simply allowing the old population to deplete, without even needing to alter the degradation rate. The two-population models have no such natural interpretation, as we'll see.

We see in the one dimensional pulse data marginals (Supplementary Fig. 11) that the posteriors of $\beta_0, \beta_1$ are roughly the same as the priors. This is expected, as the priors for these parameters were given by a Bayesian linear regression fit (see Section 3.1.2), whose distributions were passed through the ABC framework in order to propagate the uncertainty rather than to re-infer the parameters: the data which could be used to constrain these priors was already used to constrain them. We see across all models that the posteriors of many parameters ($c$, $\mu_b$, $\mu_d$) do indeed vary across orders of magnitude, justifying our use of log-uniform priors. An exception to this, $\mu_r$, has an inverse-uniform prior with the interpretation that $1/\mu_r$ corresponds to the time it takes for a replication event to finish. We see that in the two models that fit the pulse data, this is roughly 5 hours, while the preliminary two-population seems to give an unbiological estimate of 10 hours - and would have estimated higher if our prior had allowed it. We suggest that this was an effort on the models part to ensure that the replicating subpopulation contained 10% of nucleoids, in order to explain the initial accumulation of mtEdU at the start of the pulse experiment (see Supplementary Fig. 1b). Indeed, if we assume the posterior of $\mu_d$ is set by the slope of the EdU curve in Supplementary Fig. 1b) to be $\mu_d \approx 0.01$ (Supplementary Fig. 11b), we must have $\mu_r \approx 0.1$ (Equation 12) to achieve this 10% figure, implying a 10 hour replication time. The fact that this model gives an unreasonable estimate in this regard can be taken as more evidence for preferential replication as $p = 1$ is too strong a constraint, and we do not have the same problem in the two-population model, since $\mu_d$ is inferred to be higher (Supplementary Fig. 11c), and so $\mu_r$ is also. The reason why $\mu_d$ is inferred to be higher here will be discussed in Section 4.3.1.

The one- and preliminary two-population models struggle to infer a reasonable posterior for the control strength $c$ (Supplementary Fig. 11), while the two-population model gives a reasonably peaked posterior around $\log_{10}(c) \approx -3.5$, as does the the three-population model around $\log_{10}(c) \approx -1.7$, once the degeneracy in the three-population model's posteriors when fit to the pulse data only has been removed by fitting to the pulse-chase data (Supplementary Fig. 13c). The reasonable estimates of the more complex models justifies our use of the control strength summary statistic $S_{cs}$ (Section 3.3), and suggests that the simpler models fail to infer $c$ due to them being poorly suited models for the data in general. The two-population model, which fits the pulse data, displays reasonably spiked posteriors for all inferred parameters (other than $\beta_0, \beta_1$, which is expected) $\mu_d$, $\mu_r$, $c$, justifying our choice of priors and their bounds. The one- and preliminary two-population models do not enjoy the same behaviour due to their poor calibration to the data, and the three-population model for the pulse data due to parameter degeneracy.

Moving to the one dimensional pulse-chase posterior marginals (Supplementary Fig. 13), we recall that the priors for many of the parameters are perturbations of the posteriors for the pulse fit, which may have already been tight, and thus we do not necessarily expect pulse-chase posteriors of these parameters to get much tighter. Nevertheless, as previously mentioned we see reasonably tight posteriors for all pulse-portion parameters of the three-population model ($p, \mu_d, \mu_b, \mu_r, c$), justifying our choice of prior support, and summary statistics. It is instructive to observe the parameter $p$ and $p^{(chase)}$ in the two- and variable-p two-population models, both for the pulse and pulse-chase fit (Supplementary Fig. 11c, Supplementary Fig. 13, a and b). It seems as though the ideal $p$ for the pulse portion is $p < 0.3$, whereas for the chase portion, the variable-p two-population model chooses $p^{(chase)} > 0.5$ in order to attempt to fit the data. This abrupt change in a biophysical parameter does not seem biologically reasonable, but even then, this model fails to fit the data as well as the three-population model. We see similar behaviour in the two-population model's pulse-chase posteriors, where the inference for the parameter $p$ is on the edge of the prior, as the model attempts to push it as large as possible for the chase portion whilst still being compatible with the pulse-portion. The three-population model, on the other hand, has no need to move in this way, with the posterior for $p$ lying within the bounds of the prior (Supplementary Fig. 13c), and as already discussed, displaying a biologically reasonable interpretation that the cell halts initiating replication or ageing events and simply allows its old population to deplete with rate $\mu_d$.

Overall, an inspection of the posterior distributions demonstrate that not only does the three-population model fit the chase data better than the two-population model, but also displays a more biologically reasonable and interpretable parametrisation. The same can be said for the two- and three-population models as opposed to the preliminary two-population model - estimating a more reasonable replication time $1/\mu_r$.

##### 4.3.1 Turnover Rate from Different Models

It is instructive to inspect the estimated turnover rate for the preliminary two-, two-, and three-population models (Supplementary Fig. 15). The turnover rate, here, is defined as the replication, or equivalently, degradation rate per nucleoid in the whole cell, rather than per nucleoid in the subpopulation. For the three-population model, for example, this is given by $\mu_d f_o$ or $\mu_r f_r$, where $f_\alpha$ is the proportion of molecules in subpopulation $\mathcal{N}_\alpha$ (Section 2.3.1). We see that the two- and three-population models, which contain preferential replication, give very similar posteriors, demonstrating robustness between these two models. The preliminary two-population model on the other hand (which has $p = 1$) estimates a value which is roughly half that of the other two models. The reason for this is simple: in the preferential replication models, it is mainly EdU tagged molecules that are replicating, as recently replicated molecules replicate again. On the other hand, in the preliminary two-population model, it is mainly untagged molecules replicating. Thus, in the former two models, a replication event leads to one more mtEdU in the cell, whereas in the latter, a replication event leads to two more mtEdU in the cell. This leads to two different interpretations of the mtEdU curve (Supplementary Fig. 1b). In the former, we should take the slope of the curve ($\sim 0.014$) as the replication rate, while in the latter, we should take this quantity divided by two as the replication rate ($\sim 0.007$). Indeed, these crude estimates roughly correspond to the inferred posterior distributions of each model. This also serves as an explanation for $\mu_d$ being inferred to be higher for the two-population model in comparison to the preliminary two-population model (Supplementary Fig. 11, b and c): the turnover rate is higher.

##### 4.3.2 Turnover Rate from the Literature

Using their two-population model on in vitro human dermal fibroblast cells, Brüser et al. estimate a turnover rate of $> 0.5$ per nucleoid per day, or $> 2\%$ per nucleoid per hour, slightly larger than our 1.5% estimate[5], however, as discussed in Section 1.5, care needs to be taken when interpreting this result due to the proliferative nature of their cells and the difficulty of fully accounting for this behaviour in their models. Stewart et al. observe that after exposure to ethidium bromide, nucleoids from human fibroblast cells mostly deplete on the order of 7 days (their Figure 1), a similar time scale to our ethidium bromide experiment (our main text Figure 5g)[21]. Interestingly, although not discussed, a more rapid initial drop followed by a slower subsequent drop in mtDNA is also observable in their ethidium bromide data (their Figure 1), similar to ours, corroborating our two-step degradation mechanism (our main text Figure 5, i and j). Other studies have given mtDNA half-life estimates of 2-4 days[22], or rates of $0.7 - 1.4\%$ per nucleoid per hour ($\frac{\log 2}{t_{1/2}}$), matching our estimate. Longer estimates have been obtained from studies in vivo, with half lives of mitochondria (not specifically mtDNA) of 8-23 days (or rates of $0.13 - 0.36\%$ per nucleoid per hour)[23], and estimates of mtDNA turnover frequencies of 10-20 days, (or $0.2 - 0.4\%$ per nucleoid per hour)[24]. In general, in vivo studies display less turnover than in vitro studies: likely reflecting culture conditions versus tissue environment differences.

#### 4.4 Single Cell Posterior Predictive Plots

Section 4.1 showed posterior predictive plots of averaged trajectories across all cells for every parameter value. Here, we take the chosen three-population model and compare posterior predictive distributions of individual simulated cells in comparison to the real cell they are based on, to ensure that the models do not only appear to fit in bulk, but do in fact fit the data on a single-cell level.

We do this for the cells of all three assays. Supplementary Fig. 16 displays the observed nucleoid and mtEdU number within individual cells for the pulse and pulse-chase data, with 95% credible intervals for each cell. Visually, the three-population model seems to give reasonable credible intervals, with no cell lying significantly outside of the credible intervals. A closer inspection however reveals slight dispersion issues. For instance, only 78% of observed mtEdU values lie within the 95% credible intervals for the pulse data (Supplementary Fig. 16b), meaning the data is overdispersed compared to our model, while 100% of observed mtDNA values lie within the credible intervals (Supplementary Fig. 16a), revealing underdispersion in the data.

The mtDNA underdispersion can be simply explained as being an artifact of the initialisation: simulated cells begin with the same mtDNA number as their associated observed cell (Section 3.3.1), and so they are very likely to be close. The mtEdU overdispersion however is not an artifact, and points to there being more sources of stochasticity in the observed cells than the simulated cells. We discuss how we may account for this in our models in depth in Section 6.5.

#### 4.5 Higher Order Moments

Since our ABC summary statistics only included means, we probed our model to see whether it was explaining the variance and covariance between these summary statistics appropriately. Supplementary Fig. 17 displays observed moments (mean, variance, covariance of nucleoid and mtEdU number at different time points) with their 95% credible intervals computed from the three-population model. We see that exactly 95% of datapoints lie within the 95% credible intervals, demonstrating that our model fits higher order moments well despite not being explicitly fit to them.

#### 4.6 Validation with Assay 3

Whilst posterior predictive plots were performed using all three assays for visualisation purposes, the ABC fit and ABC model selection were both done using only assays 1 and 2 of the pulse and pulse-chase cells, and all three assays of the mtEdU intensity data. Here, we use the third assay to validate the three-population model and demonstrate that its validation accuracy is comparable to its training accuracy. We first simulate cells of only the third assay and compare their posterior predictive plots to the observed third assay (Supplementary Fig. 18, a and b). We see, in general, good visual agreement between the model between the posterior predictive and the observed data. Next, we simulated the cells of assays 1 and 2 using all 500 accepted parameter draws, and for each parameter draw computed the ABC distance function (Section 3.4) to obtain a posterior predictive distance. We did this for both the pulse and pulse-chase cells, obtaining training distances for both datasets. We repeated this process, but using only the cells of assay 3, obtaining validation distances for both datasets. Results are displayed in Supplementary Fig. 18c. We see that the validation distance is slightly higher than the training distance, as we would expect, however, with large overlap between distributions both for the pulse and pulse-chase cells, indicating an acceptable discrepancy. Overall, this demonstrates that our model generalises well to the third assay and is not overfitting to the training data.

### 5 Mouse Macrophages, Degradation Data, and LC3

#### 5.1 Mouse Macrophages

##### 5.1.1 No LPS

We have obtained a posterior distribution on the replication rate of human fibroblast cells with a posterior mean of 1.5%. To see if this figure is comparable across cell types, we performed the same EdU incorporation experiment in the mouse macrophages, measuring nucleoid number, mtEdU number, cellular and mitochondrial volume over time in individual cells. We opt to fit this data using a deterministic version of the two-population model using frequentist methods, for simplicity. We use the two- rather than the three-population model due to its lack of degeneracy (Section 4.3) when fit to pulse data, and since its inferred replication rate does not differ from the three-population model (Section 4.3.1).

Let $M_o(t)$ be the number of untagged molecules in the nonreplicating population $\mathcal{N}_o$ at time $t$. Assuming equilibrium dynamics, the birth rate should satisfy $\lambda_b = \mu_b$ (Equations 1, 2), which in turn must be balanced with the degradation rate $\mu_b = p\mu_d$ (Equation 11). This gives us the following differential equation:

$$\begin{aligned}\frac{dM_o}{dt} &= -(\mu_b + \mu_d)M_o \\ &= -(1+p)\mu_d M_o,\end{aligned} \tag{31}$$

which is easily solved:

$$M_o(t) = M_o^0 e^{-(1+p)\mu_d t},$$

where $M_o^0$ is the initial nucleoid number in $\mathcal{N}_o$. Assuming the cell has total nucleoid number $N$, and again, we are in equilibrium, this implies that the number of mtEdU $E(t)$ satisfies:

$$E(t) = N - M_o^0 e^{-(1+p)\mu_d t},$$

or, letting $f_r$ be the fraction of molecules in the replicating subpopulation (Equation 13), and $\mu_d^p = (1+p)\mu_d$ we get the proportion:

$$\frac{E(t)}{N(t)} = 1 - (1 - f_r)e^{-\mu_d^p t}. \tag{32}$$

We can fit this curve to the data using non-linear least squares, inferring the parameters $f_r$ and $\mu_b^p$. We obtain parameter estimates $\hat{f}_r = 0.31$, $\hat{\mu}_b^p = 0.02$, with standard error covariance matrix:

$$\Sigma = \begin{pmatrix} \sigma_{f_r}^2 & \sigma_{f_r,\mu_b^p} \\ \sigma_{f_r,\mu_b^p} & \sigma_{\mu_b^p}^2 \end{pmatrix} = \begin{pmatrix} 4.37 \times 10^{-6} & -1.64 \times 10^{-5} \\ -1.64 \times 10^{-5} & 1.61 \times 10^{-4} \end{pmatrix}$$

The turnover rate is given by the quantity $\mu_{turn} = \mu_d(1 - f_r)$, as this is the death rate per nucleoid in the cell. This in turn is given by:

$$\mu_{turn} = \mu_b^p(1 - f_r)/(1+p) \tag{33}$$

We do not attempt to infer the parameter $p$ for this data using mtEdU intensity data. Rather, we assume that $p$ is equally probable to take any value between 0 and 1, or in other words, that the sampling distribution of our parameter estimate $\hat{p}$ is distributed like $U(0,1)$, and hence we have point estimate $\hat{p} = 0.5$, and standard variance $\sigma_p^2 = 1/12$. To propagate these uncertainties, we Taylor expand:

$$\mu_{turn}(f_r, \mu_b^p, p) \approx \mu_{turn}^0 + \frac{\partial \mu_{turn}}{\partial f_r} f_r + \frac{\partial \mu_{turn}}{\partial \mu_b^p}\mu_b^p + \frac{\partial \mu_{turn}}{\partial p} p \tag{34}$$

and take variances of both sides to obtain:

$$\sigma_{\mu_{turn}}^2 \approx \left|\frac{\partial \mu_{turn}}{\partial f_r}\right|^2 \sigma_{f_r}^2 + \left|\frac{\partial \mu_{turn}}{\partial \mu_b^p}\right|^2 \sigma_{\mu_b^p}^2 + \left|\frac{\partial \mu_{turn}}{\partial p}\right|^2 \sigma_p^2 + 2\frac{\partial \mu_{turn}}{\partial f_r}\frac{\partial \mu_{turn}}{\partial \mu_b^p}\sigma_{f_r,\mu_b^p}, \tag{35}$$

where here we have assumed that the covariance between $p$ and the other two parameters is 0. Using Equations 33 and 35, we finally get the estimate:

$$\mu_{turn} = 0.9 \pm 0.2\% \tag{36}$$

##### 5.1.2 LPS

The addition of LPS causes the cells to become out of equilibrium, so the analysis of the previous subsection does not hold. We find that this data is consistent with the deterministic two-population model with $\mu_b = \mu_d = 0$, but nonzero $\mu_r, p$, allowing, in other words, the replicating subpopulation to deplete whilst disallowing any new birth and death events. The differential equations for such a setup are given by:

$$\begin{aligned} \frac{dM_o}{dt} &= 0 \\ \frac{dE_r}{dt} &= -p\mu_r E_r \\ \frac{dE_o}{dt} &= (1-p)\mu_r E_r + 2p\mu_r E_r = (1+p)\mu_r E_r, \end{aligned} \tag{37}$$

where $M_o, E_r, E_o$ are the untagged nonreplicating, tagged replicating, and tagged nonreplicating numbers, respectively. It can be verified that these differential equations are solved by:

$$\begin{aligned} M_o(t) &= (1-f_r)N^0 \\ E_r(t) &= f_r N^0 e^{-p\mu_r t} \\ E_o(t) &= (1+1/p)f_r N^0(1-e^{-p\mu_r t}), \end{aligned} \tag{38}$$

where $N^0$ is the initial nucleoid number, and hence the overall nucleoid and mtEdU curve is given by:

$$\begin{aligned} N(t) &= N^0 + \frac{f_r N^0}{p}(1-e^{-p\mu_r t}) \\ E(t) &= f_r N^0(1+\frac{1}{p}(1-e^{-p\mu_r t})). \end{aligned} \tag{39}$$

We fit the parameters $(f_r, \mu_r, p, N^0)$ to these nucleoid and mtEdU curves using non-linear least squares to obtain both point estimates and a covariance error matrix of our parameter estimates. The turnover rate is then given by $\mu_{turn}(t) = \mu_r E_r(t) = \mu_r f_r N^0 e^{-p\mu_r t}$; in particular, it is a function of time, where the turnover rate decreases as the replicating subpopulation depletes. We can plot this curve using our parameter estimates, and compute the standard error as a function of $t$ using identical methodology as Section 5.1.1 to obtain Extended Data Fig. 6j.

#### 5.2 Degradation Data Fit

The degradation data provides a unique form of evidence for the three-population model over the two-population model, in that the exponential decay we observe is better fitted by the former model. If replication is halted, the replicating subpopulation $\mathcal{N}_r$ must have depleted all of its molecules. In the two-population model, this leaves only the population $\mathcal{N}_o$, while in the three-population model, we have the two populations $\mathcal{N}_y$, $\mathcal{N}_o$ remaining. In the two-population model, the only Poisson process left is the degradation process $n_o \xrightarrow{\mu_d} \phi$. Under a deterministic treatment, this leads to the differential equation:

$$\frac{dN}{dt} = -\mu_d N,$$

solved by an exponential decay:

$$N(t) = N^0 e^{-\mu_d t},$$

where $N^0$ is the initial nucleoid number. The three-population model predicts a slightly different decay however, as we have two Poisson processes remaining: the ageing process $n_y \xrightarrow{\mu_a} n_o$, and the degradation process $n_o \xrightarrow{\mu_d} \phi$. Deterministically, this leads to the differential equation:

$$\begin{aligned} \frac{dN_y}{dt} &= -\mu_a N_y \\ \frac{dN_o}{dt} &= \mu_a N_y - \mu_d N_o. \end{aligned} \tag{40}$$

Assuming $\mu_d \neq \mu_a$, it can be verified that this differential equation is solved by:

$$\begin{aligned} N_y(t) &= N_y^0 e^{-\mu_a t} \\ N_o(t) &= (N_o^0 - \frac{\mu_a}{\mu_d - \mu_a} N_y^0) e^{-\mu_d t} + \frac{\mu_a}{\mu_d - \mu_a} N_y^0 e^{-\mu_a t}, \end{aligned} \tag{41}$$

where $N_y^0$, $N_o^0$ are the initial young and old nucleoid numbers respectively. Letting $f_o = N_o^0/N^0$ be the proportion of nucleoids in the old population initially, and $N = N_y + N_o$ the total nucleoid number, we obtain:

$$\begin{aligned} N(t) &= (N_o^0 - \frac{\mu_a}{\mu_d - \mu_a} N_y^0) e^{-\mu_d t} + \frac{\mu_d}{\mu_d - \mu_a} N_y^0 e^{-\mu_a t} \\ &= (f_o - \frac{\mu_a}{\mu_d - \mu_a}[1 - f_o])) N^0 e^{-\mu_d t} + \frac{\mu_d}{\mu_d - \mu_a}[1 - f_o] N^0 e^{-\mu_a t} \\ &= \frac{N^0}{\mu_d - \mu_a}[(\mu_d f_o - \mu_a) e^{-\mu_d t} + \mu_d (1 - f_o) e^{-\mu_a t}] \end{aligned} \tag{42}$$

We see therefore that the three-population model predicts a decay which is the sum of two exponential decays, not a single exponential decay. We fit both the ordinary exponential decay of the two-population model and the summed exponential decay of the three-population model to the degradation data. For the former, we infer the parameters $(N^0, \mu_d)$, while for the latter we infer $(\mu_a, \mu_d, f_o, N_0)$, both using nonlinear least squares. From this we obtain Fig. 5j of the main text, clearly showing a better fit from the deterministic three-population model.

An important caveat here is that we have had to re-infer the parameters $f_o, \mu_a, \mu_d$ for the degradation dataset. We get point estimates of $f_o = 0.37 \pm 0.04$, $\mu_a = 0.009 \pm 0.001$, $\mu_d = 0.33 \pm 0.1$, while an inspection of the posteriors of these parameters from the pulse-chase three-population ABC fit (Supplementary Fig. 11c, main text Fig. 6o) reveals posterior means of $f_o = 0.75$, $\mu_a = 0.2$, $\mu_d = 0.025$. A change in the parameters $\mu_a, \mu_d$ is expected due to the toxicity of ethidium bromide drastically altering the cellular dynamics, and we can interpret the increase in the degradation rate, for example, as reflecting this toxicity. We interpret the change in the old fraction from $f_o = 0.75$ to $f_o = 0.37$ as not a 'true' change, but rather reflecting a model oversimplification where we have assumed that the old and young subpopulations are distinct, where one can degrade and the other cannot. In reality, nucleoids will likely lie on a spectrum of more likely or less likely to degrade, and so we interpret this 37% figure as those molecules most likely to degrade, which are hence degraded very quickly under the conditions induced by the ethidium bromide. The 75% figure on the other hand reflects molecules which can degrade, including those which are not most likely to.

#### 5.3 LC3 Puncta

In the main text we showed that an average of $N_l = 14.22 \pm 3.13$ nucleoids are enveloped by LC3-positive vesicles at any given time, and hence in the process of degrading, and that there are $\eta_b = 13.5 \pm 2$ replication events per hour in the average cell. Suppose that a proportion of $f$ nucleoids are degraded through this pathway. The global degradation rate must balance the global replication rate, so that the global degradation rate is also $\eta_b$, and since a proportion of $f$ nucleoids are degraded through this pathway, we have a global degradation rate of $f\eta_b$ through this pathway. Hence, we have a per-capita degradation rate of $\frac{f\eta_b}{N_l}$ through this pathway. Finally, this implies that a nucleoid in this pathway would take on average $\frac{N_l}{f\eta_b}$ hours to degrade. If $f = 1$, i.e., every nucleoid degrades through this pathway, it follows from a simple uncertainty propagation that this would take $57 \pm 16$ minutes, roughly in accord with prior work on rat kidney cells[25]. The smaller $f$ is, the larger, and more biologically unreasonable, this estimate becomes, in an inversely proportional manner. For instance, selecting $f = 0.5$ leads to an estimate of $\frac{N_l}{0.5\eta_b} \approx 2$ hours, leading us to conclude that the majority of nucleoids are degraded through this pathway, in agreement with past observations of selective enrichment of nucleoids in autophagosomes in neurons[26].

### 6 Birth Rate Comparison

Here, we provide four birth rate functions which fit the form of Equation 2, including a birth rate that arises from hypothetical biological principles (Section 6.1), and perform an ABC to select between them (Section 6.2). We follow this by presenting two models which further improve on these birth rates by incorporating additional sources of stochasticity (Sections 6.4 and 6.5). Since these birth rate mechanisms are only in effect when the cells are in steady state, we focus on the pulse dataset in this section.

#### 6.1 A Biologically Motivated Mechanism: The Inhibition Model

We have already introduced differential, ratiometric, and logarithmic birth, and we reintroduce them here:

$$
\begin{aligned}
\lambda_b^{dif}(N,l) &= \mu_b + c(N_{opt}(l) - N) \text{ (Differential control)} \\
\lambda_b^{rat}(N,l) &= \mu_b + c\left(\frac{N_{opt}(l)}{N} - 1\right) \text{ (Ratiometric control)} \\
\lambda_b^{log}(N,l) &= \mu_b + c\left(\frac{\log(N_{opt}(l)/\beta_0)}{\log(N/\beta_0)} - 1\right), \text{ (Logarithmic control)}
\end{aligned}
\tag{43}
$$

where $N_{opt}(l) = \beta_0 + \beta_1 l$ as before. All of the above rates will adequately maintain linear copy number control, and adjusting the control strength $c$ will adjust the strength of this copy number control. It must be noted here that whenever we simulate using one of these birth rates, we enforce the condition that $\lambda_b \geq 0$, that is, we actually simulate with birth rate equal to $\max(0, \lambda_b)$ to ensure that the birth rate is never negative.

It can be argued that the functional choice of the above birth rates, while reasonable, have no microscopic justification. Here we present a hypothetical biological mechanism that gives rise to a birth rate mechanism satisfying the constraints of Equation 2. Suppose that when two nucleoids are near enough to each other, they inhibit each other from replicating. If this is the case, then if $N > N_{opt}$, the nucleoids will have a higher concentration within the mitochondrial network, and thus there will be more nucleoids inhibiting each other, leading to a lower birth rate. Likewise, if $N < N_{opt}$, the reverse happens: less nucleoids are inhibited, and the birth rate increases.

We model this as follows, working in a one-population framework where every molecule may replicate for now. Suppose that nucleoids within a distance $l_0$ from one another inhibit each other from replicating. This could correspond to inter-cristae space, for instance, where two nucleoids being separated by a cristae results in a large enough separation for no inhibition to occur, while two nucleoids occupying the same inter-cristae space results in inhibition due to their close proximity. We split the mitochondrial network into a series of $k$ compartments, where $k = l/l_0$. If two nucleoids are in the same compartment, then neither can replicate, while if a nucleoid is in its own compartment, it is free to replicate. Assuming that nucleoids are uniformly distributed across the network, let $(X_1,\dots,X_k) \sim$ multinomial$(1/k, N)$ be the random variables denoting the number of nucleoids in compartment $1,\dots,k$. Let $Y = \sum_{i=1}^{k} \mathbb{1}[X_i = 1]$ be the number of compartments with exactly one nucleoid, i.e., the number of nucleoids permitted to replicate with replication rate $\tilde{c}$. Then the average global replication rate is given by:

$$\tilde{c}\sum_{j=1}^{k} j\mathbb{P}(Y = j) = \tilde{c}\mathbb{E}[Y].$$

But this expected value can be computed simply:

$$\mathbb{E}[Y] = \sum_{i=1}^{k} \mathbb{E}[\mathbb{1}[X_i = 1]] = \sum_{i=1}^{k} \mathbb{P}(X_i = 1).$$

Now since $X_i \sim \text{Bin}(1/k, N)$,

$$\mathbb{E}[Y] = \sum_{i=1}^{k} \frac{N}{k}(1-\frac{1}{k})^{N-1} = N(1-\frac{1}{k})^{N-1},$$

The global replication rate is given by $\tilde{c}\mathbb{E}[Y]$, and so the per-capita rate is given by $\lambda_b^{inh} = \frac{\tilde{c}\mathbb{E}[Y]}{N}$, giving us:

$$\lambda_b^{inhib} = \tilde{c}(1-\frac{1}{k})^{N-1} \tag{44}$$

In order for this birth rate to be valid, we must show that it has form $\lambda_b^{inh} = \mu_b + f(N, N_{opt})$, satisfying Equation 2, where in particular $N_{opt} = \beta_0 + \beta_1 l$. In other words, having selected a-priori parameters $\mu_b, \beta_0, \beta_1$, we should be able to fix parameters $\tilde{c}$ and $l_0$ in order to ensure $\lambda_b^{inh}$ satisfies Equation 2. Indeed it can be shown (see Appendix B) that $\lambda_b^{inh}$ does satisfy this form, satisfying Equation 2 for $N_{opt}(l) = \frac{1}{l_o}\log\left[\frac{\tilde{c}}{\mu_b}\right] l$. In order to obtain an intercept to this linear control, we need only conjecture that the individual nucleoid birth rate $\tilde{c}$ increases slightly when the mitochondrial volume is abnormally small $\tilde{c} = c(1+\alpha/l)$, where $c$ and $\alpha$ are constants. In this case, $\lambda_b^{inh}$ satisfies Equation 2 for:

$$N_{opt}(l) = \frac{\alpha}{l_0} + \frac{1}{l_0}\log\left[\frac{c}{\mu_b}\right] l \tag{45}$$

Picking any $c > \mu_b$, and setting $\alpha$ and $l_0$ appropriately as functions of $\beta_0, \beta_1, \mu_b, c$ then gives us the copy number control $N_{opt} = \beta_0 + \beta_1 l$ we require. This demonstrates how a birth rate function with the desired copy-number control may arise by specifying a particular microscopic mechanism. In the next subsection, we demonstrate that the functional form of the inhibition model does not change when we add in different subpopulations.

##### 6.1.1 Within a Three-Population Framework

Suppose now that any some fraction $f$ of nucleoids are permitted to replicate. Let $(N_y^1,\cdots,N_y^k) \sim$ multinomial$(1/k, fN)$ denote the number of molecules which can replicate in each compartment $1,\cdots,k$, while $(N_o^1,\cdots,N_o^k) \sim$ multinomial$(1/k,(1-f)N)$, be those molecules which cannot replicate (which, with abuse of notation from Section 2, can include in principle both the old population and the replicating population). We assume again that the position of nucleoids are independent of each other, and so $(N_y^1,\cdots,N_y^k)$ is independent of $(N_o^1,\cdots,N_o^k)$. We also make the modelling choice that any molecule, replicative or not, can inhibit a replicative molecule from replicating even if the molecule itself is not replicative. Given these assumptions, the quantity of interest now becomes the number of compartments with one replicative molecule:

$$Y = \sum_{i=1}^{k} \mathbb{1}[N_y^i = 1 \cap N_o^i = 0] \tag{46}$$

Like before, the average replication rate will be $\tilde{c}\mathbb{E}[Y]$, and we may compute:

$$\begin{aligned} \mathbb{E}[Y] &= \sum_{i=1}^{k} \mathbb{P}[N_y^i = 1 \cap N_o^i = 0] \\ &= \sum_{i=1}^{k} \mathbb{P}[N_y^i = 1]\mathbb{P}[N_o^i = 0], \end{aligned} \tag{47}$$

where the second equality follows by independence. But now, $N_y^i \sim \text{Bin}(1/k, fN)$, $N_o^i \sim \text{Bin}(1/k, (1-f)N)$, so:

$$\begin{aligned} \mathbb{E}[Y] &= \sum_{i=1}^{k} \frac{fN}{k}(1-\frac{1}{k})^{fN-1} \cdot (1-\frac{1}{k})^{(1-f)N} \\ &= \sum_{i=1}^{k} \frac{fN}{k}(1-\frac{1}{k})^{N-1} \\ &= fN(1-\frac{1}{k})^{N-1}. \end{aligned} \tag{48}$$

Thus, the per-capita replication rate (replication rate per replicative molecule) is given by:

$$\lambda_b^{inhib} = \frac{\tilde{c}\mathbb{E}[Y]}{fN} = \tilde{c}(1-\frac{1}{k})^{N-1}. \tag{49}$$

In other words, the addition of more subpopulations in our model does not alter the per-capita birth rate function in the inhibition model.

#### 6.2 Heteroscedasticity and Model Selection

Having introduced four different birth rate mechanisms, we now discuss how to select between them. The difference between these control mechanisms is subtle, and comes in the form of heteroscedasticity. Supplementary Fig. 19, a and b shows plots and residuals of $N$ against $l$ for simulated cells of each control mechanism compared to real cells. We see that as we go from right to left in the figure, the simulated heteroscedasticity increases until it most matches the data for logarithmic birth.

To formalise this and perform model selection between the birth rates, we construct a moving standard deviation summary statistic to account for this heteroscedasticity. For this, we compute the set of squared residuals of the pulse data $\{R_i\}_i = \{(N - N_{opt})_i^2\}_i$, and the set of squared residuals of the corresponding simulated cells $\{\hat{R}_i\}_i = \{(\hat{N} - N_{opt})_i^2\}_i$. Here, the linear regression coefficients of $N_{opt} = \hat{\beta}_0 + \hat{\beta}_1 l$ are computed from regressing the observed (unadjusted) nucleoid number against the observed mitochondrial volume. We then compute the square root of the moving average of these quantities, resulting in a moving standard deviation for both the data $\boldsymbol{\sigma} = \{\sigma_i\}_i$ and the simulation $\hat{\boldsymbol{\sigma}} = \{\hat{\sigma}_i\}_i$ (Supplementary Fig. 19c). We then construct the 'heteroscedasticity summary statistic' $S_h$ as the Euclidean distance between these two quantities:

$$\hat{S}_h = ||\boldsymbol{\sigma} - \hat{\boldsymbol{\sigma}}|| \tag{50}$$

It can be seen in Supplementary Fig. 19c that the moving variance $\hat{\boldsymbol{\sigma}}$ most matches that of the data as we move from right to left, confirming that this summary statistic is a good measure of heteroscedastic difference, and indicating heuristically that logarithmic control is the best birth rate in terms of this metric.

To formalise this further we perform an ABC on the pulse-data for the three-population model for each of the four birth rates discussed with the exact same methodology as outlined in Section 3, but with two differences. First, we include this summary statistic $\hat{S}_h$, with the observed version of this summary statistic being set to zero: $S_h = 0$. Second, we set the burn-in-time to be $t_b = 1000h$, as opposed to $t_b = 250h$ (see Section 3.3), in order for the simulated cells to move as close to its equilibrium heteroscedastic state as possible (as discussed in Section 3.3, setting $t_b = 250h$ does not necessarily burn-in the cell fully). For the variance of the summary statistic $S_h$ (which we require for the distance function (Section 3.4), we take a similar approach to the variance of the control strength summary statistic $S_{cs}$. Namely, we compute the moving standard deviation $\boldsymbol{\sigma}^t$ of the data for the four different time points (1, 3, 7, 24 hrs) of the pulse data, and compute 6 'realisations' of the summary statistic by computing $||\boldsymbol{\sigma}^{t_1} - \boldsymbol{\sigma}^{t_2}||$ for all pairs $t_1 \neq t_2$. We then compute the variance of these 6 values, but with a manually set zero mean (i.e., the sum of the squares of this quantity), to obtain the variance of the summary statistic for the ABC distance function.

Following this ABC, we perform an ABC model selection using the same methodology as Section 4.2, putting uniform priors on each of the four birth rate models, and selecting the model with the highest posterior probability (Supplementary Fig. 19d). We see that we do indeed select logarithmic birth as the best birth rate mechanism, justifying our use of this birth rate throughout the manuscript.

Finally, uniquely for the inhibition model, we can obtain a posterior for $l_0$, the compartment size (Supplementary Fig. 19e). We see that a significant proportion of the posterior mass is concentrated on small values $l_0 < 1\mu m^3$, corresponding to a compartment length of $< 1\mu m$. The length scale of these compartments are therefore compatible with those of inter-cristae space within the mitochondria, which has been estimated to be $\sim 0.1\mu m$[27]. We therefore may interpret the inhibition model as stating that if two or more nucleoids are inhabiting the same inter-cristae space, they inhibit each other from replicating.

#### 6.3 Mathematical Explanation for Heteroscedastic Differences

We have seen quantitative differences in the heteroscedastic properties of each birth rate function, and have exploited them in order to select a birth rate which best matches the data. We can explain these differences mathematically as follows. For a given mitochondrial volume $l$, suppose that the nucleoid number deviates away from equilibrium $N_{opt}$ by some small $\delta N$. The birth rate will change accordingly in order to penalise this deviation, and the change will be given by:

$$\begin{aligned} \delta\lambda_b &= \lambda_b(N_{opt} + \delta N, l) - \lambda_b(N_{opt}, l) \\ &= \left.\frac{d\lambda_b}{dN}\right|_{N_{opt}} \delta N + O(\delta N^2), \end{aligned} \tag{51}$$

where $O$ is 'big-O' notation. Ignoring contributions of $\delta N^2$, computing the derivative for each of the three birth functions at $N = N_{opt}$, and using the fact that $N$ and $l$ scale linearly with each other, so that $N = O(l)$, we get:

$$\begin{aligned} \delta\lambda_b^{dif} &= -c\delta N = O(-\delta N) \\ \delta\lambda_b^{rat} &= -\frac{c}{N}\delta N = O\left(-\frac{1}{l}\delta N\right) \\ \delta\lambda_b^{log} &= -\frac{c}{[N/\beta_0]\log[N/\beta_0]}\delta N = O\left(-\frac{1}{l\log l}\delta N\right) \end{aligned} \tag{52}$$

For the inhibition model, we have:

$$\left.\frac{d\lambda_b^{inhib}}{dN}\right|_{N_{opt}} = c(1-\frac{l_0}{l})^{N-1}\log(1-\frac{l_0}{l})\Big|_{N_{opt}}$$
$$= \mu_b \log(1-\frac{l_0}{l}) \tag{53}$$
$$\approx -\frac{\mu_b l_0}{l},$$

and so:

$$\delta\lambda_b^{inhib} = O\left(-\frac{1}{l}\delta N\right), \tag{54}$$

giving the same heteroscedasticity properties as ratiometric birth. Thus, for the same deviation $\delta N$, differential birth penalises equally independent of $l$, ratiometric and inhibition birth penalises a factor of $\sim 1/l$ less, and logarithmic birth penalises a factor of $\sim 1/(l\log l)$ less. This dependence on $l$ gives rise to the heteroscedasticity and subsequent posterior probabilities that we observe in Supplementary Fig. 19.

#### 6.4 A Stochastic Birth Rate

Although logarithmic birth performed best out of all selected birth rate functions, we can make other birth rates work similarly by adding in extra sources of noise. Here, we discuss an adjustment to our microscopic model, inhibition control, which allows it to achieve the same, if not better, performance as logarithmic control. To achieve this, we introduce a stochastic element into the function by adding to it a mean zero Ornstein-Uhlenbeck (OU) process:

$$\lambda_b^{stoc-inh}(t) = \mu_b X(t) + \lambda_b^{inh}, \text{ (Stochastic inhibition control)} \tag{55}$$

where $X(t)$ satisfies:

$$dX = -\theta X dt + \sqrt{2\theta}\sigma dW, \tag{56}$$

where $W$ is a Wiener process. Here, we have parameterised $X$ so that $\mathrm{Var}[X] = \sigma^2$, and $\mathrm{Corr}[X(t), S(s)] = e^{\theta|t-s|}$, so that the parameter $\sigma$ modulates the variance of the fluctuations, while $\theta$ modulates the speed of fluctuations. This stochastic process encodes the biological fact that any cellular control mechanism will not be perfect. Sometimes the birth rate will be larger or smaller than it should be due to random fluctuations in concentration of replication machinery, or in the context of the inhibition model, due to imperfect inhibition between molecules. This could also capture, for example, the fact that we assume mitochondrial volume is constant in our models when in reality it is not.

For simulation purposes, we first generate this OU process using a standard Euler-Maruyama discretisation, which we then input into our simulation, treating it as a non-random function of time. Next, to simulate the stochastic three-population model, we use an inhomogeneous Gillespie algorithm. For this we use a novel technique which introduces a 'fake event' with a propensity configured so that the total propensity of every event is again constant over time, transforming the simulator into what is essentially a homogeneous Gillespie again[28]. Supplementary Fig. 20a shows the output of such a model. We see visually that the output of such a model seems even better than logarithmic control. Performing an ABC and subsequent ABC model selection with this birth rate function however, using the same methodology as Section 6.2 and placing priors $logU(10^{-3}, 10^{-1}), U(0, 0.3)$ on $\theta$, $\sigma$, respectively, reveals no preference between logarithmic control and stochastic inhibition control (Supplementary Fig. 20c). When looking at the posterior for $\sigma$ (Supplementary Fig. 20d), we see that it is constrained by the prior, but in particular it fits the data well despite its constraint $\sigma < 0.3$, demonstrating that the added fluctuations need not be large to achieve the desired heteroscedasticity (the birth rate can fluctuate by less than 30% to achieve the desired heteroscedasticity). $\theta$ is also constrained by its prior (Supplementary Fig. 20d), but has in its support many possible values that work well. $\theta$ ranging from 0.001 to 0.1 implies an autocorrelation half life ranging between $(\log 2)/\theta \approx 7 - 700$ hours, meaning these fluctuations can be slow or fast; both give the desired effect. At minimum, this demonstrates that other birth rate functions other than logarithmic birth can be made to fit the data just as well, and in particular, we should not dismiss the inhibition microscopic control mechanism on heteroscedastic grounds.

#### 6.5 Increasing mtEdU Dispersion

In section 4.4, we saw that the observed mtEdU exhibited slightly more variance than the three-population model predicted. There are many ways of adding further stochasticity into our model to account for this, with one being presented in Section 6.4. The form of stochasticity in Section 6.4 served to increase the heteroscedasticity of the model, but here, we give an adjustment which serves to increase the variance of the mtEdU specifically.

Implicitly thus far, we have been assuming that every cell is parametrised identically. Suppose instead that there is some variation in the turnover dynamics from cell to cell: some cells replicate quicker and some slower. In this case, some cells will have larger values for $\mu_b, \mu_d, \mu_a$, representing quicker dynamics, while some cells will have smaller values: a hierarchical model. This would induce more variation in EdU, since the 'fast dynamics' cells will have replicated more than our original three-population model predicts, while the 'slow dynamics' cells will have replicated less. To model this, for each cell we draw a random positive mean-1 constant $\kappa \sim Gamma(1/\theta, \theta)$, and simulate this cell with parameters $\kappa\mu_b, \kappa\mu_d, \kappa\mu_a$. $\theta$ is now a new parameter to infer, and represents the cell to cell variance in turnover dynamics. We label this hierarchical model *dispersed logarithmic birth*. Hypothetically, we could uncouple the variation in the three parameters by drawing three independent $\kappa_b, \kappa_d, \kappa_a$ and multiplying to obtain $\kappa_b\mu_b, \kappa_d\mu_d, \kappa_a\mu_a$, giving us a more general model. We find that this added complexity is not necessary for our purposes however. We perform an ABC with this model, placing a $U(0, 1)$ prior on $\sqrt{\theta}$ (the standard deviation of $\kappa$), using the methodology of Section 3. To select for parameters which give good mtEdU spread, we compute the heteroscedasticity summary statistic (introduced in Section 6.2), except we compute this summary statistic between simulated and observed mtEdU for each time point, as well as between the simulated and observed mtDNA number. This gives 4 new mtEdU heteroscedasticity summary statistics on top of the mtDNA heteroscedasticity summary statistic. We compute the variance of each of these mtEdU heteroscedasticity summary statistics (necessary for the distance function, see Section 3.4) in the same way as in Section 6.2, except using the three assays as three different measurements, rather than the four time points.

Once fit, we recompute single cell pulse posterior predictive plots using this new model (Supplementary Fig. 21a). We find that the mtEdU dispersion issues from Section 4.4 vanish, with 93% of the observed mtEdU values lying within the 95% credible intervals, as opposed to the 78% observed from from logarithmic control (Section 4.4). This demonstrates how adding a simple additional source of stochasticity can fully characterise the observed data. Despite improving on the posterior predictive plots, performing ABC model selection between logarithmic birth and dispersed logarithmic birth reveals no preference between them (Supplementary Fig. 21b); perhaps due to our heteroscedasticity summary statistics not being sensitive enough to detect a measurable difference between the models. Comparing the posteriors of this model to those displayed for logarithmic birth (Supplementary Fig. 22a), we see that there is no discernable difference for the major parameters of interest (other than the control strength $c$, which we should expect). For biological predictive purposes therefore, for example quoting a replication rate or subpopulation proportions, the models give identical predictions. We therefore choose to not display this model in the main text due to its complexity, instead displaying logarithmic birth. It is also notable that the posterior favours

| $N \to V$ | $V \to N$ | $N \to l$ | $l \to N$ | $V \to l$ | $l \to V$ |
|---|---|---|---|---|---|
| 0.051 | 0.34 | 0.007* | 0.034* | 0.66 | 0.002* |

**Table 3:** Granger Causality F-Test p-Values

smaller values of $\theta$ (Supplementary Fig. 22b), demonstrating that the cell to cell variability need not be large to achieve the desired effect. Overall, this demonstrates that we can adjust our models to fully account for the data with one simple change, but it is unnecessary for the inference of most parameters of biological interest.

##### 6.6 Interpretations

To conclude this section, we have compared 4 birth rate functions, one of which arising from a microscopic model, finding that logarithmic birth achieved the best performance. We have, however, also explored two ways of improving on these deterministic birth rate functions by adding other elements of stochasticity. Section 6.4 showed that adding an OU-process to inhibition control can induce further heteroscedasticity, matching and possibly improving on logarithmic control, and Section 6.5 showed that introducing cell to cell variability into logarithmic control can increase mtEdU dispersion, aligning with the observed data. These two sources of stochasticity were added to inhibition and logarithmic control, respectively, however, they could theoretically be added to any of the birth rate functions, and we could reasonably expect that many or all of the deterministic functions would explain the data adequately when these forms of stochasticity are added. In particular, the microscopic model underlying inhibition control is not ruled out as a possible mechanism to explain copy number control, as it can be made to explain the data as well as any other birth rate. Nevertheless, if a simple model with no added stochasticity is needed to be fit, this section demonstrates that logarithmic birth is the best choice out of the birth rates we explored, and is hence our choice throughout our modelling.

#### 7 Causality

To explore the causal relationship between nucleoid copy number, mitochondrial volume, and cell volume, we performed two Granger causal tests on our 4D imaging data, and three conditional independence tests on our static pulse data.

##### 7.1 Time Series Analysis (4D Imaging Data)

Let $(N_t^i, l_t^i, V_t^i)$ be the triplet of nucleoid number, mitochondrial network volume, and cell volume in cell $i$ at time $t$. We have 19 cells measured at 13 time points (0 to 24 hours inclusive with 2 hour increments), so $t \in \{0, 1, \ldots, 12\}, i \in \{1, 2, \ldots, 19\}$. Stationarity testing is discussed in Section 7.1.3. We perform both Granger causality[29] and neural Granger causality[30], a nonlinear generalisation of Granger causality, on this data. For Granger causality, data was normalized to have mean 0 and variance 1, while for neural Granger causality, we normalize the data to be between 0 and 1 via a min-max normalisation: $A_t^i \to \frac{A_t^i - \min_{i,t} A_t^i}{\max_{i,t} A_t^i - \min_{i,t} A_t^i}$, for $A = N, l, V$, due to evidence of better performance in prior studies[31], along with the fact that we found that the causal networks inferred were more stable.

###### 7.1.1 Granger Causality

Let $(X, Y, Z)$ be some permutation of the variables $(X, Y, Z)$. To test whether $Z$ is conditionally Granger causal for $X$ given $Y$, we fit, through ordinary least squares, an autoregressive model of the following form:

$$X_t^i = \beta_0 + \sum_{k=1}^{h} \beta_k X_{t-k}^i + \sum_{k=1}^{h} \gamma_k Y_{t-k}^i + \sum_{k=1}^{h} \delta_k Z_{t-k}^i + \epsilon_t^i \tag{57}$$

We allow the lag $h$ to be different for different permutations $(X, Y, Z)$, and determine it through 19-fold cross validation, where each fold is the data from one of the 19 cells (Supplementary Fig. 23a). Once this model had been fit, we ran an F-test with null hypothesis $H_0 : \delta_k = 0, \forall k \in \{1, \ldots, h\}$, and alternate hypothesis $H_1 : \delta_k \neq 0$, for some $k \in \{1, \ldots, h\}$. If the null was rejected with $p < 0.05$, we concluded that $Z$ is conditionally Granger causal for $X$ given $Y$, and we place a causal link from $Z$ to $X$ on the causal graph. p-values are shown in Table 3 and the resultant causal graph are shown in main text Fig. 3e.

##### 7.1.2 Neural Granger Causality

Neural Granger causality is a nonlinear generalisation of Granger causality introduced by Tank et al.[30] which hopes to account for complex data with nonlinear dependencies. Since we do not a-priori expect linearity to hold in our data, we applied this method to confirm that our traditional Granger causality analysis did not neglect any nonlinearities. Specifically, Tank et al. propose fitting a nonlinear $g$:

$$\mathbf{X}_t = g(\mathbf{X}_{<t}) + \epsilon_t, \tag{58}$$

where $\mathbf{X}_t = (N_t, l_t, V_t)^T \in \mathbb{R}^3$, $\mathbf{X}_{<t} = [\mathbf{X}_{t-1}, \mathbf{X}_{t-2}, \ldots]$ denotes past values of $\mathbf{X}_t$, and $\epsilon_t \in \mathbb{R}^3$ is a mean-0 error term. Crucially, $g$ is chosen such that the $j$'th variable (ie the $j$'th component $\mathbf{X}_{t,j}$ of $\mathbf{X}_t$) only depends on the past values of the $i$'th variable (ie the $i$'th row $\mathbf{X}_{<t,[i,:]}$ of $\mathbf{X}_{<t}$) through a vector of parameters $\mathbf{W}_{i\to j}$. That is, if $\mathbf{W}_{i\to j} = \mathbf{0}$, then the past values of the $i$'th variable have no influence on the present value of the $j$'th variable, and we may say that variable $i$ is *non neural Granger causal* for variable $j$. When training this model through proximal gradient descent, we apply a form of l1 regularisation to this set of weights $\{\mathbf{W}_{i\to j}\}_{i,j=1}^3$, analagous to a group lasso[32], which penalises each element of $\mathbf{W}_{i\to j}$ as a group, hence encouraging the model to eliminate its dependency on variables that it considers to be redundant. In other words, when selecting series that neural Granger cause series $j$, we have loss function:

$$\sum_t (\boldsymbol{X}_{j,t} - g_j(\boldsymbol{X}_{<t}))^2 + \lambda \sum_i ||\boldsymbol{W}_{i\to j}||_2. \tag{59}$$

A larger value of $\lambda$ corresponds to a harsher penalisation, selecting sparser causal connections, while smaller values correspond to a weaker penalisation and many causal connections. For more detail see Tank et al.[30]

In our modelling, we pick $g$ to be an LSTM (Long-Short Term Memory)[33], a form of recurrent neural network, due to its ability to capture long-term dependencies in time series data, and its impressive performance on small datasets[30]. We set the context length to be $h = 6$, and hidden layer size to be $H = 5$, and identified three different causal graphs, with two, three and four causal connections respectively, from three different choices of regularisation parameter (Supplementary Fig. 23, b and c). To ensure that our network was robust to different context length and hidden layer size choices, we manually tuned 9 networks, one for each pair $(H, h)$ with $H = 3, 5, 10$, $h = 4, 6, 8$, selecting the regularisation parameter to allow four causal connections. We found that all 9 networks selected the same four causal connections (Supplementary Fig. 24).

With robustness demonstrated, we focus on our three causal diagrams outputted by our selected $(H, h) = (5, 6)$ network. To distinguish between the causal diagrams, we attempted a 19-fold cross validation (Supplementary Fig. 25), where each fold are the measurements of a single cell, analogous to the linear case (Supplementary Fig. 23a). For this cross validation, for a given causal graph, if this causal graph had no causal connection between $i \to j$, then we retrain the LSTM with no

| | $H_0 : V \perp N \mid l$ | | $H_0 : V \perp l \mid N$ | | $H_0 : l \perp N \mid V$ | |
|---|---|---|---|---|---|---|
| | p-Value | Test Statistic | p-Value | Test Statistic | p-Value | Test Statistic |
| GCM | 0.02 | 2.25 | 0.00 | 3.29 | 0.00 | 6.55 |
| KCIT | 0.01 | 19.8 | 0.00 | 86.3 | 0.00 | 285 |
| CMIknn | 0.00 | 0.068 | 0.00 | 0.078 | 0.00 | 0.20 |

**Table 4:** Conditional Independence Tests

regularisation, using an Adam optimizer[34], while demanding that $\mathbf{W}_{i \to j} = \mathbf{0}$ is fixed, strictly zero, and untrainable. This ensures that the networks trained in the cross-validation have the desired causal structure in-built. With this methodology, we trained 19 networks in tandem, where network $k$ was trained on all but the $k$'th cell, and used the $k$'th cell as a validation set. We then averaged the training and validation loss across all 19 networks, and recorded the minimum (across all epochs) averaged validation accuracy.

This cross validation was repeated for five causal structures of increasing complexity, two of which being the three learned causal graphs, one being the network with no causal connections, and the last being the network where everything is causally connected (Supplementary Fig. 25a). Finally, this process was repeated 20 times, so that we had 20 validation accuracies for each of the four causal networks. Due to limited data, this cross validation was inconclusive; in general, the validation loss increased as network complexity increased, indicating that the model tended to over fit more for more complex networks (Supplementary Fig. 25b).

###### 7.1.3 Stationarity

For the above methods, weak stationarity of the data is theoretically required. However, from our modelling, we expect that the autocorrelation, within $N_t^i$ at least, is very high, and it would take far longer than 24 hours to get an accurate picture of its long-term behaviour. Thus, traditional statistical stationarity tests were not feasible. To justify this assumption therefore, we turn to the biology of the system. We expect each cell in culture to be in its steady state, given that each cell is post-mitotic. Indeed, our stochastic models of Section 1 and 2, which fit the static pulse data very well, are all stationary, and in fact are strongly stationary given that there is no explicit time dependence in the dynamics. Further, when we aggregate each of the 19 cells, the distributions of $(N_t, l_t, V_t)$ do not not significantly change over time (see main text Fig. 3d). We thus assume stationarity holds in our analysis.

##### 7.2 Conditional Independence Testing

To complement the causal time series analysis, we performed conditional independence testing on all three assays of the static pulse data to see whether we could probe conditional independence structure of $(N, l, V)$. If $(X, Y, Z)$ is some permutation of $(N, l, V)$, we remove the causal arrow from $X$ to $Y$ if we find that $X \perp Y \mid Z$ ('$X$ is conditionally independent to $Y$ given $Z$'). Formally, for each permutation, we perform the hypothesis test $H_0 : X \perp Y \mid Z$, $H_1 : X \not\perp Y \mid Z$. We use three separate test statistics to this end: a kernel based approach (KCIT)[35], a conditional covariance based approach (GCM)[36], and an information theoretic based approach (CMIknn)[37].

Each of these tests have their own merits and flaws. First of all, it should be noted that conditional independence testing is an inherently difficult problem, and Shah et al prove in their 'no-free-lunch' theorem that if some test holds a level $\alpha$ (i.e., has type-1 error bounded by $\alpha$, typically $\alpha = 0.05$), then the power of the test over an alternate distribution is at most $\alpha$[36]. This inherent difficulty necessarily means that any conditional independence test must compromise and cannot work universally in all situations. GCM explicitly recognises this difficulty and resolves it by testing for something more akin to a conditional uncorrelation than a conditional independence using a kernel ridge regression (hence being a semiparametric test), thus allowing the test to be more powerful.

KCIT, which works by estimating the Hilbert-Schmidt norm of the partial-cross covariance operator, attempts to hold a level-$\alpha$ for a broader class of null distribution due to it being a nonparametric test, but must necessarily sacrifice its power to do so due to the no-free-lunch theorem. CMIknn, which estimates the conditional mutual information, should theoretically have the same issue as it is also nonparametric, however has been demonstrated to enjoy a higher power than KCIT in simulation studies[37].

Overall, the results of any particular test should be treated with care, and in light of this we choose to employ all three in an attempt to reach a more robust conclusion. Notably, the test statistics for all three methods may be used as a proxy for conditional independence, with a lower test statistic moving 'closer' to conditional independence. With this in mind, test statistics and p-values are displayed in Table 4. We see that while every p-value is significant, indicating conditional dependence between all variables, for every test, the test statistic is smallest for $V \perp N|l$. Our interpretation of this is that $N$ and $V$ are not completely conditionally independent, and have some degree of influence on each other, however, this degree of influence is smallest when compared to the indirect connection between $(V, l)$ and $(l, N)$. As discussed in the main text, this result that the $(N, V)$ connection is smallest held true not only for the fibroblast cells, but also for HeLa cells, COS-7 cells, and primary murine hepatocytes, for all tests but one of the nine (Extended Data Fig. 2).

##### 7.2.1 Implementation Details

We implemented GCM using the R package 'GeneralisedCovarianceMeasure'[36] and KCIT using R package 'CondIndTests'[38,35], both on CRAN, while for CMIknn, we used the python package 'tigramite'[37]. We made no additions to the source code for CMIknn. GCM requires a choice of regression method, for which we chose kernel ridge regression, and both GCM and KCIT require a regularisation parameter $\delta$ for the regression, along with kernel width $\sigma_Z$. KCIT further requires choices of kernel widths $\sigma_X, \sigma_Y$. For KCIT, we chose $\delta$ to be small as in Zhang et. al[35] ($\delta = 10^{-3}$) in order to minimise bias, whereas in GCM, $\delta$ was determined through cross validation as in the original source code[36]. All width parameters were chosen to be the median distance between $X$ and $Y$, as Strobl et. al[39] found this heuristic to work better than the original hyperparameters in the source code of Zhang et. al[35]. For a more detailed discussion of the median heuristic, see Garreau et. al[40]. The null distribution for KCIT was estimated using the Monte-Carlo methods implemented in CondIndTests, and all data was normalised to have mean 0 and variance 1 prior to analysis.

#### 7.3 Interpretations

Any individual test of this section should be treated with care. We have discussed the limitations of the conditional independence testing in Section 7.2, while for the Granger tests, linear Granger causality suffers from being a linear method, thus having the potential to miss nonlinear dependencies, and neural Granger causality may overfit to the data, for instance, due to its complexity and our small dataset. Despite this, we see agreement across all tests, including the three conditional independence tests and two Granger causality tests that the strength of the connection between nucleoid number and cellular volume is weak in comparison to the indirect connection through the mitochondrial volume. We also see agreement across cell types: although most tests were done on the pulse fibroblast data, the conditional independence tests display agreement among HeLa cells, COS-7 cells, and primary murine hepatocytes (Extended Data Fig. 2). This robustness across cell types and statistical tests in spite of each tests individual limitations gives us strong grounding for our hypothesis, that we have used liberally in our stochastic modelling, that the cell controls nucleoid density with respect to the mitochondrial volume, rather than the cellular volume.

This result has some precedent in the literature. As discussed in Section 1.6, Seel et al. observe experimentally in budding yeast cells that nucleoid number is fixed by mitochondrial volume, by analysing a mutant with a smaller mitochondrial volume[11]. However, they do not find the same result for mtDNA number, rather that mtDNA number is fixed by cellular volume. Specifically, when probing mutant cells with lower mitochondrial volume, they found that they had lower nucleoid number than wildtypes, but the same mtDNA copy number, or more mtDNA per nucleoid. Although one must take care to extrapolate from mutants to wildtypes, this result highlights the critical distinction between nucleoids and mtDNA in this section, and all causal conclusions here must be taken for nucleoids specifically rather than mtDNA.

### 8 Discussion

To conclude this document, we here give a summative overview of our modelling, and the biological conclusions and interpretations that can be drawn from them.

***Key Empirical Constraints***

We start by listing the key empirical insights that constrained our modelling choices. These are discussed at more length in Section 1 and Supplementary Figures 1 and 2.

1. Nucleoid number scales linearly with mitochondrial network and cellular volume (Supplementary Fig. 1).
2. Causal analysis suggests that nucleoid number is only directly linked to mitochondrial network volume (Supplementary Fig. 1).
3. Around 10% of nucleoids get tagged immediately at the start of the pulse (Supplementary Fig. 2b).
4. At the end of the pulse, most nucleoids have been doubly tagged (Supplementary Fig. 2d).
5. During the chase, (untagged) mtDNA levels fall more rapidly than mtEdU levels (Supplementary Fig. 2f).

We will be referring back to these constraints throughout this section, and any model we build must satisfy them.

***Three Populations***

Constraint 3 motivates the addition of a replicating subpopulation which gets tagged immediately upon the addition of EdU, moving from the one-population model to the preliminary two-population model. We saw that Constraint 3 was only well satisfied once this subpopulation was introduced (Supplementary Fig. 8a).

Constraint 4 necessitates the idea of preferential replication of newly replicated molecules, moving from the preliminary two-population model to the two-population model by adding a probability $q = 1 - p$ that a newly replicated molecule begins replicating again. This makes it more probable that more doubly tagged molecules will be seen, and we saw that Constraint 5 was only well satisfied once this preferential replication was introduced (preliminary two-population model fits less well, see Supplementary Fig. 8b).

Constraint 5 motivates the idea of preferential degradation of non-newly replicated molecules. We modelled this by splitting the non-replicating population into a young population, the elements of which could initiate replication but not degradation, and an old population, the elements of which could initiate degradation but not replication, and adding a rate of transfer from the young to the old population. This three-population model was the only model which could satisfy Constraint 5 (Supplementary Fig. 9, a and b). It is also noteworthy that the three-population model is the only model which can explain the rapid initial nucleoid decline followed by a gradual decline observed when replication was inhibited without needing time varying parameters (see Section 5.2).

***Possible Deviations***

We made a number of modelling choices to incorporate the above empirical constraints. Here we discuss how a real cell may deviate from them.

First, we incorporated two modelling assumptions: the young population is forbidden from degrading, and that the old population is forbidden from replicating, in order to satisfy Constraint 5. Due to only the young population replicating, at the end of the pulse, the majority of mtEdU were in the young population. Consequently, due to only the old population degrading, during the chase, the majority of nucleoids degrading were untagged. These two modelling assumptions therefore worked well in fitting the data. However, such rigid assumptions should not be strictly necessary to satisfy this constraint. The only essential facet for a good fit is that the rate of degradation for young molecules is significantly smaller than that of old, and likewise, that the rate of replication for old molecules is significantly smaller than that of young. This should achieve the same desired result of the majority of mtEdU being in the young population following the pulse, and the majority of degradation occuring on the old population during the chase. A real cell therefore, may not shield its subpopulations from replication and degradation respectively as stringently as our model assumes.

Similarly, we assume in our model that there is no rate of transfer from the old population to the young. Such a rate of transfer may exist in reality; however, if so, it must be small in order for Constraint 5 to remain satisfied. If this transfer rate from old to young is large (and correspondingly the ageing rate from young to old is large, to keep the subpopulation sizes fixed), then the old and young subpopulations will be very well mixed. This would mean that by the end of the pulse, rather than most mtEdU being in the young population, which was the key to the three-population model satisfying Constraint 5, the mtEdU would be well mixed throughout both the young and the old. This therefore would not satisfy Constraint 5.

Finally, we have assumed that the young and old subpopulations are distinct, discrete subpopulations within the cell. It is quite possible that there is a scale of gradation between them, with molecules which are ‘older’ being more likely to degrade and less likely to replicate, and those which are ‘younger’ being more likely to replicate and less likely to degrade. Although Constraint 3 required the replicating population to be distinct from the others, Constraint 5 requires no such condition, only that the majority of molecules that degrade are untagged, which can be achieved both by discrete means, as our model does, or by continuous means.

***Parameter Interpretations***

Main text Fig. 6p highlights that molecules take 4.8±1.0 hours to undergo one replication event. This figure is compatible with previous estimates of mtDNA replication. Most recent models of mtDNA replication suggest that the strands of mtDNA are replicated consecutively, not concurrently [12], and previous studies estimate that polymerase $\gamma$ takes 1.5 hours to replicate a single strand. Therefore, one can infer a replication time of $\sim$ 3 hours from the literature, which lies in the support of our posterior (see main text Fig. 6p). Main text Fig. 6p also highlights that molecules in the young and replicating populations age into the old population after a mean of $14.4 \pm 4.4$ hours (which go on to degrade after a mean of $40 \pm 4.6$ hours), meaning that these populations are short-lived. The old molecules, therefore, should not be interpreted as mtDNA which are months or years old, for example, but rather mtDNA that have not replicated recently, within the last 15 hours on average.

Fitting the three-population model to the pulse data revealed parameter degeneracy due to to it being under-constrained (Supplementary Fig. 11d, Supplementary Fig. 12d). Focussing on the joint posterior of $p$ and $\mu_d$ in Supplementary Fig. 12d (where $q = 1 - p$ is the probability that a newly replicated molecule begins replicating again, and $\mu_d$ is the pre-capita degradation rate from the old subpopulation), this degeneracy can be interpreted as containing two modes. The first is the ‘small $p$’ regime, where $p$ is constrained to be small and the per-capita degradation rate $\mu_d$ is large, and the 'unconstrained $p$' regime whereby $\mu_d$ is small, and $p$ is unconstrained. These two modes can be interpreted as two different ways the three-population model is able to satisfy Constraint 4. When $\mu_d$ is small, the number of molecules in the old subpopulation will be large at equilibrium, proportional to $1/\mu_d$ (see Equation 18). This is in order to maintain the same global degradation rate (the per-capita rate $\mu_d$ multiplied by the number of old molecules) to match the fixed global replication rate. Given this, there will be very few molecules in the young population in the 'unconstrained $p$' regime. Therefore, as soon as a molecule finishes replicating and moves from the replicating population to the young, it is very likely to begin replicating again immediately simply because there are not many other young molecules to choose from. This leads to the three-population model satisfying Constraint 4 regardless of the value of $p$, allowing it to be unconstrained. In other words, the 'old/young' distinction is replacing the role of parameter $p$ in ensuring a large amount of EdU is incorporated into a small number of molecules, which comes at the price of a very small replicative population: $14 \pm 2\%$ if we look at only those posterior samples with $p > 0.8$ (so that $p \approx 1$, and so repeated replication is not governed by $p$). In contrast, when $\mu_d$ is larger, the old and young subpopulations are more balanced, meaning Constraint 4 is no longer enforced by a small young population, and must instead be enforced by the repeated replication probability $q = 1 - p$.

Subsequently fitting the three-population model to the chase portion of the experiment breaks this degeneracy and selects the small $p$ regime as the favoured mode (Supplementary Fig. 13c, Supplementary Fig. 14c). This is because in order to satisfy Constraint 5, the young subpopulation must be large enough ($25 \pm 8\%$) to contain the majority of the mtEdU, with the old subpopulation containing the majority of the mtDNA, as discussed in the above subsection. Additionally, in the chase portion of the fit, significant posterior mass is placed on $\mu_b^{chase} = \mu_a^{chase} = 0$ (Supplementary Fig. 13c, Supplementary Fig. 14c), or in other words, Constraint 5 can be satisfied by simply switching off any new replication events from initiating, and any ageing events moving young molecules to old. In this case, the model simply allows the old population to deplete, keeping the per-capita degradation rate $\mu_d$ the same as in the pulse portion, and allows the replicating population to finish replicating the molecules already mid-replication. This offers a parsimonious explanation of our chase data using our three-population model.

***Subpopulations and Spatial Structure***

In order to act according to (or approximating) the dynamics of the three-population model, since the mtDNA do not have an in-built clock, the cell must have some other way of distinguishing between old and young molecules. In the main text, we present evidence that molecules replicate closer to the nucleus than the average molecule. This implies that nucleoids are degraded closer to the cellular periphery. It is possible that this is due simply to increased prevalence of replication machinery such as polymerase $\gamma$ closer to the nucleus. In this case, as we move further out to the cellular periphery, molecules become less likely to replicate due to decreased prevalence of replication machinery. Through similar biophysical means, it is possible that molecules closer to the cellular periphery are more likely to be chosen to degrade. This naturally leads to an interpretation of our young subpopulation gradually shifting into the old subpopulation as we move from the nucleus to the cellular periphery, or further away from the majority of replication machinery. In other words, the distinction between the two subpopulations needed for Constraint 5 may purely be spatial. Continuing with this interpretation, a molecule which has just been replicated will necessarily already have polymerase $\gamma$ and the other replication machinery that has just replicated it in its proximity. It being likely to replicate again could simply be due to the biophysical fact that this machinery is most likely to replicate the molecule it is closest to. This allows Constraint 4 to also be satisfied from spatial means.

Non-spatial aspects may also inform the dynamical process. MtDNA naturally exists in compactified (inaccessible) or non-compactified (accessible) states[4,5]. An accessible mtDNA may be more able to replicate, and would thus be in the young population, and likewise an inaccessible mtDNA may be less likely to replicate, and would be in the old population. In this case, the young and old subpopulations would indeed be distinct. It is worth noting in this case that Isaac et al. estimate that $\sim 20\%$ of nucleoids are in this accessible state[4], compatible with the three-population model replicative (young or replicating) proportion posterior of $25 \pm 8\%$. Recalling from the previous subsection that the pulse data constrained the three-population model to two dynamic modes, one with a smaller replicative subpopulation size of $14 \pm 2\%$, this reveals that the chase portion of the experiment confined the model to the more biologically reasonable mode, under this interpretation. Further, of every model we have discussed, only the three-population model is able to be interpreted in terms of mtDNA compaction. In the one-, preliminary two-, and two-population models, every mtDNA is able to replicate - there is no notion of accessibility. Mutational burden may also inform the dynamics. It is notable, for example, how quickly the turnover events occur, with young molecules ageing in only $14.4 \pm 4.4$ hours, and molecules turning over at a rate of $1.5 \pm 0.2\%$ per hour. This may be linked to mtDNA being exposed to reactive oxygen species[41].

***Feedback Terms***

Constraints 1 and 2 necessitate a feedback whereby the cell controls its nucleoid number content based on its mitochondrial network volume. Since the degradation rate appears to be constant per-capita, this feedback is likely in the birth rate (see Section 1.1). We here discuss the conclusions drawn from our exploration of such birth rates (Section 6). We found that of the four birth rates explored (Equations 43, 44), logarithmic control fit the data the best, due to the simulated cells exhibiting heteroscedasticity the most similar to the data. The mathematical form of this birth rate is contrived deliberately so that its derivative scales to achieve this desired heteroscedasticity (see Equation 52). Although this birth rate fit the best, we demonstrated that when other sources of stochasticity within the cell are accounted for (for instance, a mitochondrial network which is fluctuating in size, rather than constant as our models assume), other birth rate mechanisms can fit the data just as well, if not better, than logarithmic birth (Section 6.4). In particular, we give an example of a hypothetical biological mechanism whereby mtDNA inhibit other mtDNA from replicating if they are close to each other in proximity, from which we can derive a birth rate. This birth rate, when simulated alongside added stochasticity, performs on-par with logarithmic control. Moreover, we demonstrated that the length scale at which this inhibition takes place is compatible with the interpretation that two nucleoids inhibit each other's replication if they share the same inter-cristae space (see Supplementary Fig. 19e and end of Section 6.2). This mechanism, which we term the inhibition model, demonstrates that copy number control of the form we observe can arise from candidate biological principles, rather than having to be mathematically imposed. In general, due to different birth rates behaving similarly when stochasticity is added, it is difficult to concretely conclude which mechanism the cell follows. The most general mathematical conclusion we can reach is that the control mechanism, whether it have a stochastic component or not, must allow for more variance about the equilibrium line as the cell gets larger. Biologically, the copy number control becomes less tight as the mitochondrial volume grows.

***Causality***

Our causal modelling adds a further layer of complexity to the problem of selecting a copy-number control mechanism. Our exploration of causality concluded that nucleoid number is only directly causally linked to mitochondrial volume (see Section 7). Throughout our stochastic modelling, we have implicitly assumed a direction for this causality, specifically that the mitochondrial volume determines the nucleoid number. This has empirical grounding in previous studies[11] (see discussion in Section 7.3), however, although not explored in our stochastic models, it is possible that the opposite direction also holds, that is, that nucleoid number influences mitochondrial volume. Indeed, we observed that drastically altering the mtDNA content of a cell did cause a change in mitochondrial volume (main text Fig. 3i). This allows for the possibility of a complex bidirectional feedback between nucleoid number and mitochondrial volume, from which a more intricate copy number control mechanism than the ones we have explored could arise. Therefore, although we have explored many mechanisms here that fit the data well, elucidating precisely which of these mechanisms, if any, the cell truly obeys remains an open problem.

Overall, our modelling gives strong evidence that recently replicated molecules tend to replicate again, are less likely to degrade, and turnover is high: $1.5 \pm 0.2\%$ per hour. If less molecules are able to replicate, this creates an effective genetic population size which is smaller than the number of mtDNA in the cell. A smaller genetic population can lead to more rapid fixation of mtDNA mutations under neutral drift with implications for mtDNA accumulation with ageing[42,43,44,45].

### A Glossary of Key Terms and Symbols

- $\boldsymbol{N}$: Nucleoid number.
- $\boldsymbol{l}$: Mitochondrial network volume.
- $\boldsymbol{V}$: Cellular volume.
- $\boldsymbol{\mathcal{N}_\alpha}$: The subpopulation of nucleoids which are replicating, if $\alpha = r$; young, if $\alpha = y$; old, if $\alpha = o$; or remnant replicating, if $\alpha = rem$.
- $\boldsymbol{\mathcal{N}_\alpha^{(i)}}$: The subpopulation of $\mathcal{N}_\alpha$ of nucleoids which have $i$ EdU tags.
- $\boldsymbol{N_\alpha}$: The number of nucleoids in $\mathcal{N}_\alpha$.
- $\boldsymbol{N_\alpha^{(i)}}$: The number of nucleoids in $\mathcal{N}_\alpha^{(i)}$.
- $\boldsymbol{n_\alpha}$: A single nucleoid in $\mathcal{N}_\alpha$.
- $\boldsymbol{n_\alpha^{(i)}}$: A single nucleoid in $\mathcal{N}_\alpha^{(i)}$.
- $\boldsymbol{E^{(i)}}$: The number of nucleoids measured as having $i$ tags.
- $\boldsymbol{\lambda_b}$: The per-capita (per nucleoid in $\mathcal{N}_y$, for the three-population model) replication rate, of the form $\lambda_b = \mu_b + f(N, l)$, where $\mu_b$ is a constant, and $f$ is a function of $N$, $l$, referred to as a feedback law, satisfying Equation 2. In particular, $f(N, N_{opt}) = 0$, where $N_{opt} = \beta_0 + \beta_1 l$ is the optimal copy number within a cell, which scales linearly with mitochondrial network volume, and $\beta_0, \beta_1$ are constants.
- $\boldsymbol{\mu_b}$: The constant per-capita (per nucleoid in $\mathcal{N}_y$, for the three-population model) replication rate at equilibrium ($N = N_{opt} = \beta_0 + \beta_1 l$).
- $\boldsymbol{\beta_0, \beta_1}$. Constants dictating the linear equilibrium between nucleoid number and mitochondrial network volume within a cell: $N_{opt} = \beta_0 + \beta_1 l$.
- $\boldsymbol{c}$: The constant control strength. Parametrises the birth rate feedback law function $f$.
- $\boldsymbol{\mu_d}$: The constant per-capita (per nucleoid in $\mathcal{N}_o$, for the three-population model) degradation rate.
- $\boldsymbol{\mu_r}$: The constant per-capita (per nucleoid in $\mathcal{N}_r$, for the three-population model) replication-termination rate, i.e, the rate at which nucleoids which are currently replicating finish replicating.
- $\boldsymbol{\mu_a}$ The constant per-capita (per nucleoid in $\mathcal{N}_y$, for the three-population model) ageing rate, i.e, the rate at which nucleoids which are in $\mathcal{N}_y$ move to $\mathcal{N}_o$.
- $\boldsymbol{p}$: The probability that following a replication termination event, both daughter nucleoids move to $\mathcal{N}_y$. Otherwise, with probability $q = 1 - p$, one nucleoid remains in $\mathcal{N}_r$.
- $\boldsymbol{x^{chase}}$. The parameter $x$ from the pulse portion of the experiment ($x = \mu_b, \mu_d$, etc) now in the chase portion of the experiment, whose value may have changed (i.e, $x \neq x^{chase}$).
- **Three-population model**: Contains all three populations $\mathcal{N}_r$, $\mathcal{N}_y$, $\mathcal{N}_o$. Replication acts on $\mathcal{N}_y$, degradation acts on $\mathcal{N}_o$. There is preferential replication of newly replicated molecules, and preferential degradation of old molecules.
- **Two-population model**: Contains two populations $\mathcal{N}_r, \mathcal{N}_o$, with no young population, and no ageing. Both replication and degradation act on $\mathcal{N}_o$. There is preferential replication of newly replicated molecules, but no preferential degradation.

- **Preliminary two-population model**: The two-population model where $p$ is set to $p = 1$. There is neither preferential replication or preferential degradation.
- **One-population model**: There is only one population. The only events acting are replication and degradation with rates $\lambda_b$, $\mu_d$, respectively.
- **Differential Birth/Control**: Birth rate feedback law of the form $f(N, l) = c(N_{opt}(l) - N)$.
- **Ratiometric Birth/Control**: Birth rate feedback law of the form $f(N, l) = c\left(\frac{N_{opt}(l)}{N} - 1\right)$.
- **Logarithmic Birth/Control**: Birth rate feedback law of the form $f(N, l) = c\left(\frac{\log(N_{opt}(l)/\beta_0)}{\log(N/\beta_0)} - 1\right)$.
- **Inhibition Birth/Control**: Birth rate feedback law arising from a hypothetical biological mechanism, of the form $\lambda_b(N, l) = \tilde{c}(1 - \frac{1}{k})^{N-1}$ (see Section 6.1).

### B Inhibition Model Equilibrium

Suppose we have already selected constants $\mu_b, \beta_0, \beta_1$. We can write $\lambda_b^{inhib} = \mu_b + f(N, l)$ where $f(N, l) = \tilde{c}(1 - \frac{1}{k})^{N-1} - \mu_b$. In order for this birth rate to be valid for this selection of parameters $\mu_b, \beta_0, \beta_1$, we must be able to satisfy Equation 2 by fixing $\tilde{c}$ and $l_0$, specifically for $N_{opt} = \beta_0 + \beta_1 l$. In other words, this birth rate must maintain linear copy number control. This section will demonstrate that selecting $\tilde{c}$ and $l_0$ as functions of $\beta_0, \beta_1, \mu_b$ will give us precisely this behaviour. Specifically, we wish to show that:

$$f(N_{opt}(l), l) \equiv 0,$$

for all $l$, or equivalently,

$$\mu_b \equiv \tilde{c}\left(1 - \frac{1}{k(l)}\right)^{N_{opt}(l)-1},$$

for all $l$ (where $k(l) = l/l_0$, $N_{opt}(l) = \beta_0 + \beta_1 l$), for the right choice of parameters $\tilde{c}, l_0$. To that end, we compute:

$$\begin{aligned}
\mu_b &= \tilde{c}(1 - \frac{1}{k})^{N_{opt}-1} \implies \\
N_{opt} &= 1 + \frac{\log(\mu_b/\tilde{c})}{\log(1 - 1/k)} \\
&\approx 1 + \frac{\log(\mu_b/\tilde{c})}{-1/k - 1/2k^2} \\
&= 1 - (k - \frac{1}{2})\log\left[\frac{\mu_b}{\tilde{c}}\right] + \frac{1}{2(2k+1)}\log\left[\frac{\mu_b}{\tilde{c}}\right] \\
&\approx 1 - (k - \frac{1}{2})\log\left[\frac{\mu_b}{\tilde{c}}\right] \\
&= \left(1 - \frac{1}{2}\log\left[\frac{\tilde{c}}{\mu_b}\right]\right) + \left(\frac{1}{l_0}\log\left[\frac{\tilde{c}}{\mu_b}\right]\right) l \\
&\approx \frac{1}{l_0}\log\left[\frac{\tilde{c}}{\mu_b}\right] l.
\end{aligned} \tag{60}$$

Suppose that we allow $\tilde{c}$ to vary as a function of $\alpha$, i.e., $\tilde{c} = c(1 + \alpha/l)$, where $c$ is constant:

$$
\begin{aligned}
N_{opt} &= \frac{1}{l_0}\log\left[\frac{c^{\frac{l+\alpha}{l}}}{\mu_b}\right] l \\
&= \frac{1}{l_0}\log\left[\frac{l+\alpha}{l}\right] l + \frac{1}{l_0}\log\left[\frac{c}{\mu_b}\right] l \\
&\approx \frac{\alpha}{l_0} + \frac{1}{l_0}\log\left[\frac{c}{\mu_b}\right] l,
\end{aligned} \tag{61}
$$

giving us the intercept term we require. Picking any value $c > \mu_b$, we can now simply set:

$$
\begin{aligned}
l_0 &= \frac{\beta_1}{\log[c/\mu_b]} \\
\alpha &= \beta_0 l_0
\end{aligned} \tag{62}
$$

in order to obtain $N_{opt} = \beta_0 + \beta_1 l$. Having set these parameters, these computations can then be carried out in reverse to show that if $N_{opt} = \beta_0 + \beta_1 l$, it follows that $\mu_b = \tilde{c}(1 - \frac{1}{k(l)})^{N_{opt}(l)-1}$ for all $l$, and so $f(N_{opt}(l), l) \equiv 0$. Performing the same computation but with inequalities rather than equalities verifies that $f$ does indeed satisfy Equation 2, as required.

This demonstrates that having selected $\mu_b, \beta_0, \beta_1$ (and another parameter $c > \mu_b$), we are able to ensure $\lambda_b^{inh}$ fits Equation 2 by fixing constants $l_0, \alpha$ as functions of these parameters, hence demonstrating that the inhibition model gives rise to a valid birth rate.

### C Colocalisation Error

We can model the number of nucleoids missed due to colocalisation error using the same framework as the inhibition model. Split the mitochondrial network of volume $l$ into $k$ compartments, and suppose that if two nucleoids share the same compartment, then they are sufficiently close as to appear to be one nucleoid. Then the expected number of nucleoids we observe ($N_{obs}$) will be:

$$
N_{obs} = \mathbb{E}\left[\sum_{i=1}^{k} \mathbb{1}[X_i \geq 1]\right],
$$

where $(X_1, \ldots, X_k) \sim \text{multinomial}(1/k, N)$ as before denotes the number of nucleoids in compartment $1, \ldots, k$. Now:

$$
\begin{aligned}
N_{obs} = \mathbb{E}\left[\sum_{i=1}^{k} \mathbb{1}[X_i \geq 1]\right] &= \sum_{i=1}^{k}[1 - \mathbb{P}[X_i = 0]] \\
&= k\left(1 - \left(1 - \frac{1}{k}\right)^N\right)
\end{aligned} \tag{63}
$$

Our resolution is $\sim 100nm$ in diameter (for lattice structured illumination microscopy, i.e, the pulse and pulse-chase experiments), which corresponds to a resolution of roughly $\sim 10^6 nm^3 = 10^{-3} \mu m^3$ in volume. Then $k = \frac{l}{10^{-3}}$, and so the number of nucleoids we miss on average is:

$$
N - N_{obs} = N - 10^3 l\left(1 - \left(1 - \frac{1}{10^3 l}\right)^N\right) \approx N_{opt} - 10^3 l\left(1 - \left(1 - \frac{1}{10^3 l}\right)^{N_{opt}}\right)
$$

where $N_{opt} = \beta_0 + \beta_1 l$. But now, using our estimated values for $\beta_0, \beta_1$ (Section 3.1.2), it is easily verifiable that this is a decreasing function of $l$, and that $N - N_{err} < 1$ for $l > 20$. In our data, we have $l$ on the order of $10^3$, rendering this error negligible.

### D Stochastic Systems

Here we detail full equations for the pulse and chase portion dynamics for the two- and three-population models. We begin with the two-population pulse:

$$
\begin{aligned}
&n_o^{(0)} \xrightarrow{\lambda_b} n_r^{(0)}\\
&n_o^{(1)} \xrightarrow{\lambda_b} n_r^{(1)}\\
&n_o^{(2)} \xrightarrow{\lambda_b} n_r^{(2)}\\
\\
&n_r^{(0)} \xrightarrow{p\mu_r} 2n_o^{(1)}\\
&n_r^{(1)} \xrightarrow{p\mu_r} n_o^{(1)} + n_o^{(2)}\\
&n_r^{(2)} \xrightarrow{p\mu_r} 2n_o^{(2)}\\
\\
&n_r^{(0)} \xrightarrow{(1-p)\mu_r} n_o^{(1)} + n_r^{(1)}\\
&n_r^{(1)} \xrightarrow{(1-p)\mu_r/2} n_o^{(2)} + n_r^{(1)}\\
&n_r^{(1)} \xrightarrow{(1-p)\mu_r/2} n_o^{(1)} + n_r^{(2)}\\
&n_r^{(2)} \xrightarrow{(1-p)\mu_r} n_o^{(2)} + n_r^{(2)}\\
\\
&n_o^{(0)} \xrightarrow{\mu_d} \phi\\
&n_o^{(1)} \xrightarrow{\mu_d} \phi\\
&n_o^{(2)} \xrightarrow{\mu_d} \phi.
\end{aligned}
\tag{64}
$$

Two-population chase:

$$
\begin{aligned}
&n_o^{(2)} \xrightarrow{\lambda_b} o_r^{(2)}\\
&n_o^{(1)} \xrightarrow{\lambda_b} n_r^{(1)}\\
&n_o^{(0)} \xrightarrow{\lambda_b} n_r^{(0)}\\
\\
&n_r^{(2)} \xrightarrow{p\mu_r} 2n_o^{(2)}\\
&n_r^{(1)} \xrightarrow{p\mu_r} n_o^{(1)} + n_o^{(0)}\\
&n_r^{(0)} \xrightarrow{p\mu_r} 2n_o^{(0)}
\end{aligned}
\tag{65}
$$

$$
\begin{aligned}
&n_r^{(2)} \xrightarrow{(1-p)\mu_r} n_o^{(1)} + n_r^{(1)}\\
&n_r^{(1)} \xrightarrow{(1-p)\mu_r/2} n_o^{(0)} + n_r^{(1)}\\
&n_r^{(1)} \xrightarrow{(1-p)\mu_r/2} n_o^{(1)} + n_r^{(0)}\\
&N_r^{(0)} \xrightarrow{(1-p)\mu_r} n_o^{(0)} + n_r^{(0)}\\
\\
&n_{rem}^{(0)} \xrightarrow{p\mu_r} 2n_o^{(1)}\\
&n_{rem}^{(1)} \xrightarrow{p\mu_r} n_o^{(1)} + n_o^{(2)}\\
&n_{rem}^{(2)} \xrightarrow{p\mu_r} 2n_o^{(2)}\\
\\
&n_{rem}^{(0)} \xrightarrow{(1-p)\mu_r} n_o^{(1)} + n_r^{(1)}\\
&n_{rem}^{(1)} \xrightarrow{(1-p)\mu_r/2} n_o^{(2)} + n_r^{(1)}\\
&n_{rem}^{(1)} \xrightarrow{(1-p)\mu_r/2} n_o^{(1)} + n_r^{(2)}\\
&n_{rem}^{(2)} \xrightarrow{(1-p)\mu_r} n_o^{(2)} + n_r^{(2)}\\
\\
&n_o^{(0)} \xrightarrow{\mu_d} \phi\\
&n_o^{(1)} \xrightarrow{\mu_d} \phi\\
&n_o^{(2)} \xrightarrow{\mu_d} \phi.
\end{aligned}
$$

Three-population pulse:

$$
\begin{aligned}
&n_y^{(0)} \xrightarrow{\lambda_b} n_r^{(0)}\\
&n_y^{(1)} \xrightarrow{\lambda_b} n_r^{(1)}\\
&n_y^{(2)} \xrightarrow{\lambda_b} n_r^{(2)}\\
\\
&n_r^{(0)} \xrightarrow{p\mu_r} 2n_y^{(1)}\\
&n_r^{(1)} \xrightarrow{p\mu_r} n_y^{(1)} + n_o^{(2)}\\
&n_r^{(2)} \xrightarrow{p\mu_r} 2n_y^{(2)}
\end{aligned}
\tag{66}
$$

$$
\begin{aligned}
&n_r^{(0)} \xrightarrow{(1-p)\mu_r} n_y^{(1)} + n_r^{(1)} \\
&n_r^{(1)} \xrightarrow{(1-p)\mu_r/2} n_y^{(2)} + n_r^{(1)} \\
&n_r^{(1)} \xrightarrow{(1-p)\mu_r/2} n_y^{(1)} + n_r^{(2)} \\
&n_r^{(2)} \xrightarrow{(1-p)\mu_r} n_y^{(2)} + n_r^{(2)} \\
\\
&n_y^{(0)} \xrightarrow{\mu_a} n_o^{(0)} \\
&n_y^{(1)} \xrightarrow{\mu_a} n_o^{(1)} \\
&n_y^{(2)} \xrightarrow{\mu_a} n_o^{(2)} \\
\\
&n_o^{(0)} \xrightarrow{\mu_d} \phi \\
&n_o^{(1)} \xrightarrow{\mu_d} \phi \\
&n_o^{(2)} \xrightarrow{\mu_d} \phi
\end{aligned}
$$

Three-population chase:

$$
\begin{aligned}
&n_y^{(2)} \xrightarrow{\lambda_b} o_r^{(2)} \\
&n_y^{(1)} \xrightarrow{\lambda_b} n_r^{(1)} \\
&n_y^{(0)} \xrightarrow{\lambda_b} n_r^{(0)} \\
\\
&n_r^{(2)} \xrightarrow{p\mu_r} 2n_y^{(2)} \\
&n_r^{(1)} \xrightarrow{p\mu_r} n_y^{(1)} + n_y^{(0)} \\
&n_r^{(0)} \xrightarrow{p\mu_r} 2n_y^{(0)}
\end{aligned}
\tag{67}
$$

$$n_r^{(2)} \xrightarrow{(1-p)\mu_r} n_y^{(1)} + n_r^{(1)}$$
$$n_r^{(1)} \xrightarrow{(1-p)\mu_r/2} n_y^{(0)} + n_r^{(1)}$$
$$n_r^{(1)} \xrightarrow{(1-p)\mu_r/2} n_y^{(1)} + n_r^{(0)}$$
$$N_r^{(0)} \xrightarrow{(1-p)\mu_r} n_y^{(0)} + n_r^{(0)}$$

$$n_{rem}^{(0)} \xrightarrow{p\mu_r} 2n_y^{(1)}$$
$$n_{rem}^{(1)} \xrightarrow{p\mu_r} n_y^{(1)} + n_y^{(2)}$$
$$n_{rem}^{(2)} \xrightarrow{p\mu_r} 2n_y^{(2)}$$

$$n_{rem}^{(0)} \xrightarrow{(1-p)\mu_r} n_y^{(1)} + n_r^{(1)}$$
$$n_{rem}^{(1)} \xrightarrow{(1-p)\mu_r/2} n_y^{(2)} + n_r^{(1)}$$
$$n_{rem}^{(1)} \xrightarrow{(1-p)\mu_r/2} n_y^{(1)} + n_r^{(2)}$$
$$n_{rem}^{(2)} \xrightarrow{(1-p)\mu_r} n_y^{(2)} + n_r^{(2)}$$

$$n_y^{(0)} \xrightarrow{\mu_a} n_o^{(0)}$$
$$n_y^{(1)} \xrightarrow{\mu_a} n_o^{(0)}$$
$$n_y^{(2)} \xrightarrow{\mu_a} n_o^{(0)}$$

$$n_o^{(0)} \xrightarrow{\mu_d} \phi$$
$$n_o^{(1)} \xrightarrow{\mu_d} \phi$$
$$n_o^{(2)} \xrightarrow{\mu_d} \phi$$

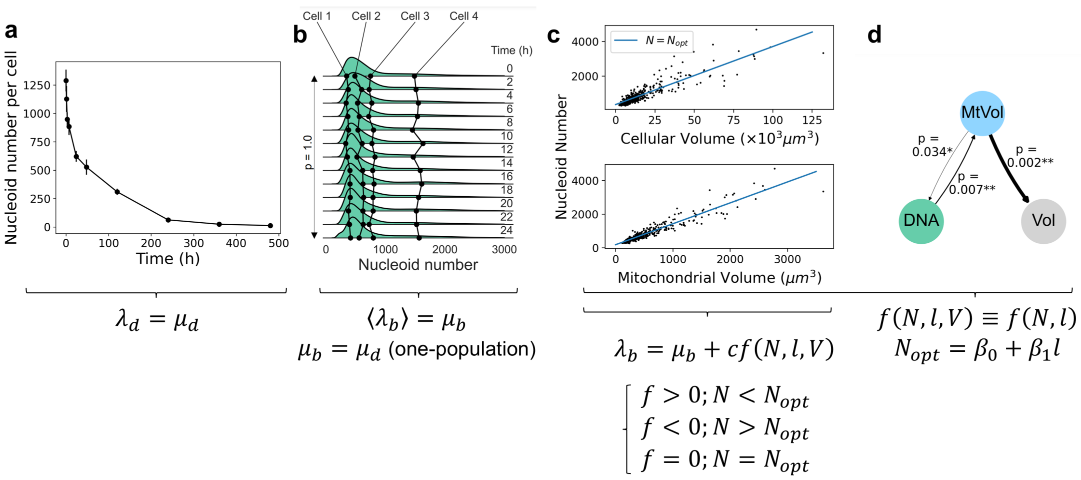

**Supplementary Fig. 1: Key empirical observations that inform replication and degradation rates.** **a**, Under exposure to ethidium bromide, nucleoid number exhibits an exponential-like decay, necessitating a constant per-capita degradation rate. **b**, Distribution of nucleoid number from 4D lattice sheet imaging is constant over time, necessitating that the replication rate is on average equal to the degradation rate. **c**, The optimal nucleoid number for a cell scales linearly with cellular and mitochondrial volume, necessitating an adaptive birth rate. **d**, Causal network from linear Granger causality, and other causal techniques, necessitate that the replication rate is not a function of cellular volume.

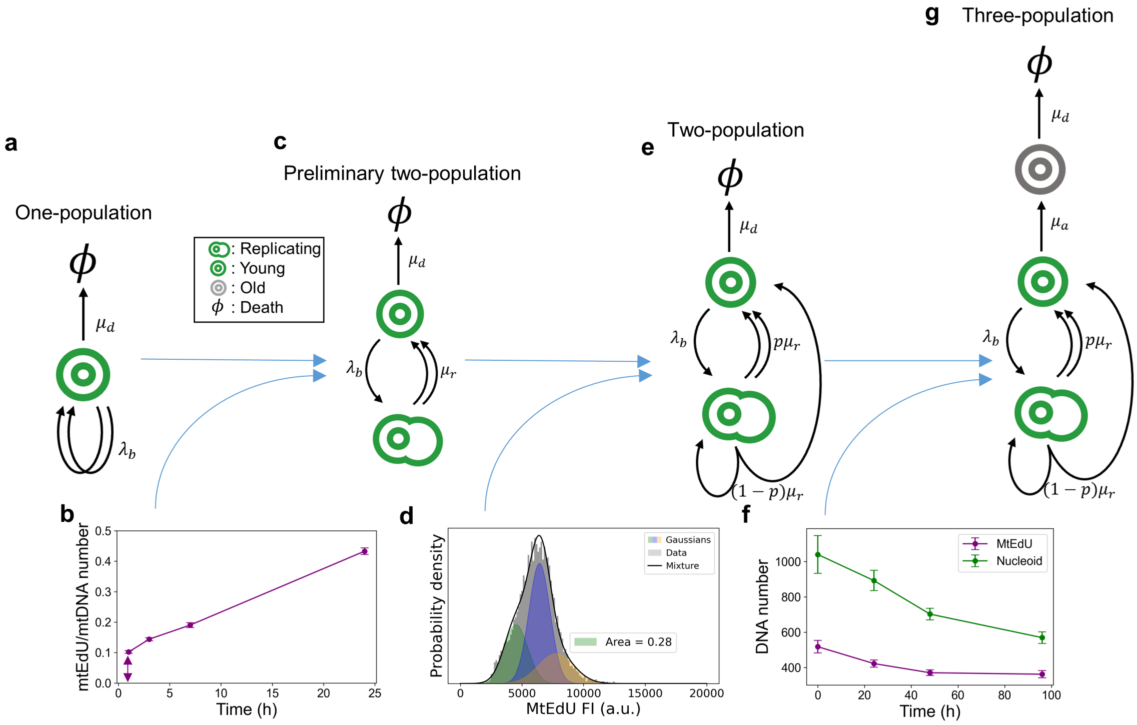

**Supplementary Fig. 2: Key empirical observations that drive model design.** **a**, Schematic of the one-population model. **b**, MtEdU number does not start from zero, necessitating the addition of a replicating subpopulation that becomes tagged immediately. **c**, Schematic of the preliminary two-population model. **d**, Gaussian mixture modelling demonstrates that the majority of mtEdU is doubly tagged following the pulse, necessitating the addition of preferential replication of newly replicated molecules. **e**, Schematic of the two-population model. **f**, MtEdU does not drop as rapidly as mtDNA during the chase, necessitating the addition of preferential degradation of older molecules. **g**, Schematic of the three-population model. Single arrows indicate a movement event, while double arrows indicate a doubling and movement event, with each arrow representing the movement of each daughter mtDNA. Labels are Poisson rates.

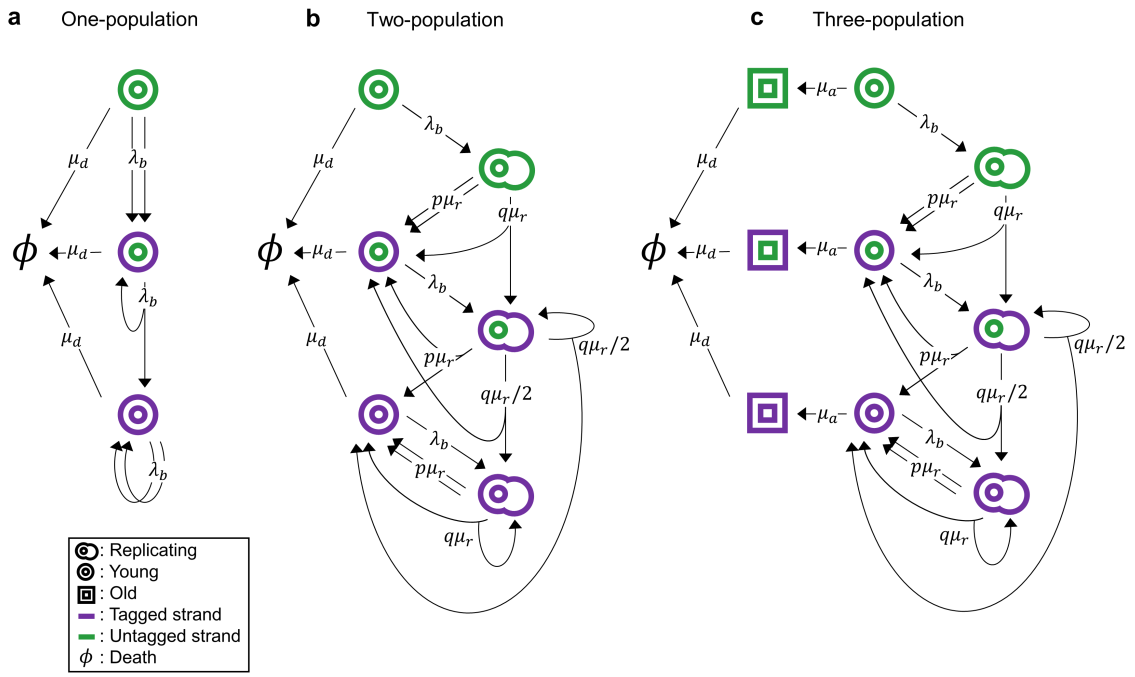

**Supplementary Fig. 3: Expanded model diagrams with EdU incorporation**. **a,** One-population model. **b**, Two-population model. **c**, Three-population model. Single arrows indicate a movement event, while double arrows indicate a doubling and movement event, with each arrow representing the movement of each daughter mtDNA. Labels are Poisson rates. Despite the apparent complexity, diagrams represent models with no more than 5 distinct Poisson rates.

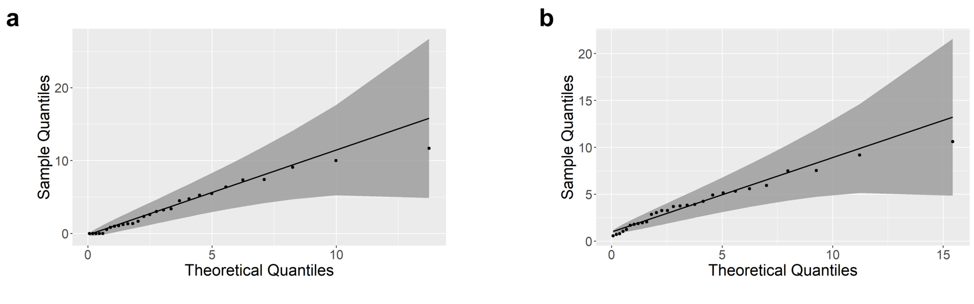

**Supplementary Fig. 4: Error quantification**. Exponential Q-Q plots with 95% confidence band of percentage of: **a**, tagged molecules undercounted, **b**, untagged molecules undercounted.

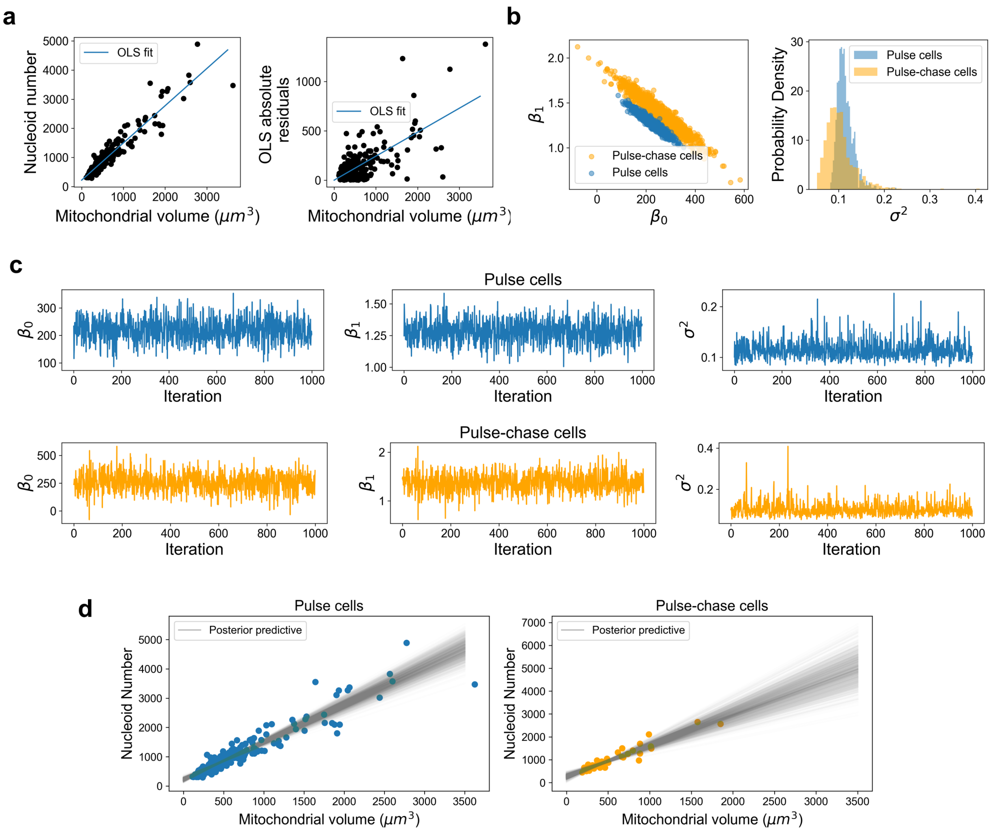

**Supplementary Fig. 5: Heteroscedastic Bayesian linear regression $\beta_0$, $\beta_1$.** **a**, OLS regression of nucleoid number against mitochondrial voume (left) and its absolute residuals (right), demonstrating linear heteroscedasticity. **b**, $\beta_0, \beta_1$ joint posterior (left), $\sigma^2$ posterior (right). **c**, Gibbs sampling trace plots for the pulse cells (top) and the pulse-chase cells (bottom), showing good mixing. **d**, Posterior predictive distribution of the linear regression line, for the pulse (left) and pulse-chase (right) cells.

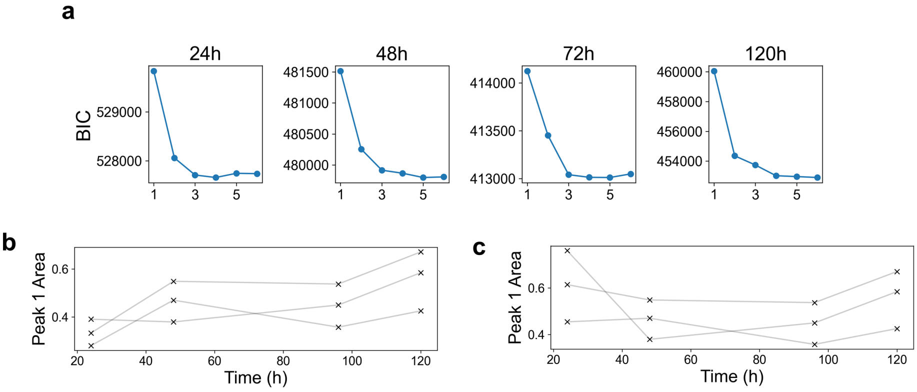

**Supplementary Fig. 6: Gaussian mixture modelling peak number selection, for chase portion**. **a**, Averaged BIC across all assays of the Gaussian mixture models with different peak number, showing sharp drop at 2 peaks, and less consequential drops thereafter. **b**, Inferred singly tagged proportions for the three assays over time, where the 24 hour time point has been fit with 3 peaks, displaying the parsimonious biological interpretation that the proportion of singly tagged molecules steadily increases over the chase. **c**, As in b, but with the 24 hour time point fit with 2 peaks, showing an unreasonable biological interpretation that the singly tagged proportion is largest at the beginning of the chase.

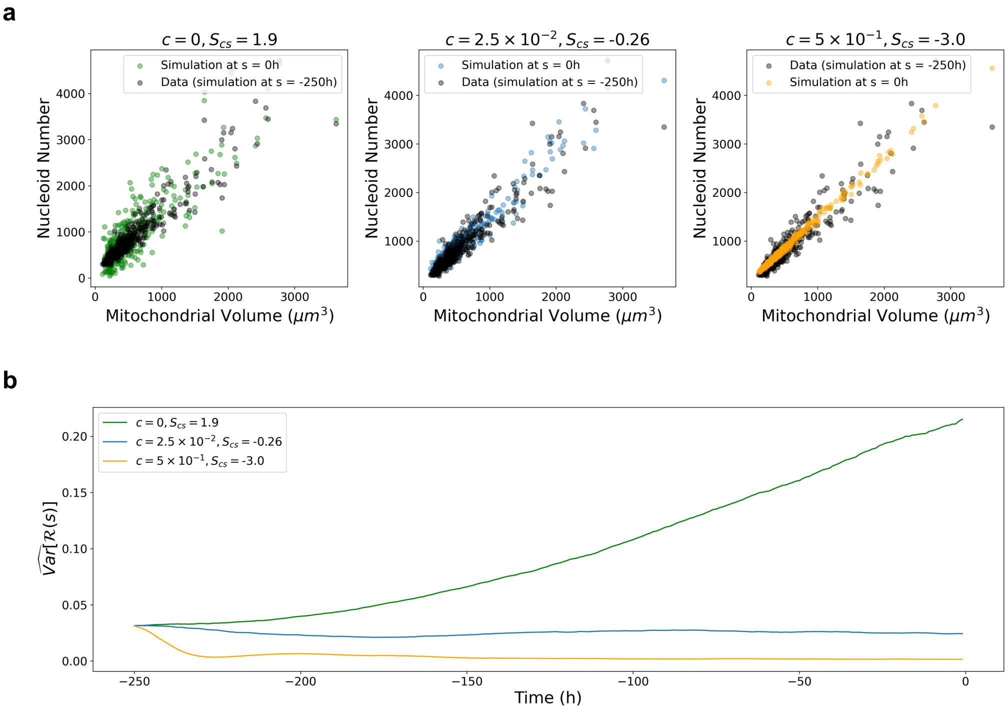

**Supplementary Fig. 7: The control strength parameter $c$ and summary statistic $S_{cs}$. a**, Three-population simulations for 3 choices of $c$, displaying a positive $\hat{S}_{cs}$ when $c$ is too small and control strength is too weak (left), a roughly 0 $\hat{S}_{cs}$ when $c$ is appropriately specified (middle), and a negative $\hat{S}_{cs}$ when $c$ is too large and the control strength is too strong (right). **b**, The variance of the log-residuals $\widehat{\mathrm{Var}}[\mathcal{R}(s)]$ over time during the burn-in period for the 3 choices of $c$ as in a, with the resulting control strength summary statistic.

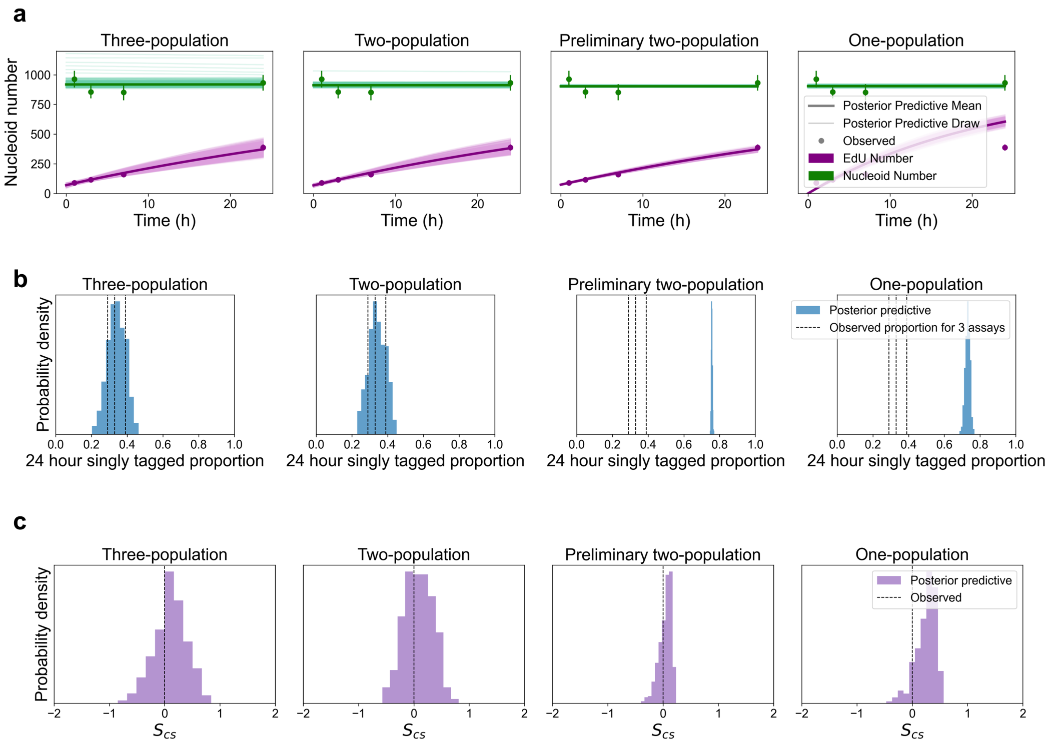

**Supplementary Fig. 8: Pulse data posterior predictive plots.** **a**, Posterior predictive trajectories of nucleoid and mtEdU number for each model. Thin lines indicate one draw from the posterior predictive distribution. Thick lines are posterior predictive means. **b**, Posterior predictive of 24h singly tagged proportion for each model. **c**, Posterior predictive of control strength summary statistic $S_{cs}$ for each model.

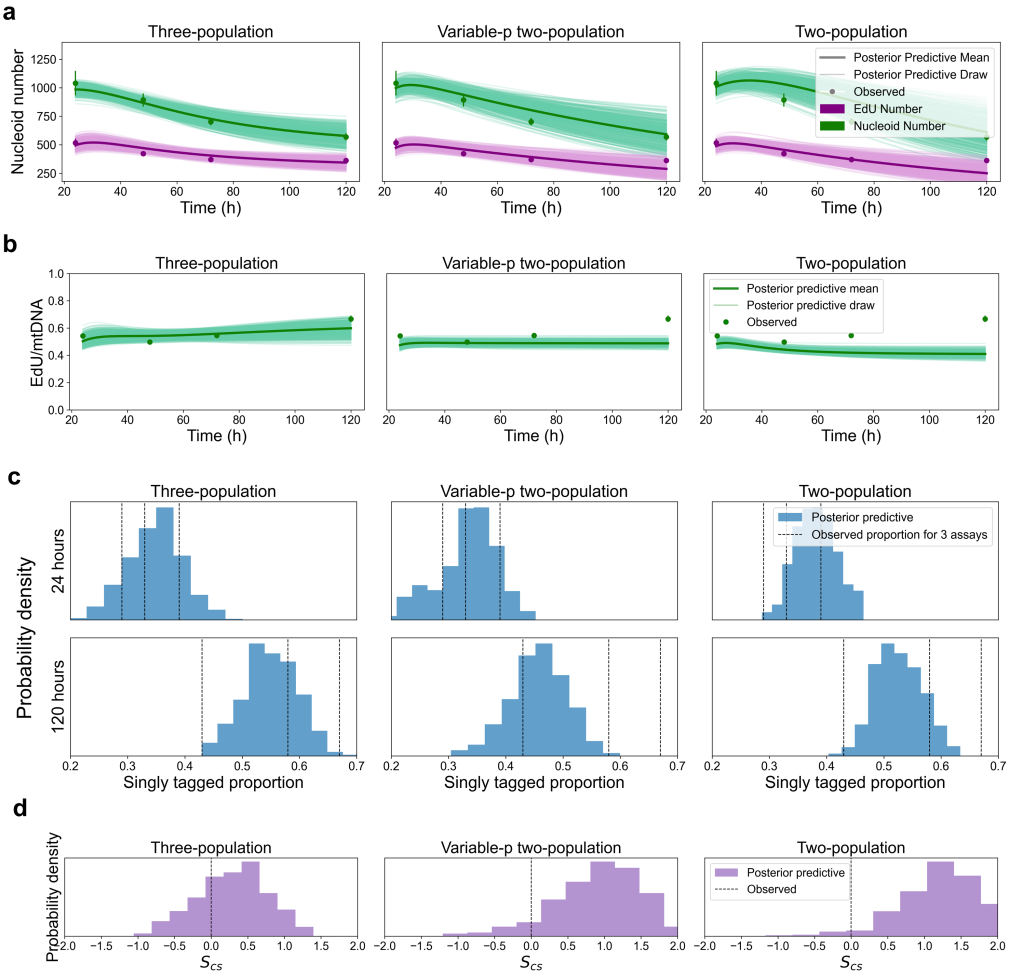

**Supplementary Fig. 9: Pulse-chase bulk posterior predictive plots. a**, Posterior predictive trajectories of nucleoid and mtEdU number for each model. **b**, Posterior predictive trajectories of mtEdU/mtDNA proportion for each model. For sub-figures a and b, thin lines indicate one draw from the posterior predictive distribution, and thick lines are posterior predictive means. **c**, Posterior predictive of 24h and 120h singly tagged proportion for each model. **d**, Posterior predictive of control strength summary statistic $S_{cs}$ for each model.

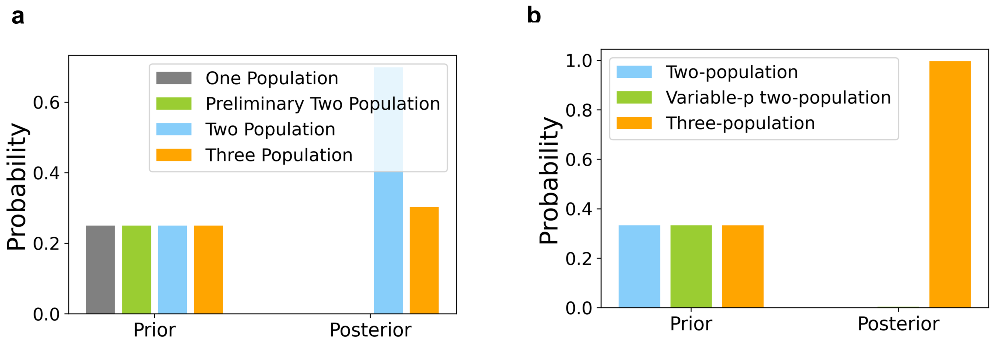

**Supplementary Fig. 10: ABC model selection** for pulse data fitting (**a**) and pulse-chase data fitting (**b**). Uniform priors were placed on each relevant model, and the ABC rejection algorithm was run, selecting the two- and three-population models for the pulse data fitting, and the three-population model for the pulse-chase data fitting.

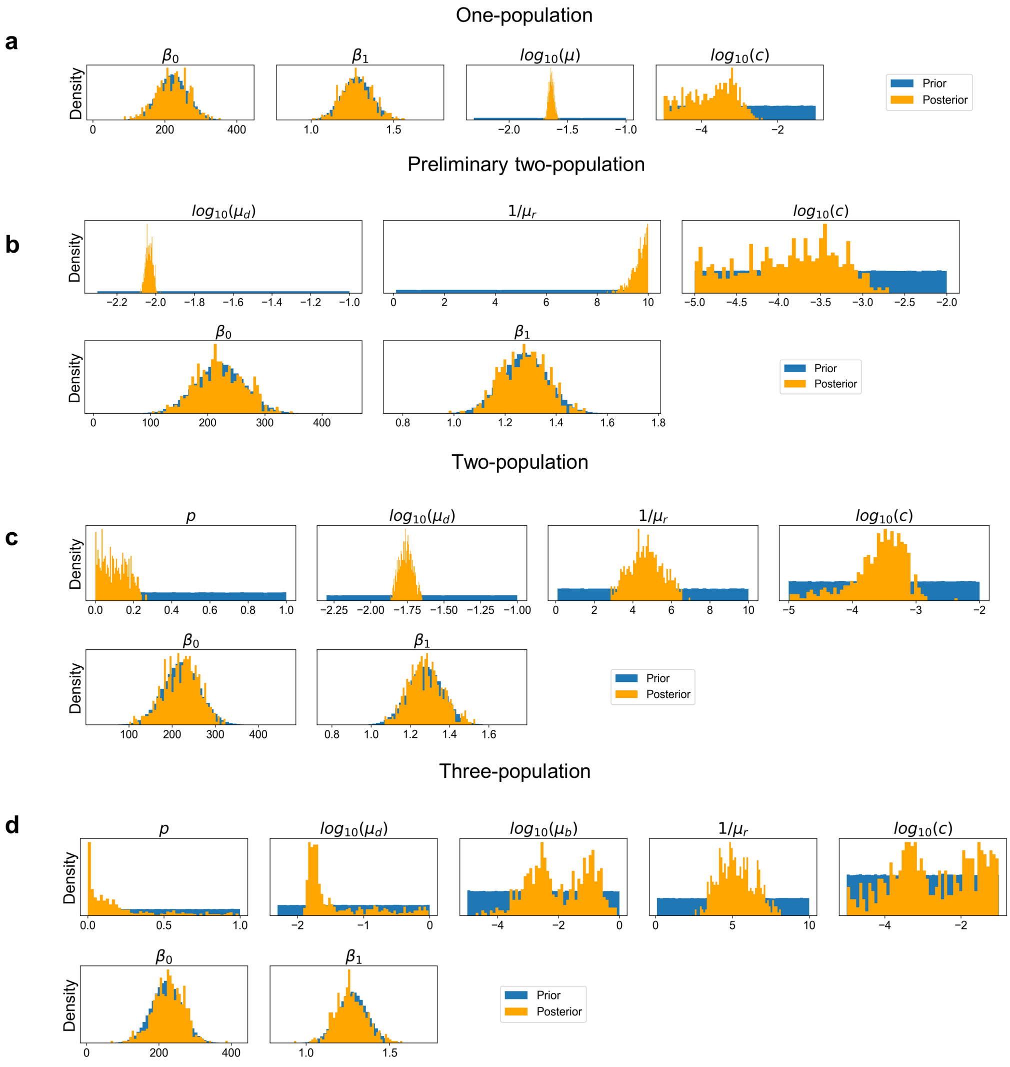

**Supplementary Fig. 11: One-dimensional marginal priors and posteriors for the pulse data fitting** for the one-population model (**a**), preliminary two-population model (**b**), two-population model (**c**), and three-population model (**d**). Priors for $\beta_0, \beta_1$ are Bayesian linear regression inferences, so posteriors appropriately match the priors for all models as all information has already been extracted for these parameters. The one- and preliminary two-population models have parameters $c$; $c, \mu_r$, respectively, extending to the edge of the prior, indicating model misspecification. The two-population model has all posteriors within the bounds of the prior (note that $p > 0$ by definition), indicating appropriate model specification. The three-population model displays degeneracy in its parameter space due to it being under-constrained by not yet being fit to the pulse-chase data, whereby it substantially reduces (Supplementary Fig. 13). Here, $\mu_b$ is the constant portion of the per-capita birth rate $\lambda_b = \mu_b + f$, with $f$ being the logarithmic control feedback term.

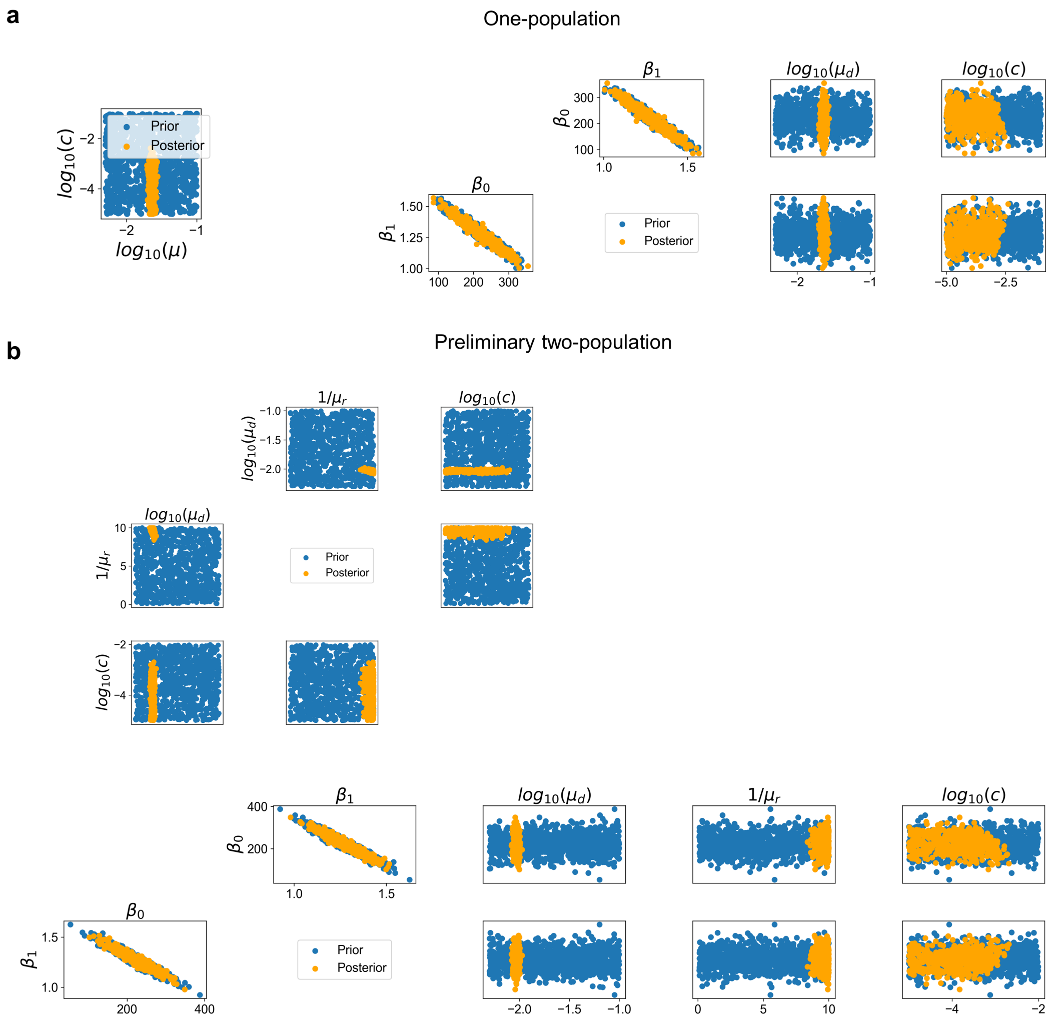
a
One-population
Prior
Posterior
$log_{10}(c)$
$log_{10}(\mu)$
$\beta_1$
$\beta_0$
$log_{10}(\mu_d)$
b
Preliminary two-population
$1/\mu_r$
$log_{10}(\mu_d)$
$log_{10}(c)$
$\beta_0$
$\beta_1$

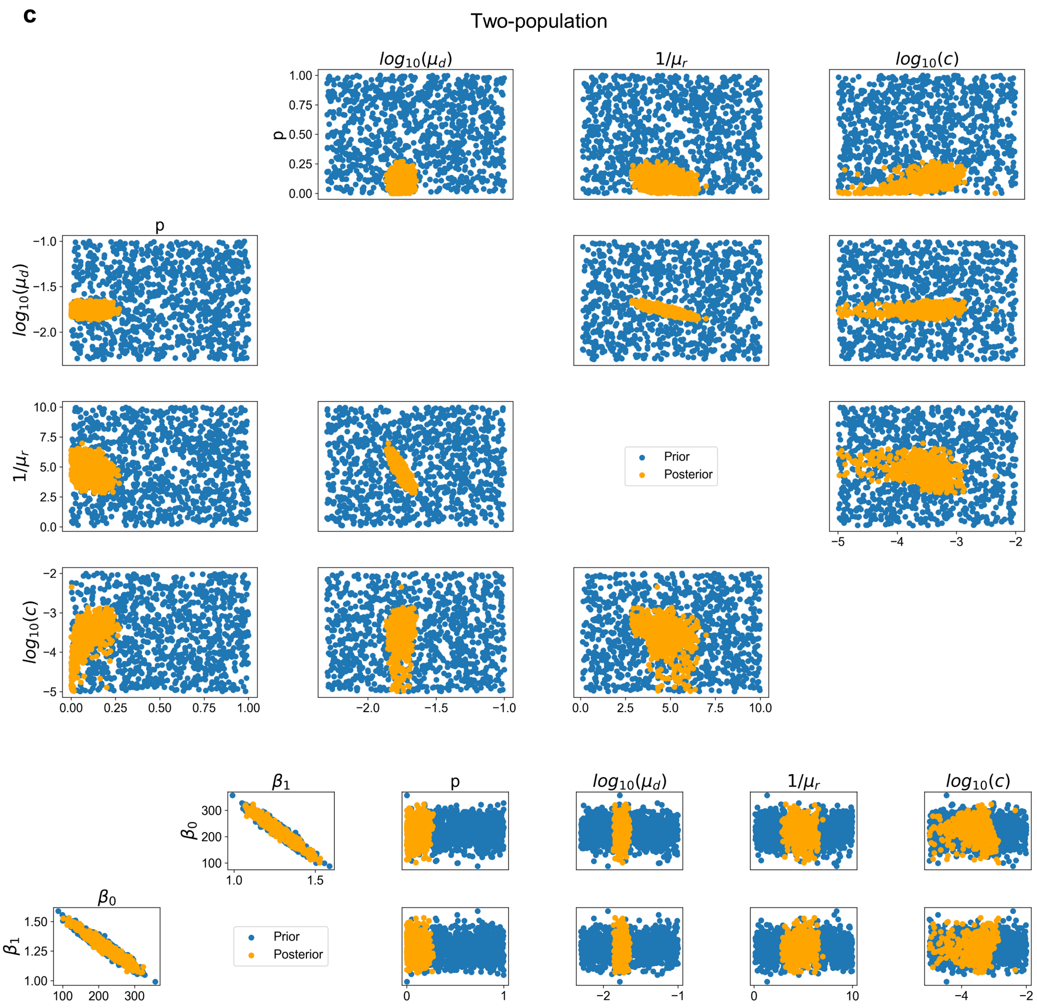
c
Two-population
$log_{10}(\mu_d)$
$1/\mu_r$
$log_{10}(c)$
p
$log_{10}(\mu_d)$
$1/\mu_r$
$log_{10}(c)$
Prior
Posterior
$\beta_1$
$\beta_0$
$\beta_0$
$\beta_1$
Prior
Posterior
p
$log_{10}(\mu_d)$
$1/\mu_r$
$log_{10}(c)$

**d**

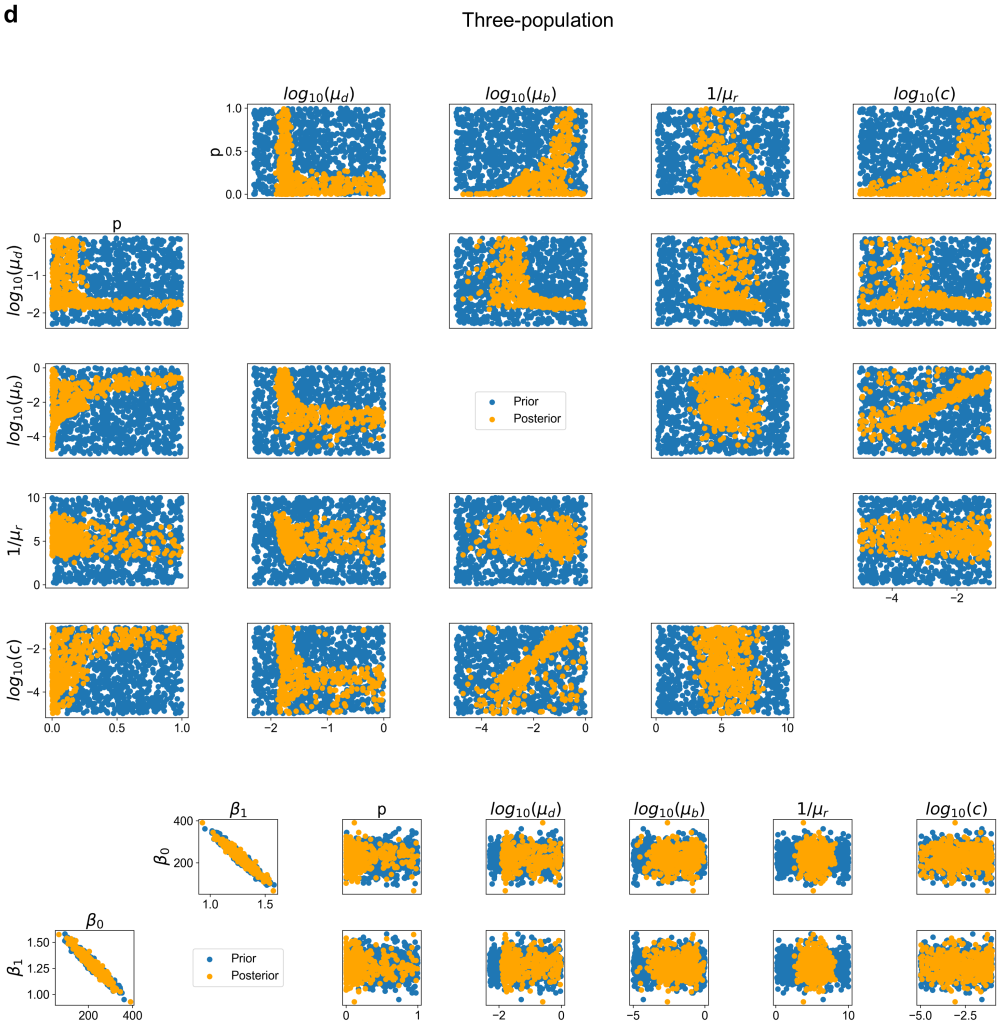

**Supplementary Fig. 12: Two-dimensional marginal priors and posteriors for the pulse data fitting** for the one-population model (**a**), preliminary two-population model (**b**), two-population model (**c**), and three-population model (**d**). Priors for $\beta_0, \beta_1$ are Bayesian linear regression inferences, so posteriors appropriately match the priors for all models as all information has already been extracted for these parameters. The one- and preliminary two-population models have parameters $c$; $c, \mu_r$, respectively, extending to the edge of the prior, indicating model misspecification. The two-population model has all posteriors within the bounds of the prior (note that $p > 0$ by definition), indicating appropriate model specification. The three-population model displays degeneracy in its parameter space due to it being under-constrained by not yet being fit to the pulse-chase data, whereby it substantially reduces (Supplementary Fig. 14).

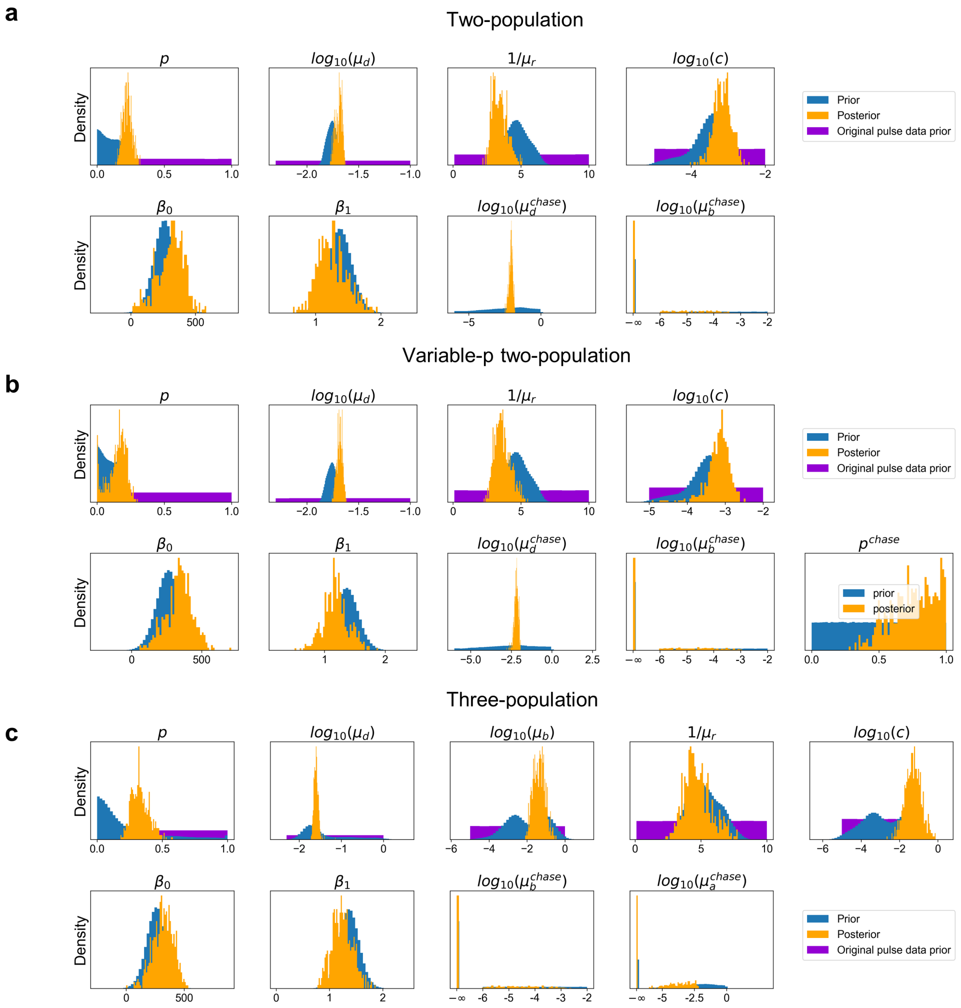

**Supplementary Fig. 13: One-dimensional marginal priors and posteriors for the pulse-chase data fitting** for the two-population model (**a**), variable-p two-population model (**b**), and three-population model (**c**). Priors for $\beta_0, \beta_1$ are Bayesian linear regression inferences, so posteriors appropriately match the priors for all models as all information has already been extracted for these parameters, with only a slight shift due to the pulse-chase cells being out-of-equilibrium. For the two-population model, pulse portion parameters $(p, \mu_d, \mu_r, c)$ are pinned to the edge of their prior, indicating model misspecification. We see similar issues in the variable-p two-population model. In the three-population model, posteriors lie within the bounds of the priors, indicating appropriate model specification. Moreover, parameter degeneracy is substantially reduced when the information from the pulse-chase cells is included (see $p, \mu_d, \mu_b, c$). We see that $\mu_b^{chase} = \mu_a^{chase} = 0$ is a favoured mode for the three-population model, with even the nonzero posterior mass being concentrated on very small values. This indicates that the data is compatible with the three-population model where no more replication or ageing is initiated, and the replicating population is allowed to finish replicating its constituents, and old populations slowly depletes (see also Supplementary Fig. 14).

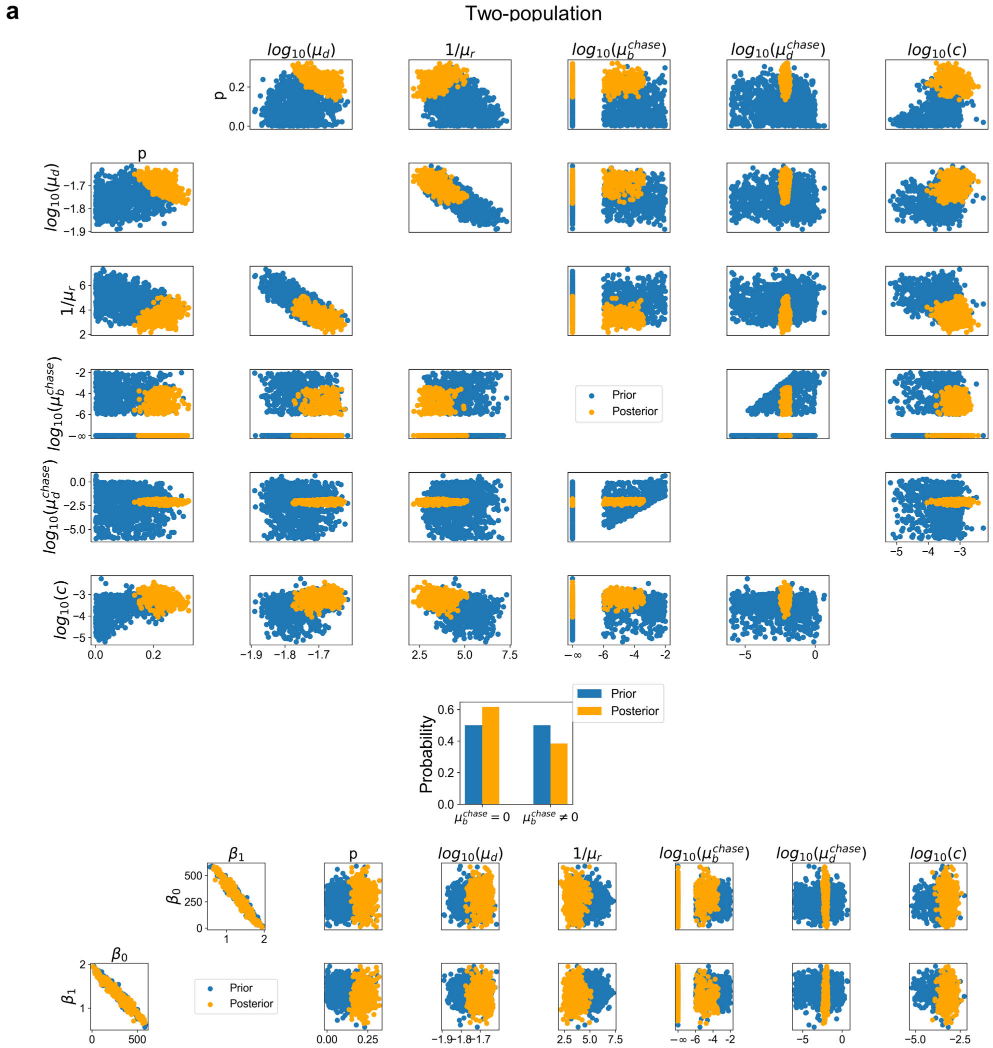

a
Two-population
Prior
Posterior
Probability

**b**

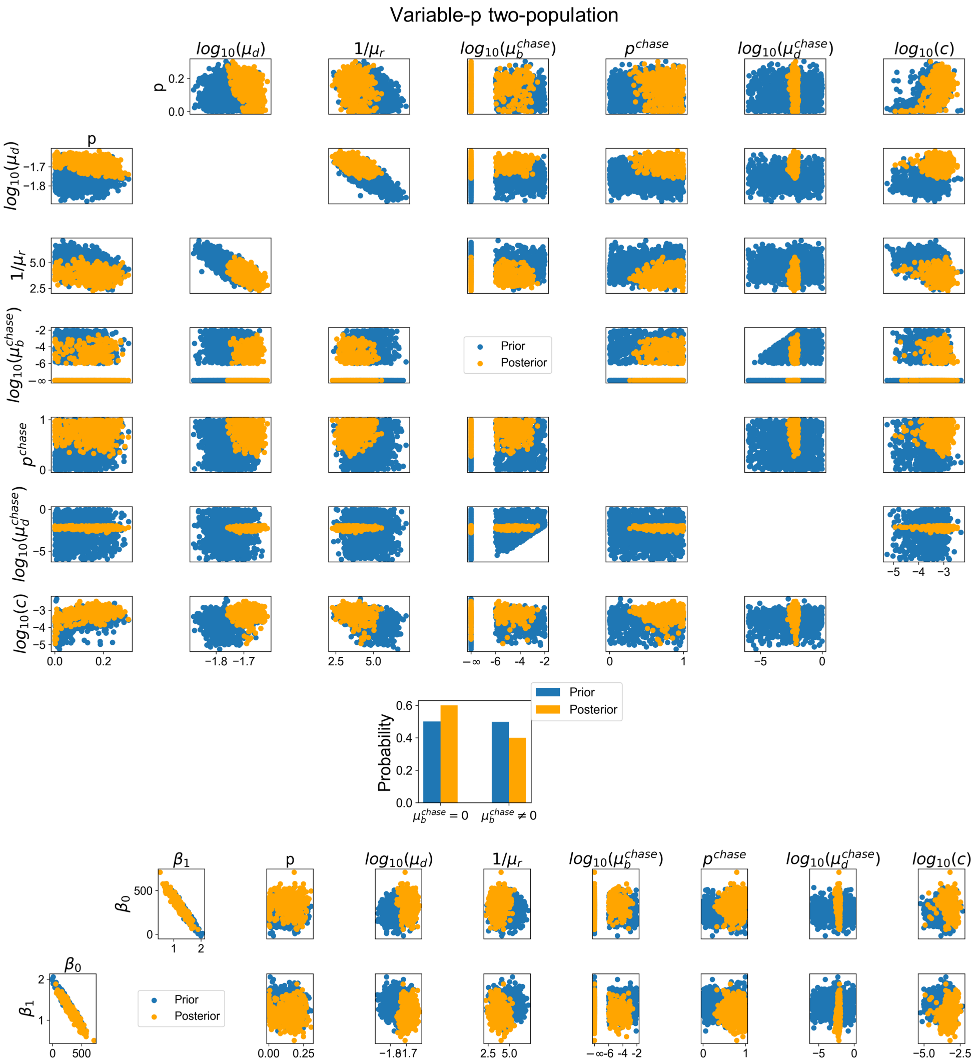
Variable-p two-population
$log_{10}(\mu_d)$
$1/\mu_r$
$log_{10}(\mu_b^{chase})$
$p^{chase}$
$log_{10}(\mu_d^{chase})$
$log_{10}(c)$
p
Prior
Posterior
Probability
$\mu_b^{chase} = 0$
$\mu_b^{chase} \neq 0$
$\beta_1$
$\beta_0$

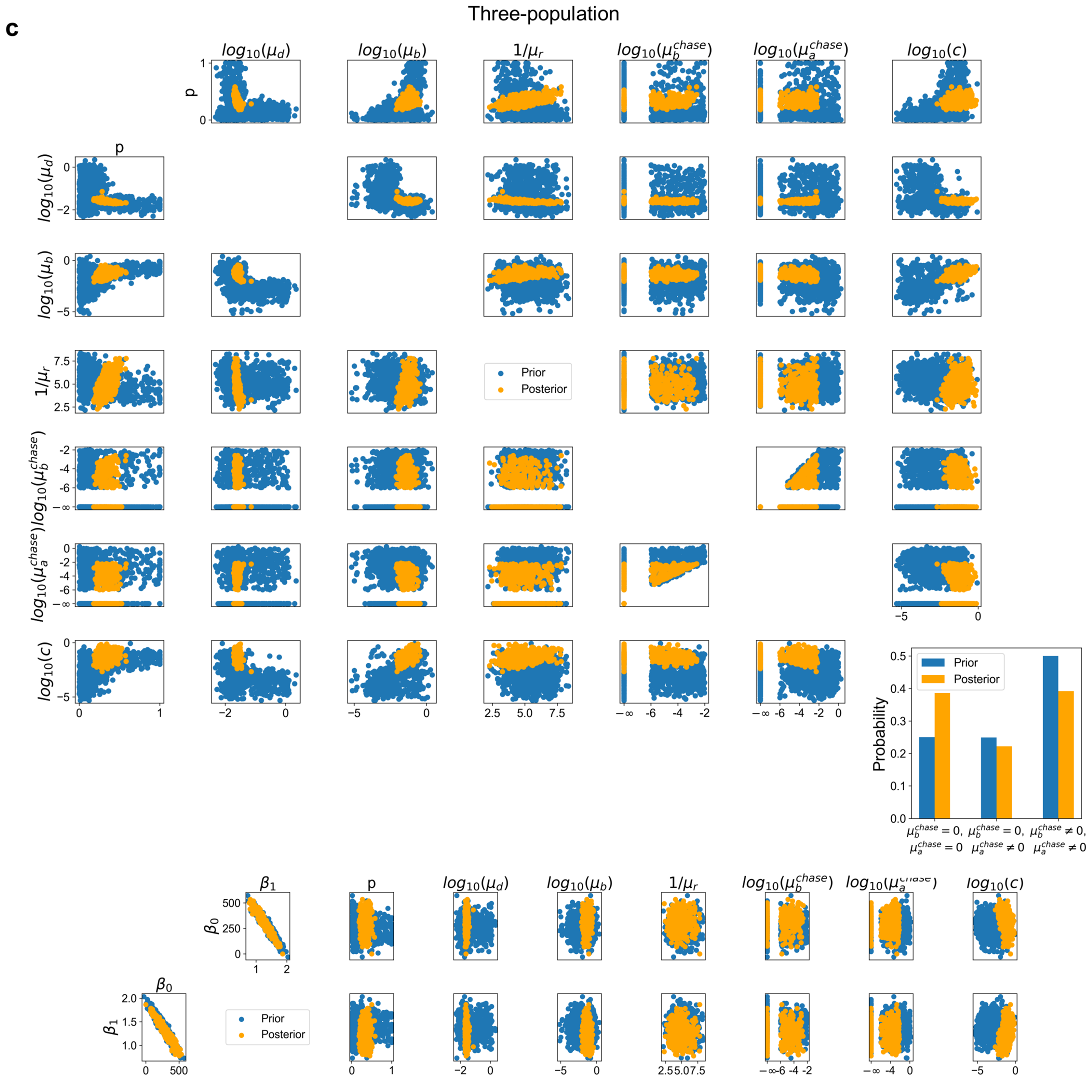

**Supplementary Fig. 14: Two-dimensional marginal priors and posteriors for the pulse-chase data fitting** for the two-population model (**a**), variable-p two-population model (**b**), and three-population model (**c**). Priors for $\beta_0, \beta_1$ are Bayesian linear regression inferences, so posteriors appropriately match the priors for all models as all information has already been extracted for these parameters, with only a slight shift due to the pulse-chase cells being out-of-equilibrium. For the two-population model, pulse portion parameters $(p, \mu_d, \mu_r, c)$ are pinned to the edge of their prior, indicating model misspecification. We see similar issues in the variable-p two-population model. In the three-population model, posteriors lie within the bounds of the priors, indicating appropriate model specification. Moreover, parameter degeneracy is substantially reduced when the information from the pulse-chase cells is included (see $p, \mu_d, \mu_b, c$). We see that $\mu_b^{chase} = \mu_a^{chase} = 0$ is a favoured mode for the three-population model, with even the nonzero posterior mass being concentrated on very small values (see Supplementary Fig. 13). This indicates that the data is compatible with the three-population model where no more replication or ageing is initiated, and the replicating population is allowed to finish replicating its constituents, and old populations slowly depletes.

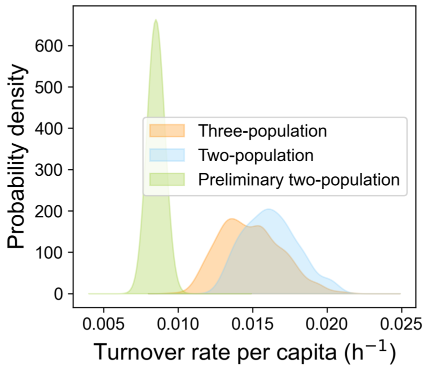

**Supplementary Fig. 15: Turnover rate for different models**. For the three- and two-population models, EdU tagged molecules are more likely to replicate again, meaning that a replication event produces one more mtEdU. For the preliminary two-population model, untagged molecules are more likely to replicate (as there are more of them), meaning that a replication event produced two more mtEdU. This difference leads to a turnover rate estimate which is double for the former two models than for the latter.

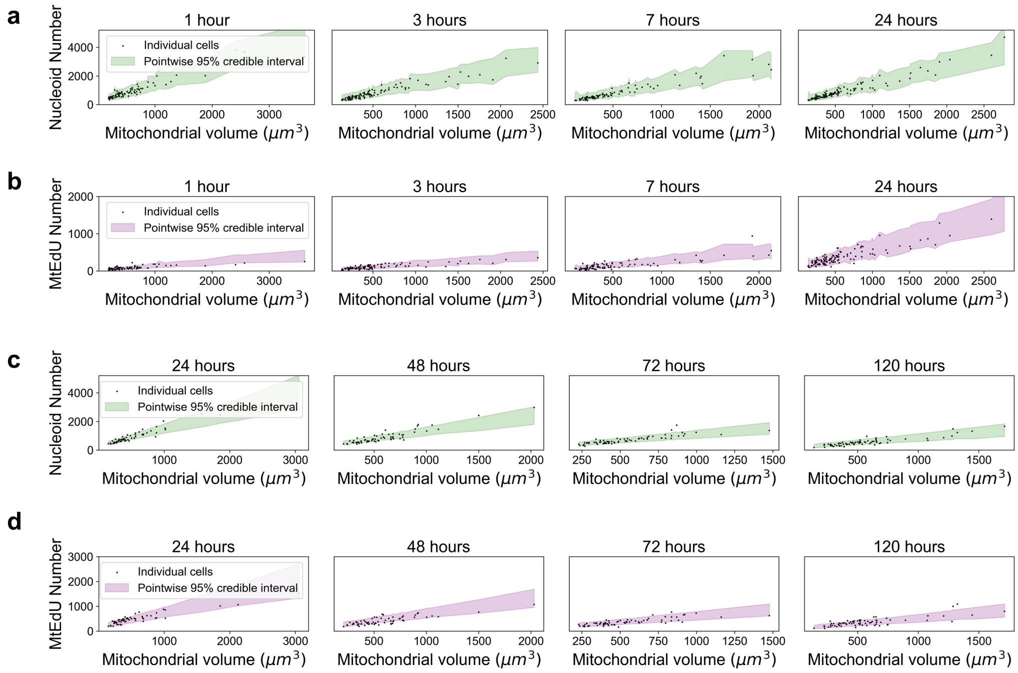

**Supplementary Fig. 16: Single cell posterior predictive plots for three-population model**. **a**, Pulse data nucleoid number. **b**, Pulse data mtEdU number. **c**, Pulse-chase data nucleoid number. **d**, Pulse-chase data mtEdU number. We see slight overdispersion in sub-figure b, with 78% of data points lying within the 95% credible intervals.

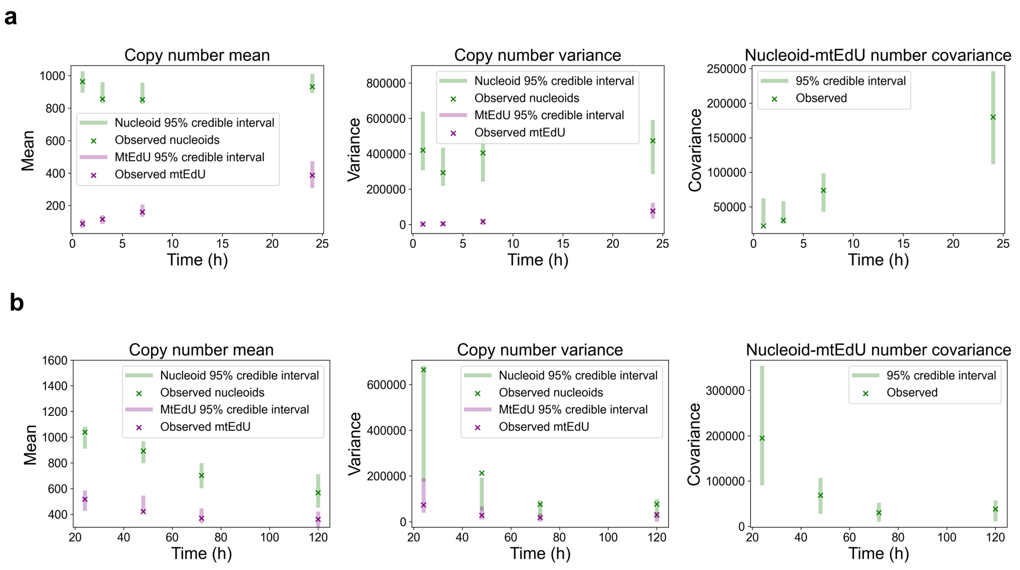

**Supplementary Fig. 17: Higher order moments**. **a**, Observed copy number mean, variance, and covariance, for the pulse data, with three-population model 95% credible intervals. **b**, Observed copy number mean, variance, and covariance, for the pulse-chase data, with three-population model 95% credible intervals.

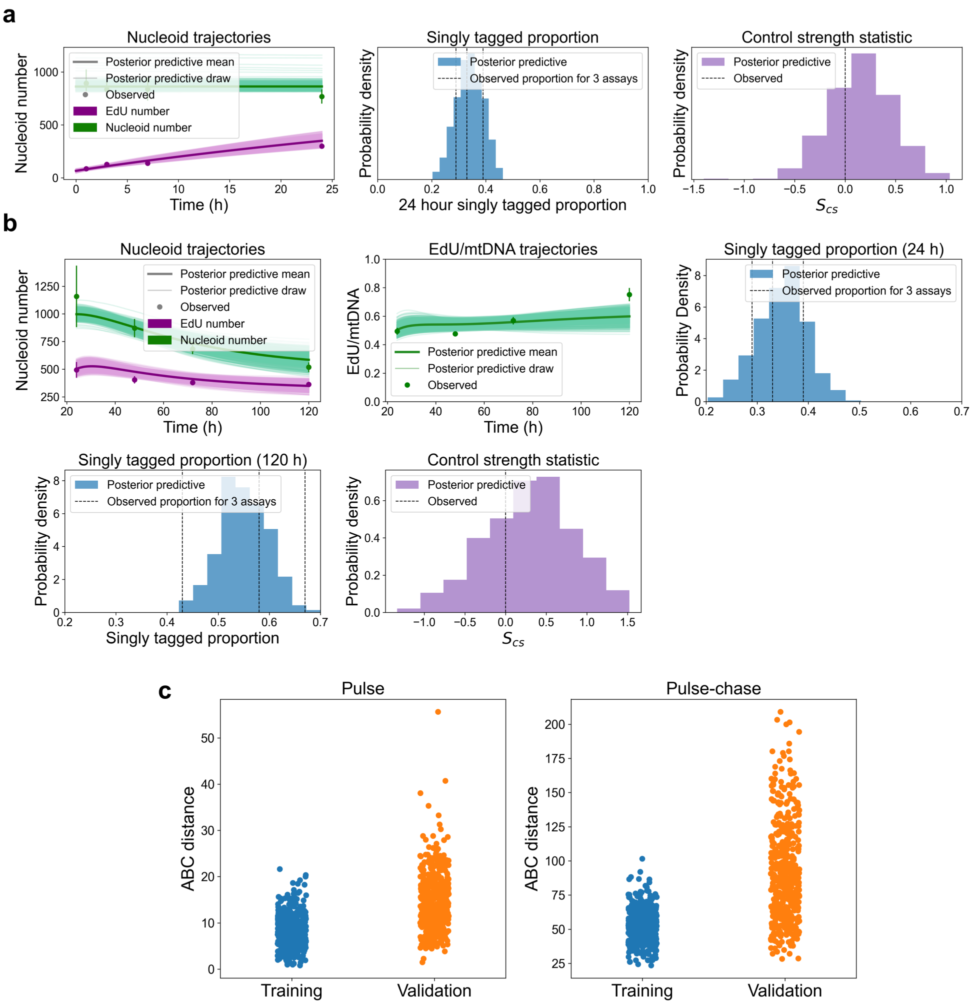

**Supplementary Fig. 18: Assay 3 validation for the three-population model**. **a**, Pulse cells posterior predictive plots using only assay 3. **b**, Pulse-chase cells posterior predictive plots using only assay 3. **c**, Training and validation ABC distances for the pulse and pulse-chase cells.

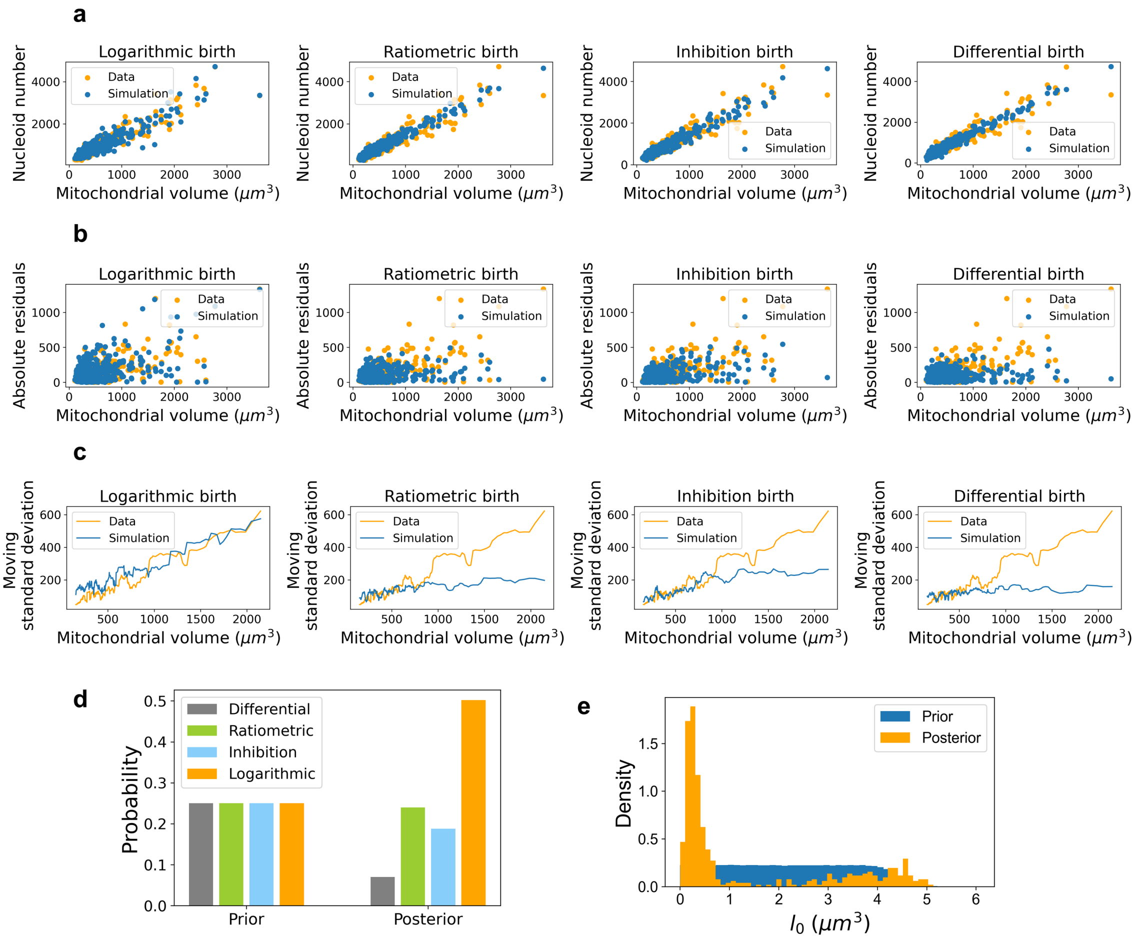

**Supplementary Fig. 19: Birth rate mechanisms are distinguished through heteroscedasticity.** **a**, Simulated cells from the three-population model for different birth rates compared to observed cells. **b**, Absolute residuals of a. **c**, Moving standard deviation of a. **d**, ABC model selection between the four birth rates. **e**, ABC posterior on $l_0$, the compartment size, for the inhibition model.

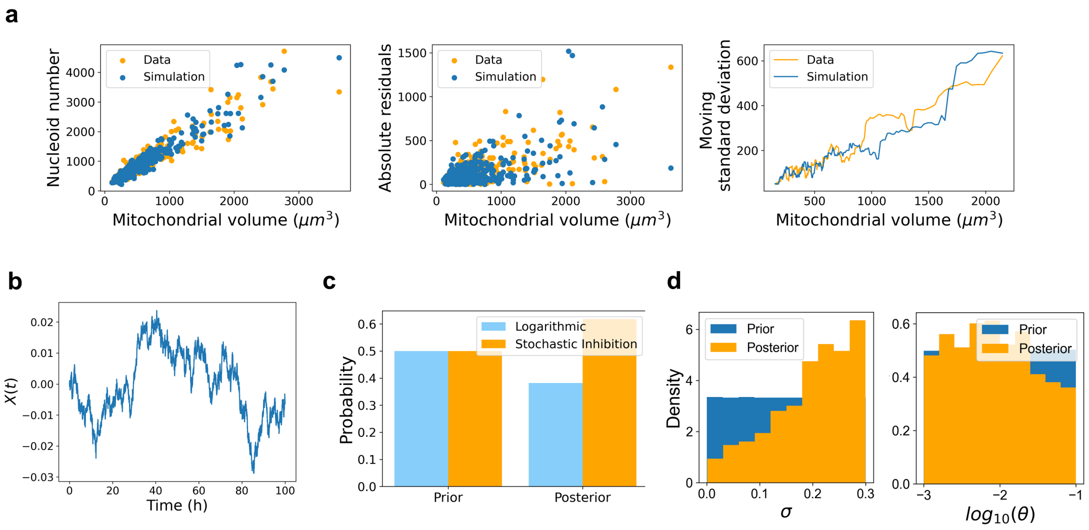

**Supplementary Fig. 20: Stochastic inhibition control.** **a**, Stochastic inhibition simulations display matching heteroscedasticity to data. **b**, An example OU process. **c**, ABC model selection between logarithmic and stochastic inhibition control. **d**, ABC posteriors for $\sigma$ and $\theta$ for stochastic inhibition control. Discussion of the shapes of the posteriors can be found in Section 6.4.

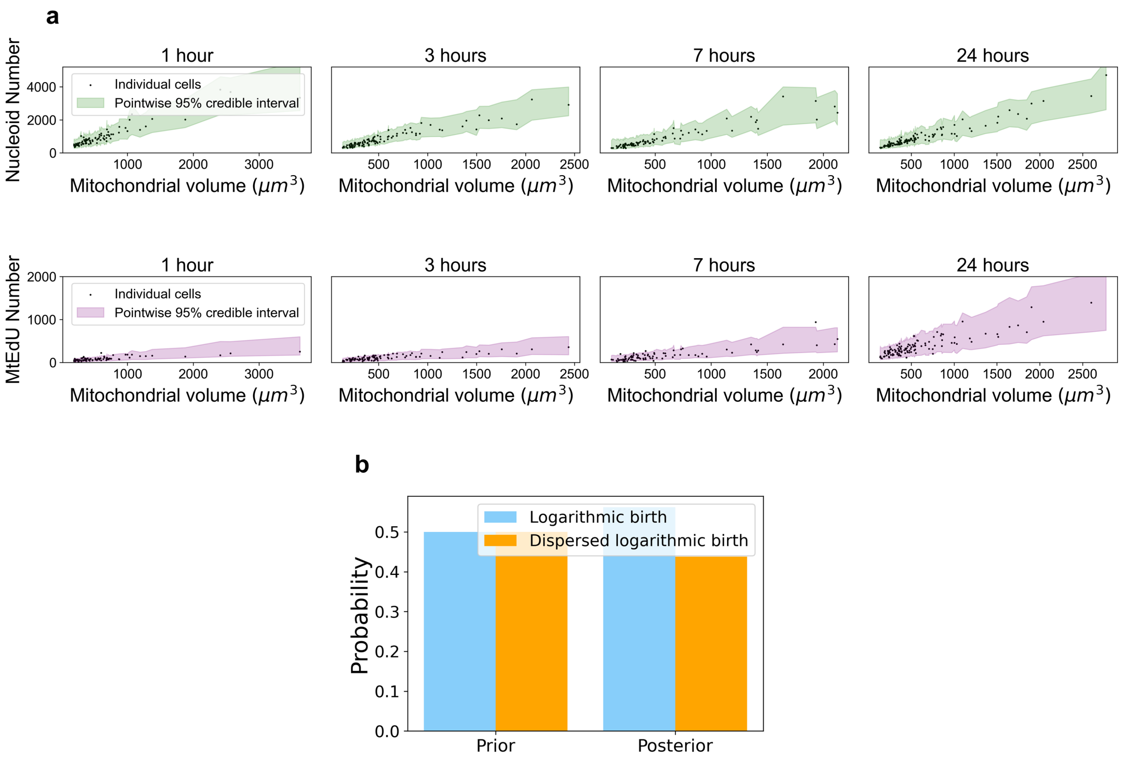

**Supplementary Fig. 21: Dispersed logarithmic control.** **a**, Single cell posterior predictive plots of nucleoid number (top) and mtEdU number (bottom). We see that 93% of mtEdU data points lie within the 95% credible intervals, as opposed to 78% for the ordinary three-population model (Supplementary Fig. 6b), solving the overdispersion issues. **c**, ABC model selection between logarithmic birth and dispersed logarithmic birth.

**a**

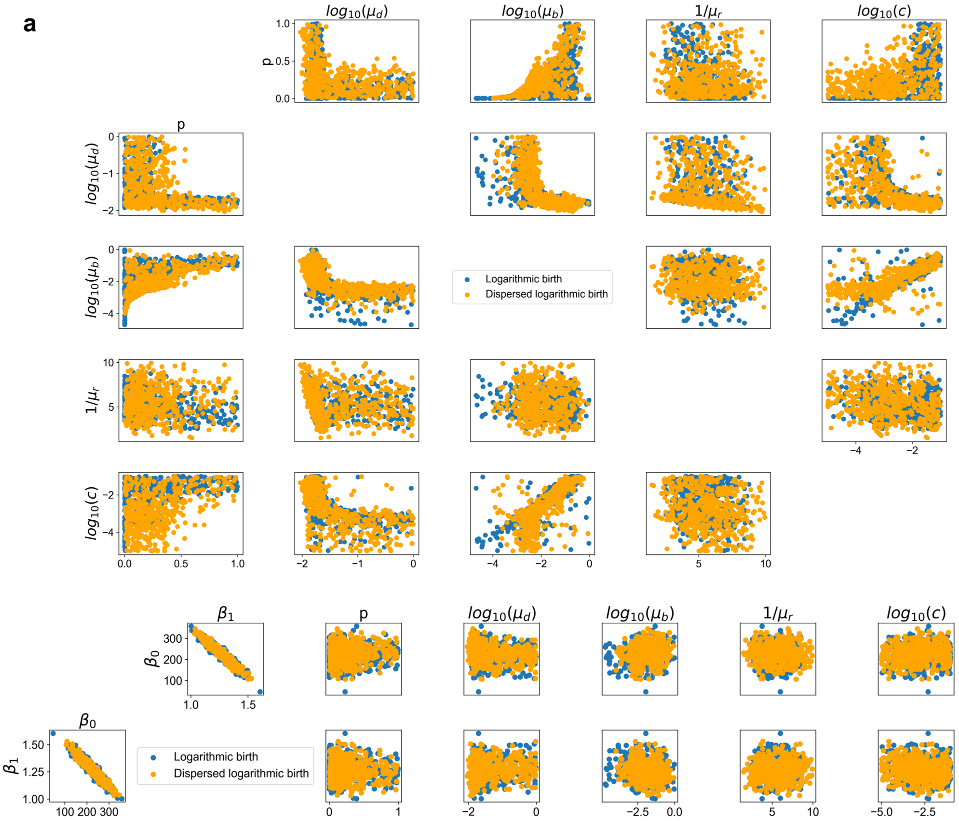

**b**

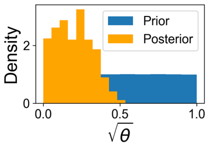

**Supplementary Fig. 22: Posteriors of logarithmic birth and dispersed logarithmic birth**. **a**, Showing that other than $c$, posteriors are indistinguishable between logarithmic and dispersed logarithmic birth. **b**, $\theta$ posterior for dispersed logarithmic birth.

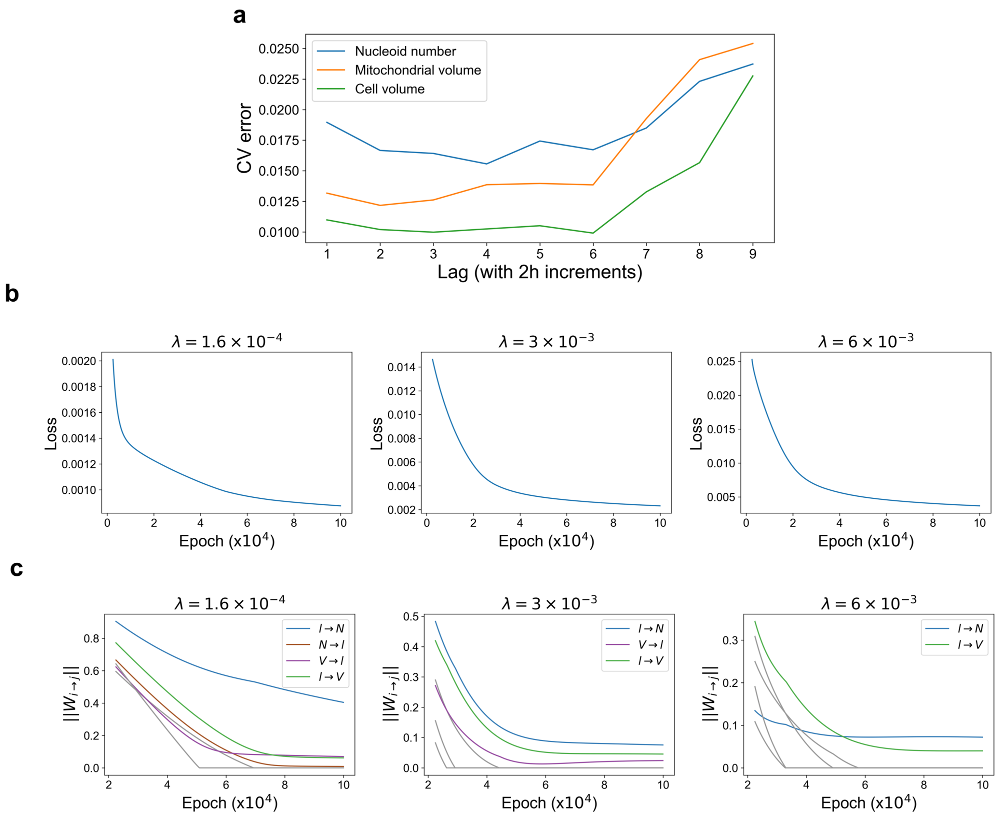

**Supplementary Fig. 23: Linear Granger causality cross validation, and neural Granger causality training**. **a**, Cross validation error for linear Granger causality when predicting nucleoid number (blue, optimal lag 4), mitochondrial volume (orange, optimal lag 2) and cell volume (green, optimal lag 6), for different choices of lag parameter. **b**, Training loss for neural Granger causality during training for three different regularisation parameters. **c**, Parameter trajectories for neural Granger causality during training for three different regularisation parameters.

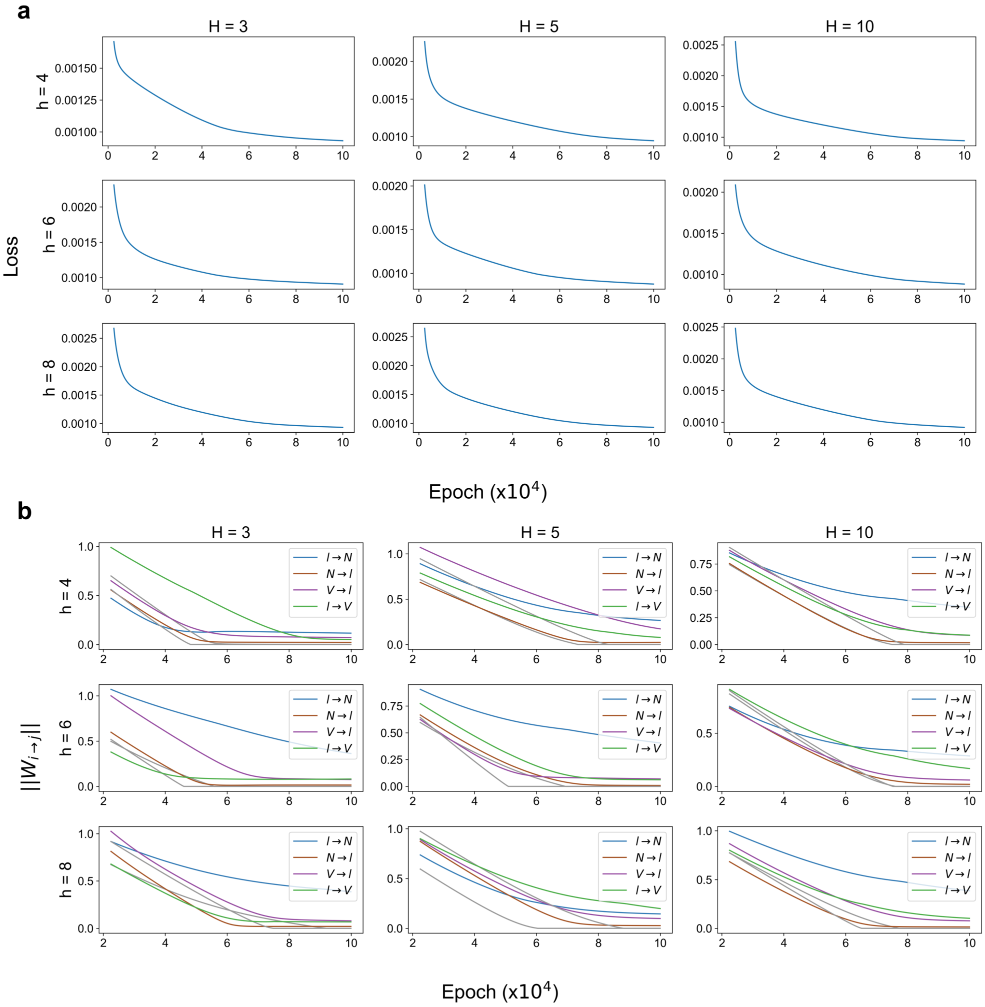

**Supplementary Fig. 24: Neural Granger causality results are robust to different choices of context length** ($h$) **and hidden layer size** ($H$). **a**, Training loss over time for 9 models of different context lengths (rows) and hidden layer sizes (columns). **b**, Parameter trajectories during training for each of the 9 models in (a). Every model selects the same four nonzero parameters when the regularisation parameter is tuned to select the four strongest causal connections.

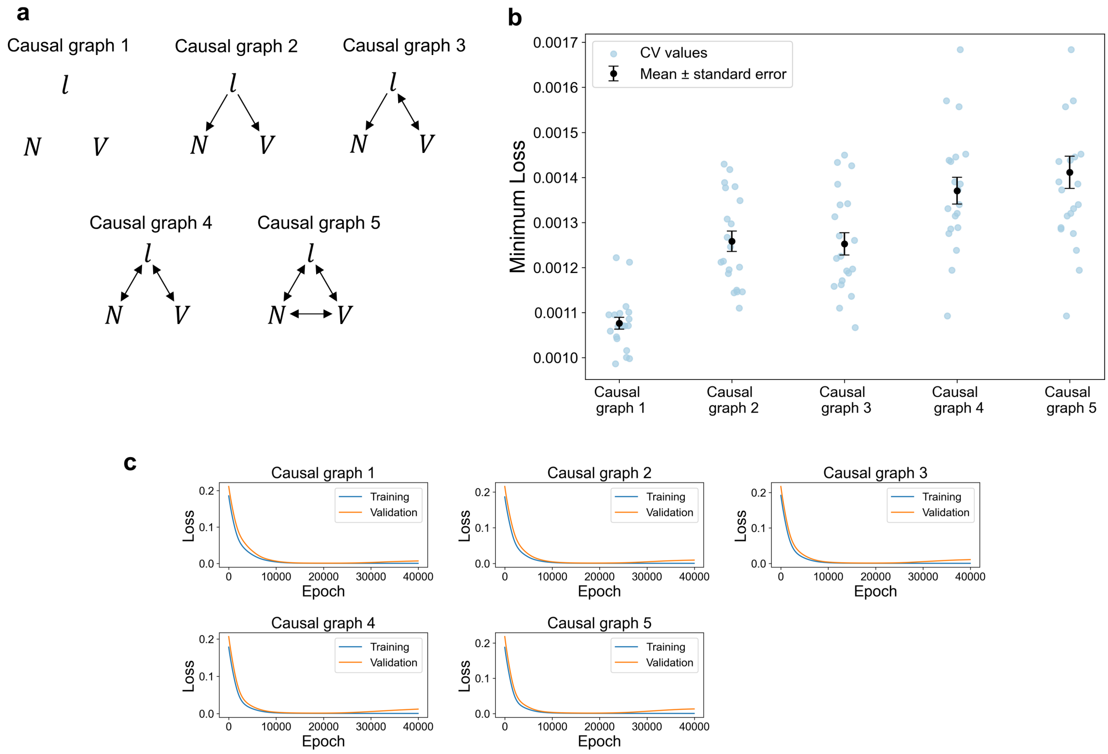

**Supplementary Fig. 25: Neural Granger causality cross validation. a**, Labels of the five causal graphs used in the cross validation. **b**, Minimum cross validation loss recorded for 20 iterations, for each causal graph, and their means. **c**, Mean over 20 iterations of the training and cross validation loss for all causal graphs.
